# Molecular basis for positional memory and its reprogrammability in limb regeneration

**DOI:** 10.1101/2023.10.27.564423

**Authors:** L Otsuki, SA Plattner, Y Taniguchi-Sugiura, EM Tanaka

## Abstract

Upon limb amputation in salamanders, anterior and posterior connective tissue cells form distinct signalling centres that together fuel successful regeneration. The molecular properties that distinguish anterior and posterior cells prior to injury, which enable them to initiate different signalling centres after amputation, are not known. These anterior and posterior identities, crucial for regeneration, were thought to be established during development and to persist through successive regeneration cycles as positional memory. However, the molecular nature of these memory states and whether these identities can be engineered have remained outstanding questions. Here, we identify a positive feedback mechanism encoding posterior identity in the axolotl limb, which can be used to newly encode positional memory in regenerative cells. Posterior cells express residual levels of the bHLH transcription factor *Hand2* from development and this is a priming molecule necessary and sufficient to establish a *Shh* signalling centre after limb amputation. During regeneration, *Shh* feeds back and reinforces *Hand2* expression in nearby cells. *Hand2* is sustained following regeneration, safeguarding posterior memory, while *Shh* is shut off. As a consequence of this *Hand2-Shh* system, anterior and posterior identities are differentially susceptible to alteration. Posterior cells are stabilised against anteriorisation as their expression of *Hand2* poises them to trigger the *Hand2-Shh* loop. In contrast, anterior cells can be reprogrammed: a transient exposure to *Shh* during regeneration causes anterior cells to gain *Hand2* expression and a lasting competence to express *Shh*. In this way, regeneration is an opportunity and entry point to re-write positional memory. Our results implicate positive feedback in the stability of positional memory and explain why positional memory is more easily altered in one direction (anterior to posterior) than the other. Because modifying positional memory changes signalling outputs from regenerative cells, our findings have wider implications for tissue engineering.

## Main

Many adult cells retain positional information, as evidenced by spatially organised differences in gene expression and chromatin that appear to be inherited from embryonic development ^1,2^. As such information is salient to restore tissue after injury, an exciting prospect would be to harness this patterning information to engineer functional tissues for regenerative purposes. An important example illustrating this potential is the salamander limb, which is capable of regenerating throughout life ^3^. Upon injury or surgery that changes positional information in one of the three major limb axes, proximal-distal (‘upper arm to hand’), anterior-posterior (‘thumb to little finger’) or dorsal-ventral (‘back of hand to palm’), salamander limb cells induce regenerative outgrowths. Among the salamander limb cells, connective tissue cells are known to encode positional information ^4,5^, and any positional discontinuities in the limb are likely recognised at both gene expression and chromatin levels. For example, proximal and distal connective tissue cells express different levels of the cell surface molecules *Prod1* and *Tig1* and harbour differential epigenetic marking on *Hox* genes and other patterning gene loci ^6–8^. Similarly, dorsal, but not ventral, connective tissue cells transcribe the transcription factor *Lmx1b* ^9,10^. It is generally assumed that these features allow regenerative cells to sense and organise themselves correctly in space, and to launch appropriate patterning gene programmes. However, it has been difficult to assay such functions due to a paucity of molecular readouts and no manipulation of these genes has to date resulted in the expected changes to limb morphology.

Among the limb axes, the anterior-posterior axis plays a particularly important role in launching and sustaining regeneration. After a limb amputation in axolotls, *Fgf8* secreted from anterior blastema cells interacts with *Shh* secreted from posterior blastema cells in an evolutionarily conserved positive feedback loop to induce limb outgrowth ^11–13^. As a consequence, perturbing the anterior-posterior system results in predictable alterations to limb morphology. For instance, when assembling anterior-only or posterior-only limbs through surgery, these fail to regenerate properly after amputation ^14,15^. Conversely, it is possible to generate an accessory (extra) limb by transplanting posterior limb skin to an innervated anterior wound (or *vice versa*) in a defined assay that allows study of conditions that induce or block anterior-posterior discontinuity (ALM, accessory limb model) ^16^. Interestingly, *Fgf* ligands in other vertebrates are expressed in the distal, apical ectodermal ridge (AER) of the limb bud rather than in anterior connective tissue cells as in salamanders ^13,17–19^. Thus, the critical role of anterior-posterior interactions in regeneration arose due to spatial re-wiring of *Fgf/Shh* in salamanders. Importantly, *Fgf8* and *Shh* are not expressed in the uninjured salamander limb meaning that, despite almost 50 years passing since the establishment of the above-mentioned molecular-morphogenetic assays, the source of anterior-posterior identity is not known. This source has been referred to as positional memory in view of its stability and role in instructing positional information over successive regeneration cycles ^3^. Although the ability to alter positional memory would have implications for regenerative engineering, progress thus far has been hindered by an incomplete understanding of the underlying mechanisms.

Here, and to identify how positional identity is encoded in limb cells, we tracked living cells in the developing and injured axolotl using fluorescent reporters, lineage tracing and genetic/pharmacological perturbations. We found that, after embryonic development, posterior cells retain residual levels of *Hand2*, while shutting down *Shh*. Residual *Hand2* acts as a posterior memory by priming cells to re-express *Shh* following an amputation. Reciprocally, amputation-induced *Shh* induces *Hand2* in nearby cells, re-affirming posterior memory in the regenerated limb before being shut down again. In this way, a positive feedback loop between *Hand2* and *Shh* stably maintains posterior positional identity through successive regeneration cycles. We found that the positional identity of anterior cells is transiently reprogrammable during regeneration. By inducing the *Hand2-Shh* loop in regenerating anterior cells, we overwrote them with a posterior memory, enabling them to subsequently express *Shh.* Our results pinpoint how positive feedback instructs acquisition of stable positional memory and shed light on the rules governing malleability of positional memory in regenerative cells.

### Posterior information is not limited to the *Shh* cell lineage

We first sought to identify the posterior cells that give rise to a *Shh* signalling centre during regeneration. Given that the expression domains of *Shh* and *Fgf8* during limb regeneration resemble those in the developing embryonic limb (**Fig. 1a-b**) ^13,18–20^, we hypothesised that embryonic *Shh* cells are retained in the fully formed limb, and that these cells encode the posterior information required to express regenerative *Shh*. To test this hypothesis, we established a transgenic axolotl model in which embryonic *Shh* cells and regenerative *Shh* cells are labelled with different fluorescent proteins in the same animal. In previous work, the evolutionarily conserved enhancer sequence ZRS (ZPA regulatory sequence, also known as MFCS1) was demonstrated to be both necessary and sufficient to direct *Shh* expression in vertebrate limb buds ^21,22^. Thus, we identified the axolotl ZRS and used this sequence to drive co-expression of TFP (teal fluorescent protein) and a tamoxifen-inducible Cre recombinase in transgenic axolotls. We designate these as ZRS>TFP axolotls (**Fig. 1c**) and summarise the design and expression domains of all transgenic axolotls generated in this study in **Extended Data Fig. 1a-e**. Having confirmed that TFP labelled *Shh* cells during limb development and regeneration (**Extended Data Fig. 2a-c)**, we performed lineage tracing of embryonic *Shh* cells by crossing the ZRS axolotl with our previously published loxP-mCherry fate mapping axolotl (**Fig. 1c**) ^8^. In this model, we expected limb cells derived from embryonic *Shh* lineage cells to express mCherry and cells switching on *Shh* after amputation to express TFP. When we performed a transient treatment with 4-OHT (4-hydroxytamoxifen) at limb bud stage 42, most embryonic *Shh* cells were indeed permanently labelled with mCherry, allowing us to track their fates in the mature limb (labelling efficiency 72.7 ± 18.3 %, *n* = 9 limbs, methodology detailed in **Extended Data Fig. 2d**).

**Fig. 1:**
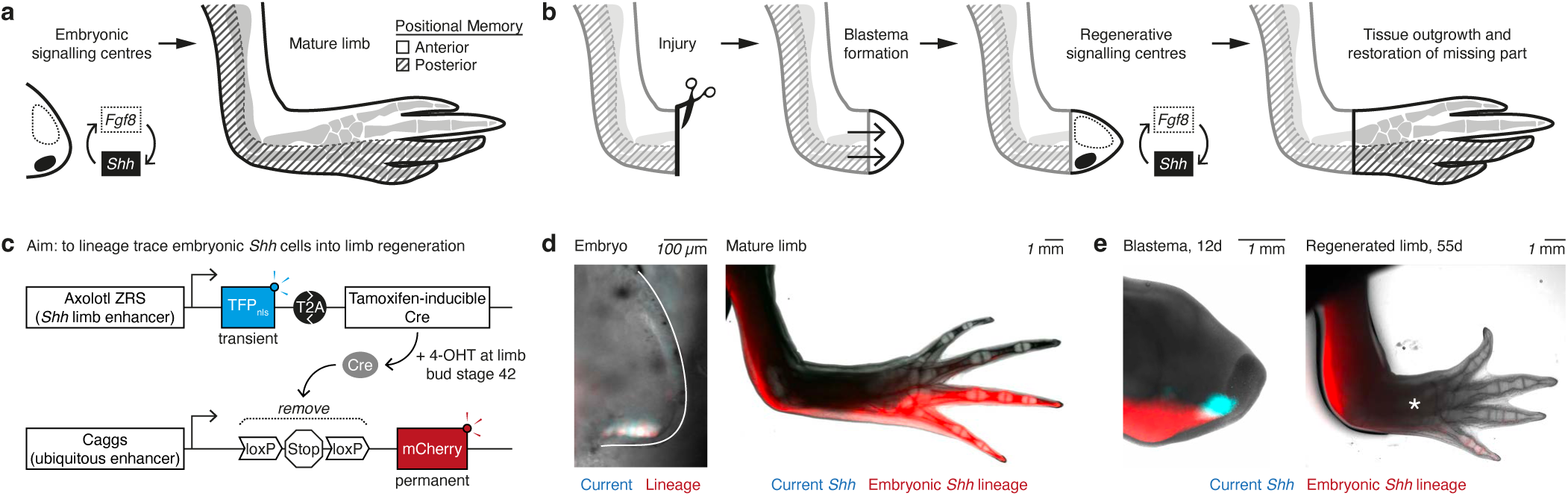
Regenerative *Shh* cells can arise from outside of the embryonic *Shh* lineage. **a,** Schematic of limb development. An Fgf8 signalling centre (anterior) and a Shh signalling centre (posterior) establish the anterior-posterior axis of the limb. **b,** Schematic of limb regeneration. A proliferative cell mass (blastema) drives regeneration from the amputation plane. As during development, Fgf8 and Shh signalling centres drive limb outgrowth and patterning. **c,** Genetic strategy to label embryonic Shh lineage cells and regenerative Shh cells. The ZRS enhancer drives co-expression in embryonic Shh cells of nuclear-localised TFP and tamoxifen-inducible Cre (ERT2-Cre-ERT2), which are co-translationally separated at the intervening T2A sequence. Treatment with 4-OHT induces ERT2-Cre-ERT2 to translocate to the nucleus, resulting in recombination and removal of the loxP-Stop-loxP cassette and permanent labelling of embryonic Shh cells with mCherry. The ZRS enhancer is not expressed in the mature limb, but re-expresses in Shh cells after amputation, enabling comparison between embryonic Shh cells (mCherry, red) and regenerative Shh cells (TFP, cyan) in the blastema. **d,** Left: Stage 44 ZRS>TFP limb bud labelled through the procedure depicted in (c). Current Shh expression (cyan) and Shh lineage label (red) overlap. Right: Mature ZRS>TFP limb (11 cm axolotl). Shh is not expressed (no cyan), but the contributions of embryonic Shh cells to the posterior limb are visible (red). **e,** Left: The same limb as in (d), imaged at 12 days post-amputation. Many regenerative Shh cells (cyan) are mCherry-negative. 23.1 ± 22.1% of TFP signal overlapped mCherry signal in widefield images (n = 10). Right: The fully regenerated limb is depleted for embryonic Shh cells (red) in the regenerated part (asterisked). Images were acquired with a widefield microscope from the ventral side for limb buds and from the dorsal side post-embryonically.

We found that embryonic *Shh* cells contributed to all limb segments in axolotls: the posterior ∼20 % of the upper arm, the posterior ∼20 % of the lower arm and the posterior 1.5 digits of the hand (**Fig. 1d**, **Extended Data Fig. 3a**). Pulsing with 4-OHT later during limb development labelled the most distal subset of cells in the lower arm and hand (**Extended Data Fig. 3b**). To test if the embryonic *Shh* lineage-derived cells gave rise to the regenerative *Shh* cells, we amputated forelimbs through the lower arm and tracked blastema outgrowth. Although embryonic *Shh* cells (mCherry-positive) contributed to the posterior blastema, the vast majority of regenerative *Shh* cells (TFP-positive) were mCherry-negative (**Fig. 1e**). This striking difference suggests that regenerative *Shh* cells primarily arise *de novo* and not through the embryonic *Shh* lineage. Consistent with this interpretation, we found that embryonic *Shh* cells were depleted from the regenerated part of the limb (**Fig. 1e**, **Extended Data Fig. 3c**).

Next, we sought to remove the source of embryonic *Shh* by surgically removing mCherry-positive embryonic *Shh* lineage cells prior to amputation (**Extended Data Fig. 3d-e**). Overall, we achieved 88.7 ± 6.1 % depletion of mCherry-positive cells from the 9 days post-amputation (dpa) blastema, the time point at which regenerative *Shh* is clearly induced in control animals (*n* = 6 limbs). When we conducted lineage tracing after amputations, we found that animals lacking embryonic *Shh* cells retained capacity for inducing TFP and fully regenerated limbs with comparable timing to control animals (**Extended Data Fig. 3f-g**). Taken together, these data confirm that embryonic *Shh* cells are dispensable for generating a regenerative *Shh* signalling centre after an amputation. We conclude that the information required to express *Shh* during posterior limb regeneration is not restricted to the embryonic *Shh* lineage but that, instead, a wider domain of posterior cells in the mature limb is competent to express *Shh*.

### Anterior and posterior cells are primed by locally expressed transcription factors

Given the widespread competence for *Shh* expression, we next compared the transcriptomes of connective tissue cells isolated from the anterior and posterior halves of axolotl limbs to identify posterior identity genes (**Fig. 2a**). To this end, we purified transgenically labelled *Prrx1*+ dermal connective tissue cells, which are known to be strong carriers of positional memory (as previously demonstrated by surgical assays including ALM) and are expected to be less heterogeneous than interstitial (deeper) connective tissue lineages ^16,23,24^.

**Fig. 2:**
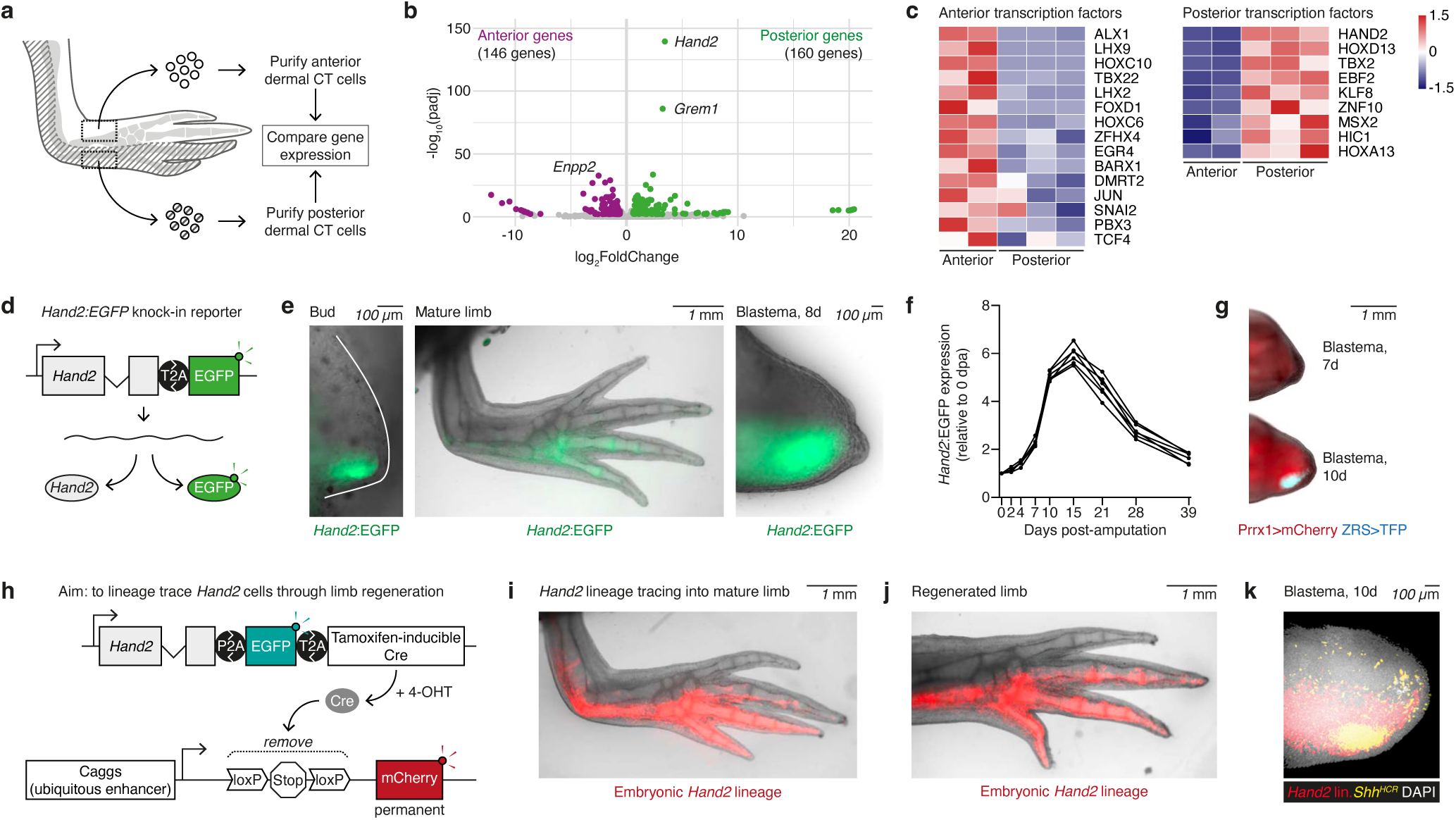
Posterior memory cells increase *Hand2* expression, then give rise to regenerative *Shh* cells. **a,** Experimental strategy to identify anterior and posterior identity genes. Anterior or posterior skin was harvested from lower forelimbs in which connective tissue cells had been genetically labelled with mCherry (see Methods). Dermal connective tissue cells were purified from these skin samples by FACS, then processed for 3’ RNA-sequencing (QuantSeq, Lexogen). **b,** Volcano plot depicting genes differentially expressed in anterior dermal cells (magenta) or posterior dermal cells (green) (DESeq2 analysis). The most statistically significant anterior gene (Enpp2) and posterior gene (Hand2), as well as the second most significant posterior gene (Grem1) are indicated. **c,** Heatmaps depicting the relative expression of transcription factors differentially expressed in anterior and posterior cells. Genes are ordered by decreasing significance. Heatmap scores are normalised by row. **d,** Hand2:EGFP knock-in axolotl, in which the last exon of endogenous Hand2 was replaced by a knock-in cassette encoding Hand2 last exon-T2A-EGFP. **e,** Hand2:EGFP expression (green) in limb bud, mature limb and 8 dpa blastema. We confirmed expression in mature limbs in axolotls up to 15 cm size (sexually mature). **f,** Mean Hand2:EGFP fluorescence in regenerating limbs. n = 6. Each line connects longitudinal data from 1 limb. **g,** Shh expression during regeneration, assessed using double transgenic axolotls that express Prrx1>mCherry (red, connective tissue cells) and ZRS>TFP (cyan, regenerative Shh cells). 1 out of 7 axolotls expressed ZRS>TFP at 7 dpa. 7 out of 7 axolotls expressed ZRS>TFP at 10 dpa. **h,** Lineage tracing of Hand2 cells, using a comparable strategy to Fig. 1c. **i,** Mature limb labelled for embryonic Hand2 lineage cells (red). **j,** A regenerated limb from the same experimental cohort as in (i), at 55 dpa. The Hand2 lineage regenerates a similar spatial domain as it contributed to originally. **k,** Wholemount image of a Hand2 lineage-traced blastema at 10 dpa, stained for Shh mRNA (yellow) and DAPI (white). N = 4 blastemas.

We found ∼300 differentially expressed genes between anterior and posterior dermal cells (DESeq2, *α* < 0.01) (**Fig. 2b**). Of these, *Hand2* expression dominated the posterior cell signature as ordered by stastistical significance (**Fig. 2b**, **Extended Data Table 1**). *Hand2* encodes a bHLH transcription factor that is expressed posteriorly in vertebrate limb buds and has an evolutionarily conserved function in inducing *Shh* in mouse (*Mus musculus*), zebrafish (*Danio rerio*) and chick (*Gallus gallus*) embryos ^25–27^. Although *Hand2* has not been implicated in post-embryonic positional memory, this gene would fit with a previous assertion that the source of posterior information is located intracellularly, based on enzymatic removal of cell surface molecules ^28^. Scrutiny of other top differentially expressed genes, ordered by *p*-value or by fold-change (**Extended Data Tables 1-4**), revealed that several transcription factors with an anterior-posterior expression pattern in mouse limb buds were expressed in the corresponding domain of the mature axolotl limb. For example, posterior cells expressed *Hoxd13* and *Tbx2* and anterior cells expressed *Alx1, Lhx2* and *Lhx9* (**Fig. 2c**, **Extended Data Fig. 4a-c**). This contrasted with general forelimb connective tissue genes like *Tbx5* and *Prrx1*, which were expressed in both anterior and posterior cells (**Extended Data Fig. 4d**). Thus, dermal cells in the mature axolotl continuously express transcription factors in spatial domains resembling those in the embryo. A similar phenomenon was first reported in regenerative zebrafish pectoral fins ^29^.

Gene Ontology (GO) analysis revealed that differentially expressed genes were enriched in extracellular matrix and cell adhesion terms (**Extended Data Fig. 4e-h**, **Extended Data Tables 5-6**). These molecules, including collagen subtypes, might contribute towards distinct extracellular signalling environments in the anterior and posterior of the limb. This led us to survey signalling pathways known to be important during limb development. We found that *Shh*, *Fgf* and *Wnt* pathway components were either not detected, or were equivalently expressed between anterior and posterior cells in the mature limb (**Extended Data Fig. 4i-k**). We only observed differential anterior-posterior expression in a few genes belonging to the BMP and RA (Retinoic acid) signalling pathways (**Extended Data Fig. 4l-m**). Of these, the secreted BMP antagonist *Grem1*, reported to act as a relay intermediate between *Shh* cells and *Fgf* cells during mouse limb development and to potentially play a similar role in salamanders ^13,30,31^, was enriched posteriorly (**Fig. 2b**). For the RA pathway, the aldehyde dehydrogenase enzyme *Aldh1a3* was enriched anteriorly whereas its responder gene, encoding for the retinoic acid receptor responder 1 (*Rarres1*, also known as *Tig1*), was enriched posteriorly. *Tig1/Rarres1* was recently implicated in proximal-distal positional identity through misexpression assays and effects on genes known to be involved in proximal-distal patterning ^7^. However, the loss of function phenotype of *Tig1/Rarres1 in vivo* is not known, nor is it known if this gene can reprogramme positional memory for subsequent regeneration.

### *Hand2* cells are the source of regenerative *Shh* cells

We found that *Hand2* is the gene with the most significant differential expression in the anterior-posterior axis of the mature limb (**Fig. 2b**). Given that *Hand2* in mouse limb buds is necessary and sufficient for *Shh* expression and directly binds to the ZRS enhancer of the *Shh* gene ^27,32,33^, we hypothesised that steady-state *Hand2* primes axolotl cells to express *Shh* during regeneration.

To better understand the dynamics of *Hand2* expression, we generated a *Hand2:*EGFP knock-in reporter in which endogenous *Hand2* is co-expressed with EGFP (enhanced green fluorescent protein) (**Fig. 2d**). We detected *Hand2*:EGFP in the posterior limb bud, in the posterior steady-state limb and in the posterior blastema following amputation (**Fig. 2e**). Thus, posterior connective tissue cells continuously express *Hand2* from embryonic development. In steady-state limbs, we detected *Hand2*:EGFP in both dermal cells and interstitial connective tissue cells (**Extended Data Fig. 5a**). In all contexts, we expect *Hand2* protein (and not only mRNA) to be present, as EGFP production requires translation of upstream *Hand2* protein and ribosome ‘skipping’ in the T2A sequence ^34,35^. After limb amputation, the mean *Hand2:*EGFP fluorescence increased (5.9 ± 0.4 fold) in the regenerating blastema between 0 and 15 dpa, before returning to steady-state levels (**Fig. 2f**). When quantifying individual cells by flow cytometry, we found a similar, five-fold increase in mean *Hand2*:EGFP fluorescence per cell between 0 and 14 dpa (**Extended Data Fig. 5b**). The rise in *Hand2* expression begins before *Shh* is induced: *Hand2:*EGFP fluorescence increased more than two-fold (2.3 ± 0.2) prior to the onset of ZRS>TFP expression at 7 dpa (**Fig. 2f-g**).

Next, we assessed if *Hand2* cells give rise to regenerative *Shh* cells. Because *Hand2*-expressing cells have not previously been lineage-traced in the limb in any organism, we generated a *Hand2* knock-in axolotl and performed lineage tracing using the same strategy as for the ZRS>TFP axolotl (**Fig. 2h**, **Extended Data Fig. 5c**). We found that embryonic *Hand2* cells contribute to the posterior half of the axolotl upper arm and lower arm, and to the posterior 2.5 digits of the hand, which resembles the active *Hand2* expression domain (**Fig. 2e,i**). After amputation, *Hand2*-lineage cells regenerated a comparable spatial domain (**Fig. 2j**, **Extended Data Fig. 5d-e**). Thus, unlike the embryonic *Shh* lineage, the *Hand2* lineage is maintained in the posterior limb following regeneration. When we labelled *Hand2* cells at steady state, we obtained qualitatively similar results: they were restricted to the posterior half of the limb. We note however, that a higher 4-OHT dose and multiple treatments were required to label steady-state *Hand2* cells – likely due to weaker *Hand2* expression (and consequently Cre expression) than during development/regeneration (**Extended Data Fig. 5f-g**). Importantly, and as validated by *in situ* hybridisation and 3D imaging, *Hand2* cells give rise to the entire *Shh* cell population during regeneration (**Fig. 2k**, **Extended Data Fig. 5h**). Taken together, we show that the posterior *Hand2* domain is the source of *Shh* cells during regeneration and that this cell population increases *Hand2* expression prior to *Shh* induction.

### *Hand2* encodes posterior identity, but cells lacking *Hand2* do not default to an anterior identity

In mouse and zebrafish embryos, *Hand2* is necessary to express *Shh* in the limb/fin bud ^25,27^, and we therefore sought to assess if this embryonic function is conserved in axolotls. To this end, we co-injected fertilised eggs with Cas9 protein and two efficient sgRNAs targeting the *Hand2* translational start codon to generate axolotls mutant for *Hand2* (**Fig. 3a**, **Extended Data Fig. 6a-b**). We elected to conduct analysis of mosaic mutations given that homozygous *Hand2^-/-^* mutant mice die during embryonic development due to heart defects ^36^. In mosaically mutated F0 axolotls (which we refer to as *Hand2* CRISPants) we found that more *Hand2* CRISPant axolotls died before digit patterning relative to control axolotls (52% [*n =* 60/116 injected eggs] vs 14% [*n* = 4/28]), likely due to a high homozygous knockout rate (**Fig. 3b**). When we characterised the limb phenotypes of ‘escaper’ CRISPants hypomorphic for *Hand2* (**Fig. 3b**), we found that roughly half had defects in digit number or outgrowth (45% of *Hand2* CRISPant limbs [*n* = 50/112 limbs]), with the most severe defects represented by no limb outgrowth beyond the shoulder girdle (**Fig. 3c**, *n* = 9). This phenotype is reminiscent of that of *Shh* CRISPant axolotls^37^. We obtained a range of digit number phenotypes, as expected from mosaic inactivation (**Extended Data Fig. 6c-d**). When amputated, almost all *Hand2* CRISPant limbs regenerated fewer digits than they had originally, including those that originally harboured the correct number of digits (**Fig. 3d**, **Extended Data Fig. 6e-f**). Taken together, these data support that *Hand2* is required in limb regeneration.

**Fig. 3:**
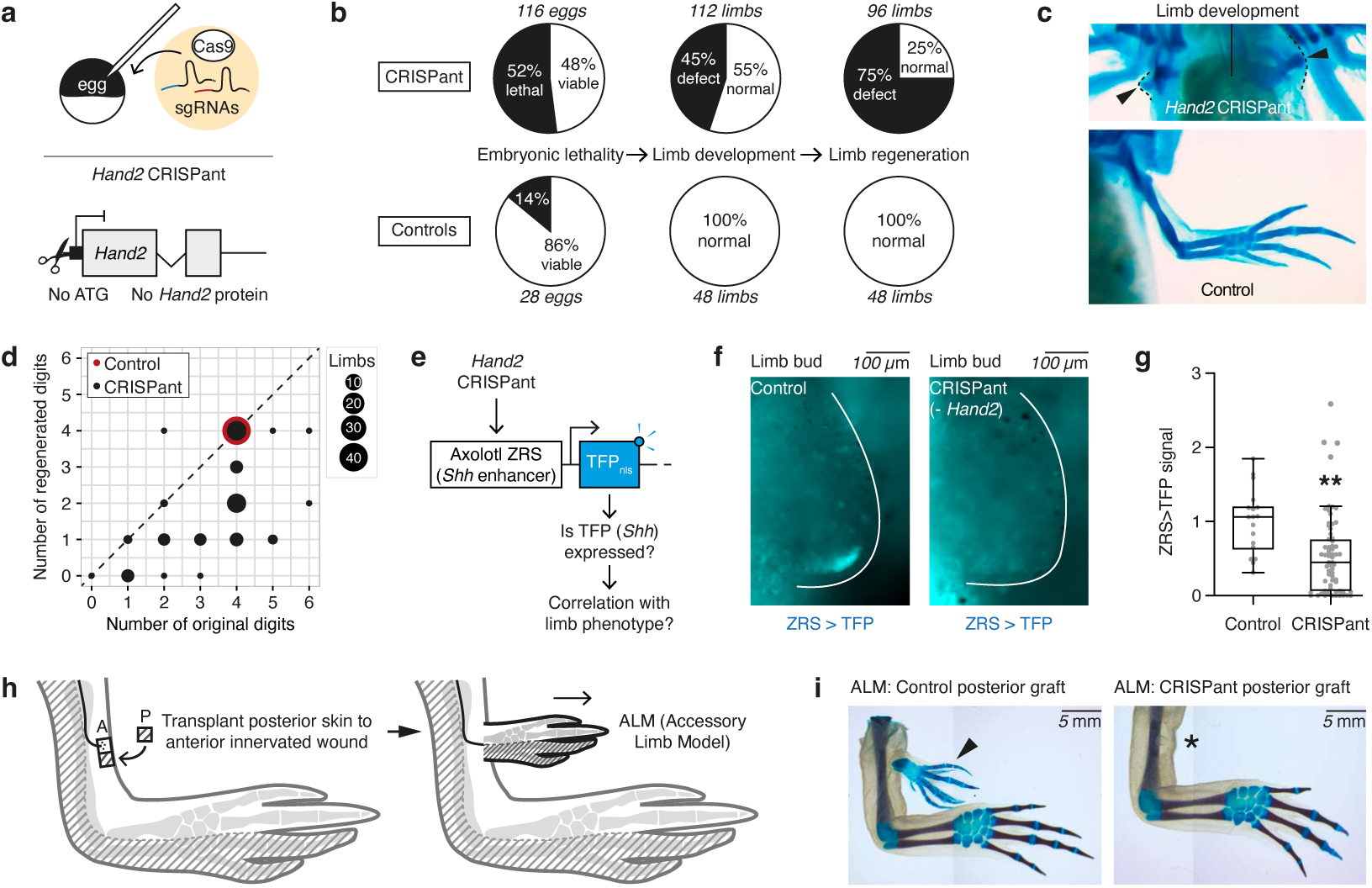
*Hand2* is necessary for *Shh* expression and posterior limb identity. **a,** Experimental strategy to mutate Hand2 in axolotl embryos. CRISPR/Cas9-mediated cutting removes the translational start from Hand2. **b,** Pie charts comparing selected phenotypes of Hand2 CRISPant axolotls (top) and control axolotls (bottom). Control axolotls were injected with Cas9 but not sgRNA. Some Hand2 CRISPant axolotls died between limb development and the regeneration assay. **c,** Alcian blue (cartilage) staining depicting a severe Hand2 CRISPant phenotype in which no limb skeleton forms beyond the shoulder girdle. n = 9 out of 112. **d,** Dot plot comparing the number of digits in Hand2 CRISPant limbs at the end of development (x-axis) with the number of digits that the same limbs regenerated after amputation (y-axis). The size of the dot corresponds to the number of limbs, with a total of 96 limbs in the experiment. The diagonal line indicates the situation where limbs regenerated the same number of digits as they had originally. Most dots reside below this line, indicating that most Hand2 CRISPant limbs regenerated fewer digits than they had originally. **e,** Experimental strategy to assess if Hand2 is necessary for Shh expression, using ZRS>TFP axolotls. **f,** Hand2 CRISPants have less ZRS>TFP (cyan) than controls at limb bud stage 42. **g,** Quantification of (f). An integrated fluorescence intensity (mean fluorescence x fluorescence intensity) was calculated for each limb bud, then normalised to the mean of the control cohort (set to ‘1’). Each dot represents a measurement from 1 limb bud. **: p = 6.9e-3, Kolmogorov-Smirnov test. n = 68 (CRISPant) or 18 (control). **h,** Schematic depicting the anterior ALM assay. A patch of posterior limb skin transplanted to an innervated anterior wound elicits the outgrowth of an accessory limb. **i,** Anterior ALM assays performed with control posterior skin (left) or Hand2 CRISPant posterior skin (right). 3 out of 6 control grafts elicited ALMs (arrowed). 0 out of 6 Hand2 CRISPant grafts elicited ALMs (asterisked). Limbs were stained with Alcian Blue and Alizarin Red to highlight cartilage and bone respectively.

Next, we asked if *Hand2* is required for *Shh* expression. To this end, we generated *Hand2* CRISPant axolotls in the ZRS>TFP genetic background to read out *Shh* expression. If *Hand2* is necessary for *Shh* expression, then *Hand2* CRISPants should express less TFP than controls (**Fig. 3e**). When we live-imaged TFP during development/regeneration, we found that fewer cells expressed TFP or did so at a lower expression level in *Hand2* CRISPant limb buds relative to control limb buds (**Fig. 3f-g**). The *Hand2* CRISPant limb buds with the lowest TFP signal generated limbs with 0-3 digits (**Extended Data Fig. 6g**). We also found a direct positive correlation between TFP signal during development and regeneration of the same animal (Spearman’s rank test, *r* = 0.74, *p* = 2.40e-3, *n* = 14 limbs) (**Extended Data Fig. 6h**). Thus, *Hand2* is necessary for *Shh* expression during axolotl limb development and regeneration, in a function that appears to be conserved in other vertebrates.

An outstanding question in regenerative biology is what happens to a cell’s positional information if it fails to acquire its normal identity. The fact that developing and regenerating *Hand2* CRISPants appeared to lose posterior identity (*Shh* expression), allowed us to answer this question in axolotls. Thus, we tested if *Hand2* CRISPant cells default to an anterior or other positional identity in the Accessory Limb Model (ALM), a tissue engineering model that allows for examining components required for successful regeneration. In the ALM model, an ectopic limb will grow out from an innervated anterior wound if grafted with a patch of mature posterior skin (or *vice versa*) due to positional discontinuity (**Fig. 3h**) ^16^. Here, we first confirmed that grafts of normal posterior skin induced accessory limbs when transplanted to an anterior wound site (*n* = 3 ALMs from 6 surgeries, **Fig. 3i**). If *Hand2* specifies posterior identity, then posterior skin mutant for *Hand2* should lack the capacity to induce accessory limbs in this assay. Indeed, upon transplantation to the anterior wound site of an unmutated host, posterior skin lacking *Hand2* lost its ability to induce accessory limbs (*Hand2* CRISPant skin, *n* = 0 ALMs from 6 surgeries, **Fig. 3i**). Next, we used ALM to test if *Hand2* loss causes cells to acquire an anterior identity. However, and unlike anterior skin, *Hand2* CRISPant posterior skin did not induce ALMs at an innervated posterior wound (*n* = 0 ALMs from 6 surgeries) (**Extended Data Fig. 6i-j**). Taken together, these results show that *Hand2* is necessary for posterior identity and suggest that cells that fail to acquire a posterior identity do not default to an anterior identity.

### Strong *Hand2* misexpression posteriorizes limb and, despite *Shh* induction, blocks limb outgrowth

Because *Hand2* misexpression can trigger ectopic *Shh* expression and supernumerary digits (polydactyly) in mouse or chick limb/wing buds ^26,27^, we next asked if transgenic expression of *Hand2* is sufficient for *Shh* expression in axolotl. To this end, we injected fertilised eggs with an integrating construct harbouring the mouse *Prrx1* limb enhancer ^38^ that would misexpress axolotl *Hand2* N-terminally tagged with mCherry throughout the limb bud and limb blastema (**Fig. 4a**, **Extended Data Fig. 7a**). In F0 animals, we detected mosaic expression of mCherry-*Hand2* at different expression levels, presumably depending on the number of transgene copies integrated into the genome. We found that *Hand2*-misexpressing limb buds expressed ectopic ZRS>TFP (*n =* 7/9 limb buds carrying the TFP transgene) and developed polydactyly (*n* = 7/16 limbs total) (**Fig. 4b**, **Extended Data Fig. 7b-d**). In two cases, anterior *Hand2* misexpression resulted in a complete ectopic limb growing from the upper arm, a phenotype not reported in other species but consistent with the axolotl’s reported ability to generate accessory limbs at sites of anterior-posterior discontinuity (**Fig. 4c**). Taken together, we conclude that singular misexpression of axolotl *Hand2* is sufficient to trigger *Shh*, polydactyly and, in extreme cases, formation of an ectopic limb. Importantly, we found that these phenotypes occurred in animals expressing higher levels of mCherry-*Hand2*, as determined by mCherry intensity (**Extended Data Fig. 7e**).

**Fig. 4:**
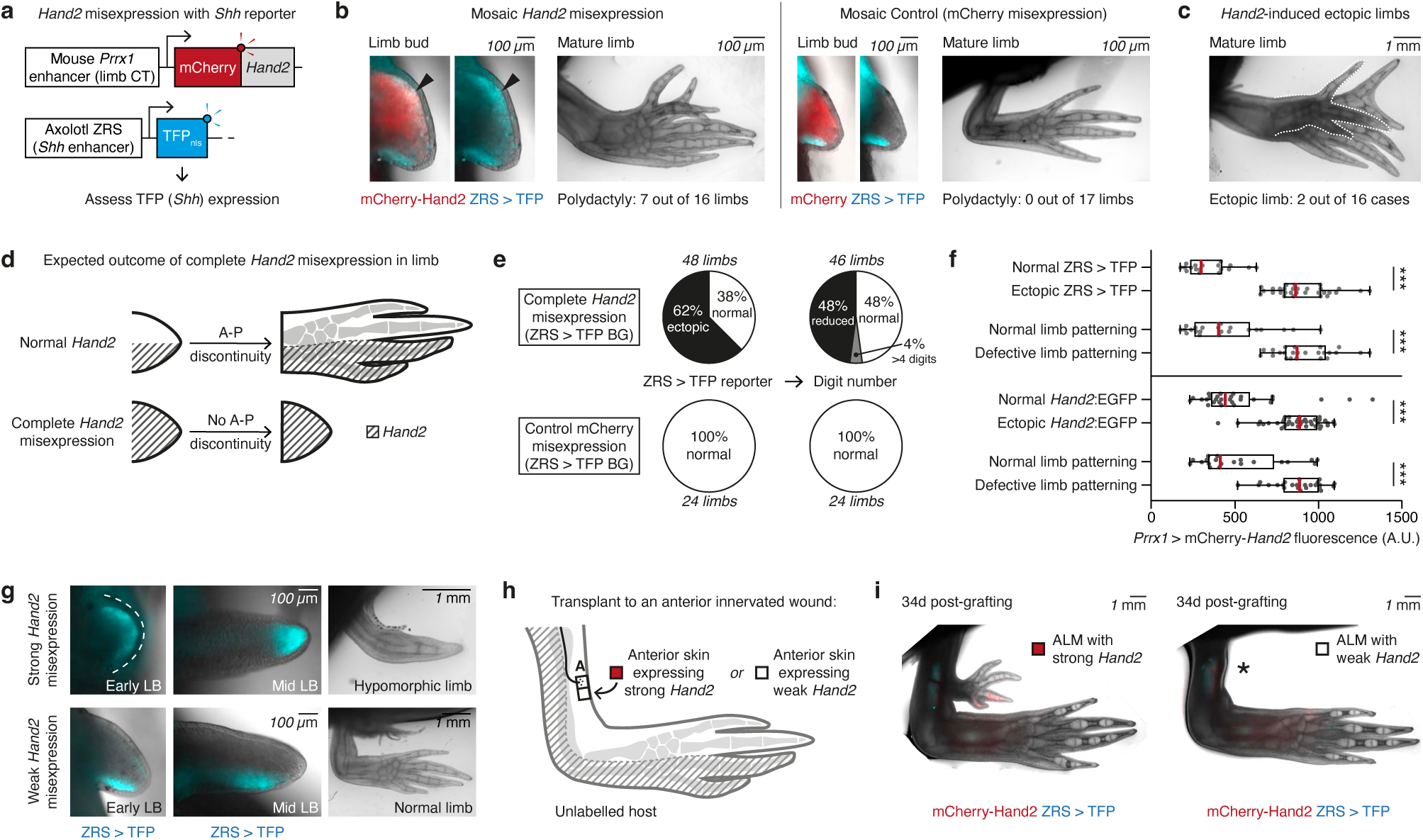
*Hand2* is sufficient for *Shh* expression and posterior limb identity. **a,** Strategy to misexpress mCherry-tagged Hand2 in connective tissue cells, with ZRS>TFP reporter to estimate Shh expression. Control animals misexpressed mCherry-only instead of mCherry-Hand2. **b,** Mosaic mCherry-Hand2 misexpression during limb development induced ectopic anterior ZRS>TFP (cyan, arrowed) and polydactyly. **c,** Mosaic mCherry-Hand2 misexpression induced an ectopic limb (dotted line) in 2 out of 16 cases. **d,** Complete Hand2 misexpression in connective tissue cells should eliminate anterior-posterior discontinuity and limb outgrowth. **e,** Pie charts summarising selected phenotypes arising from complete misexpression of mCherry-Hand2 (top) or mCherry alone (bottom) in the ZRS>TFP background (BG). **f,** Box plots comparing the level of mCherry-Hand2 expression (x-axis) with the resulting limb phenotypes (y-axis). Each dot represents one limb. Data summarised from two experiments - one in the ZRS>TFP genetic background (top, n = 48) and one in the Hand2:EGFP genetic background (bottom, n = 68). From top to bottom, ***: p = 5.07e-8, p = 1.57e-6, p = 1.96e-6, p = 1.60e-4, Kruskal-Wallis tests followed by post-hoc tests. **g,** Comparison of ZRS>TFP expression (cyan) in axolotls misexpressing strong mCherry-Hand2 (top) or weak mCherry-Hand2 (bottom). Strongly misexpressing limbs expressed ZRS>TFP throughout the distal mesenchyme and developed hypomorphic limbs, while weakly misexpressing limbs had posteriorly-restricted TFP, similar to mCherry-only controls. **h,** Anterior ALM to assess the positional identity of cells misexpressing strong Hand2 or weak Hand2. **i,** Anteriorly located skin misexpressing strong Hand2 can induce an accessory limb from an innervated anterior wound (left, n = 8 ALMs from 8 grafts), while a graft misexpressing weak Hand2 cannot (right, n = 0 ALMs from 8 grafts).

In the context of limb outgrowth, the source of positional discontinuity in axolotls for both development and regeneration is the anterior mesenchyme-posterior mesenchyme, whereas in other vertebrates, it is the distal epidermis-posterior mesenchyme ^17^. Potentially, complete *Hand2* misexpression throughout limb bud mesenchyme might eliminate positional discontinuity and limb outgrowth in axolotls (**Fig. 4d**). This prediction contrasts with the polydactyly resulting from mosaic *Hand2* misexpression in axolotl, mouse or chick but would be similar to the phenotype seen after amputation of surgically assembled double posterior salamander limbs ^14,15^.

To test this prediction, we generated germline transmitted F1 axolotls uniformly misexpressing mCherry-*Hand2* in limb connective tissue cells. As expected, inheritance of different numbers of Prrx1>mCherry-*Hand2* transgenes through the germline resulted in different expression levels, which could be distinguished by mCherry fluorescence intensity. To detect posteriorisation, we performed this experiment in the genetic background of ZRS>TFP, which reports *Shh* expression, or *Hand2*:EGFP, which reports endogenous *Hand2* expression. As predicted, *Hand2* misexpression triggered ectopic expression of ZRS>TFP, ectopic *Hand2*:EGFP and generation of hypomorphic limbs in a subset of animals (**Fig. 4e**, **Extended Data Fig. 7f-i**). As in the mosaic experiments, *Hand2* misexpression level was a significant predictor of these phenotypes (**Fig. 4f**). Many axolotls with strong *Hand2* misexpression generated hypomorphic spikes or did not outgrow any limb, concomitant with ZRS>TFP expression extending across the anterior-posterior axis of the outgrowing tip instead of being restricted posteriorly (**Fig. 4g**). *Hand2*:EGFP was expressed uniformly through the limb bud, extending more proximally than ZRS>TFP (**Extended Data Fig. 7j**). Thus, exogenous mCherry-*Hand2* conferred full posterior identity to the limb field and impeded outgrowth. By contrast, we did not find any obvious defects in ZRS>TFP*, Hand2:*EGFP or final limb patterning in sibling axolotls with weak *Hand2* misexpression (**Fig. 4g**, **Extended Data Fig. 7j**). Interestingly, the two-fold mean expression difference between ‘strong’ and ‘weak’ *Hand2* in this experiment (**Fig. 4f**) is similar to the 2.39-fold rise in *Hand2* expression in regenerating posterior cells prior to *Shh* induction (**Fig. 2f**). These data support an unappreciated role for *Hand2* expression levels in inducing *Shh.* Importantly, and as predicted, complete *Hand2* misexpression ablated positional discontinuity.

To demonstrate that *Hand2*-misexpressing limbs were fully posteriorized, we used the ALM assay. As donor grafts, we harvested skin from the anterior side of post-embryonic limbs expressing weak *Hand2* (control) or strong *Hand2* (test). When we grafted the skin patches (which also carried ZRS>TFP in their genetic background) to innervated anterior wounds on unlabelled host animals, we found that all strong *Hand2* skin grafts induced ZRS>TFP expression and anterior ALMs whereas weak *Hand2* skin did neither (*n* = 8 grafts per condition) (**Fig. 4h-i**, **Extended Data Fig. 7k**). Taken together, these results confirm that *Hand2* misexpression during development resulted in posterior identity post-embryonically.

### Anterior-posterior positional memory is reprogrammable during regeneration

Our results thus far supported that *Hand2*, in and of itself, drives acquisition of posterior identity during development and competence to express *Shh* during limb regeneration and generation of accessory limbs. However, whether anterior-posterior identity is fixed after being established during embryonic development remains an open question with wide implications for tissue engineering.

One way to test if anterior-posterior identity is fixed or reprogrammable is to move cells out of their normal location and track changes to their positional identity using molecular criteria. We used transplantation to move labelled anterior or posterior cells to the opposite side of an unlabelled host limb. By transplanting uninjured, mature cells to an uninjured host arm, we could test if mature cells have a fixed or flexible identity. By subsequently amputating the host arm, we could test if positional identity changes in the regenerating blastema. To perform this assay, we generated double reporter axolotls in which anterior and posterior identity can be discriminated through fluorophore expression (**Fig. 5a-b**). As before, *Hand2*:EGFP (green) indicates posterior identity. To report anterior identity, we knocked mCherry into the *Alx4* locus, which labels anterior cells in mice ^39^. *Alx4*:mCherry (magenta) labelled anterior connective tissue cells in axolotl limbs and blastemas, but also labelled a subset of posterior connective tissue cells, explaining why we did not recover *Alx4* as an anterior-specific gene in our transcriptional profiling (**Extended Data Fig. 8a-b**). Nevertheless, this double reporter axolotl is adequate for our assay as it differentially labels anterior identity (mCherry-positive plus EGFP-negative) and posterior identity (EGFP-positive) *in vivo*.

**Fig. 5:**
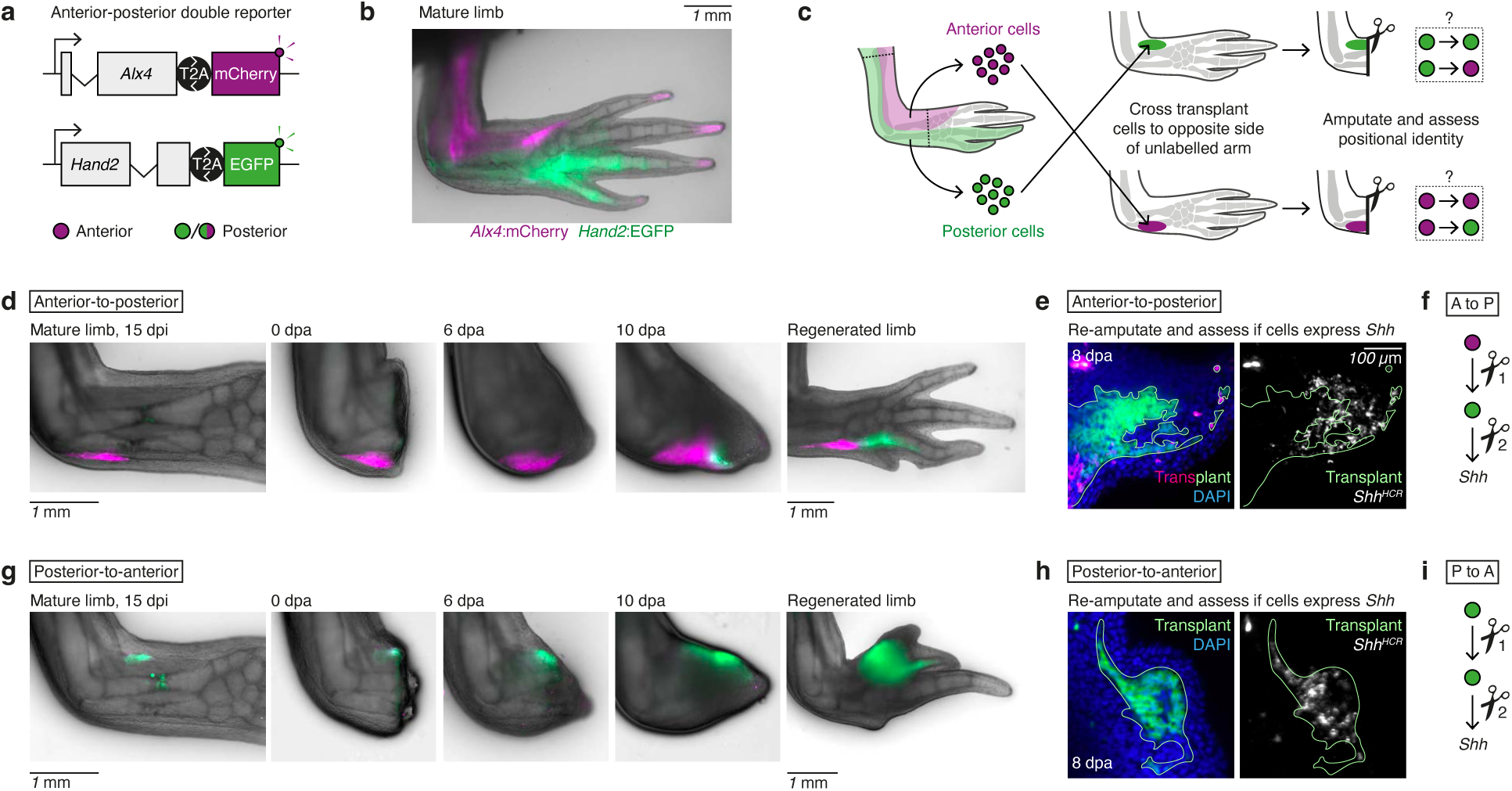
Anterior-posterior positional memory is reprogrammable during regeneration. **a,** Double knock-in axolotl to label anterior cells (magenta only) and posterior cells (green). **b,** Mature limb of an Alx4:mCherry_Hand2:EGFP axolotl. Alx4:mCherry is expressed anteriorly and also in the digit tips. Hand2:EGFP labels posterior cells. **c,** Experimental design to assay if positional identity is fixed. Anterior or posterior connective tissue cells were purified by FACS from Alx4:mCherry_Hand2:EGFP mature limbs, then injected into the opposite side of an unlabelled host limb. After 2 weeks, injected limbs were amputated through the transplant and fluorescence was tracked during regeneration. **d,** Summary of anterior-to-posterior cell injections. mCherry cells remained mCherry+ and EGFP-for 2 weeks after injection. After amputation, cells in the blastema became EGFP+ while cells in the stump remained EGFP-. **e,** Assay to test if newly arising EGFP+ cells can express Shh. Regenerated limbs were re-amputated through the EGFP+ region of the transplant. 8 days later, blastemas were fixed and stained in wholemount for Shh mRNA (white). A subset of EGFP+ transplant cells expressed Shh mRNA. n = 3. **f,** Schematised result of the anterior-to-posterior transplantations. **g,** Summary of posterior-to-anterior cell injections. EGFP cells remained EGFP+ for 2 weeks after injection. After amputation, cells remained EGFP+ in the blastema. These limbs regenerated with aberrant pattern. **h,** Assay to show that EGFP+ cells can still express Shh. Regenerated limbs were re-amputated through the regenerated part of the transplant. 8 days later, blastemas were fixed and stained in wholemount for Shh mRNA (white). EGFP+ transplant cells expressed Shh mRNA. n = 3. **i,** Schematised result of the posterior-to-anterior transplantations.

First, we purified anterior mature limb cells (mCherry+/EGFP-) by fluorescence-activated cell sorting (FACS) and injected these into the posterior forelimb of an unlabelled and uninjured host (**Fig. 5c**). Two weeks after transplantation, the injected anterior cells remained contiguous and mCherry-positive, consistent with a fixed anterior positional identity (**Fig. 5d**). However, we were intrigued to see that amputation revealed a different cellular behaviour. During the first week post-amputation, we found that mCherry-labelled cells invaded the growing blastema. However, from 8 dpa and onward, posterior identity EGFP+ cells started to appear (**Fig. 5d**, **Extended Data Fig. 8c**). Given that the host cells do not carry the *EGFP* transgene, the only source of the EGFP+ cells must be the transplanted anterior cells. As regeneration proceeded, the proportion of EGFP+ cells in the blastema increased and remained the dominant population upon completion of regeneration. At the same time, transplanted cells that had not been recruited into the blastema remained mCherry-positive (**Fig. 5d**). These divergent behaviours (EGFP+ in the regenerated region and mCherry+ in the limb stump) are a visual indication that anterior-posterior identity is fixed at steady-state but flexible in the blastema.

If the anterior cells acquired a stable, posterior identity in this assay, then they should be competent to express *Shh* following a further amputation. Thus, we performed a second amputation through the EGFP+ part of the regenerated limb. Here, we found that the originally anterior cells produced ‘posterior’ descendants expressing EGFP+, and, importantly, that a minority of those descendants expressed *Shh* (**Fig. 5e-f**). Given that this change in positional identity was manifested in the subsequent round of regeneration, we inferred that the positional memory of the anterior cell was stably posteriorized in the transplantation assay.

### Posterior cells retain posterior memory

In the reciprocal experiment, we purified posterior mature connective tissue cells (EGFP+) from double reporter axolotls and injected these into the anterior forelimb of an unlabelled host. Two weeks after injection, the cells remained EGFP+ with no overt increase in mCherry fluorescence (**Fig. 5g**). Thus, like anterior cells, posterior cells have a fixed identity in the unamputated limb. However, unlike anterior cells, we found that transplanted posterior cells retain their identity following amputation. Thus, fluorescent cells remained EGFP+ during regeneration, increasing EGFP expression like normal posterior cells before giving rise to ectopically patterned regenerates (*n* = 3/3 limbs, **Fig. 5g**, **Extended Data Fig. 8d**). These cells retained a posterior memory, as they expressed *Shh* following a second amputation (**Fig. 5h-i**). We cannot exclude that some transplanted cells lost EGFP fluorescence during the experiment and became untraceable; however, such cells did not acquire an overt anterior identity (mCherry expression) (**Extended Data Fig. 8d**). Taken together, these results indicate that anterior blastema cells readily acquire a posterior memory, while posterior blastema cells retained their posterior memory in this transplantation assay.

### Endogenous *Shh* signalling can instate posterior positional memory

Next, we hypothesised that *Shh* signalling could be the cue that instructs reprogramming of anterior cells into posterior memory cells. In the anterior-to-posterior transplantations, transplanted anterior cells would have been exposed to *Shh* secreted by endogenous posterior cells. Thus, we repeated the anterior-to-posterior transplantation in the presence of a pharmacological *Shh* signalling inhibitor, with some modifications to the assay. One limitation of the double reporter is that cells would lose fluorescence if they lost both anterior and posterior identity. To circumvent this issue, we substituted the *Alx4*:mCherry transgene for a transgene in which the mouse *Prrx1* enhancer drives mCherry expression in all connective tissue cells regardless of positional identity. This enabled the transplant to be tracked continuously by mCherry expression, while *Hand2:*EGFP labelled posterior cells (**Fig. 6a**). Due to availability, we used hindlimbs, rather than forelimbs, as donors given that positional memory codes in the salamander forelimb and hindlimb are compatible ^40^. To reduce the number of required donors, we also transplanted cells by skin grafting instead of cell injections (**Fig. 6b**). The modified assay resulted in the same outcome as the original cell injection assay; anterior cells acquired posterior identity (*Hand2*:EGFP) in the blastema. We then performed the assay while injecting test axolotls intraperitoneally at the conical blastema stage with a single dose of BMS-833923 (a *Shh* pathway inhibitor that more potently blocks *Shh* signalling in axolotls than the commonly used cyclopamine ^37^). Our data show that BMS-833923 completely prevented transplanted anterior cells from acquiring posterior identity (*Hand2*:EGFP) (**Fig. 6c**, **Extended Data Fig. 8e**). Thus, we inferred that endogenous *Shh* activity posteriorized the transplanted anterior cells.

**Fig. 6:**
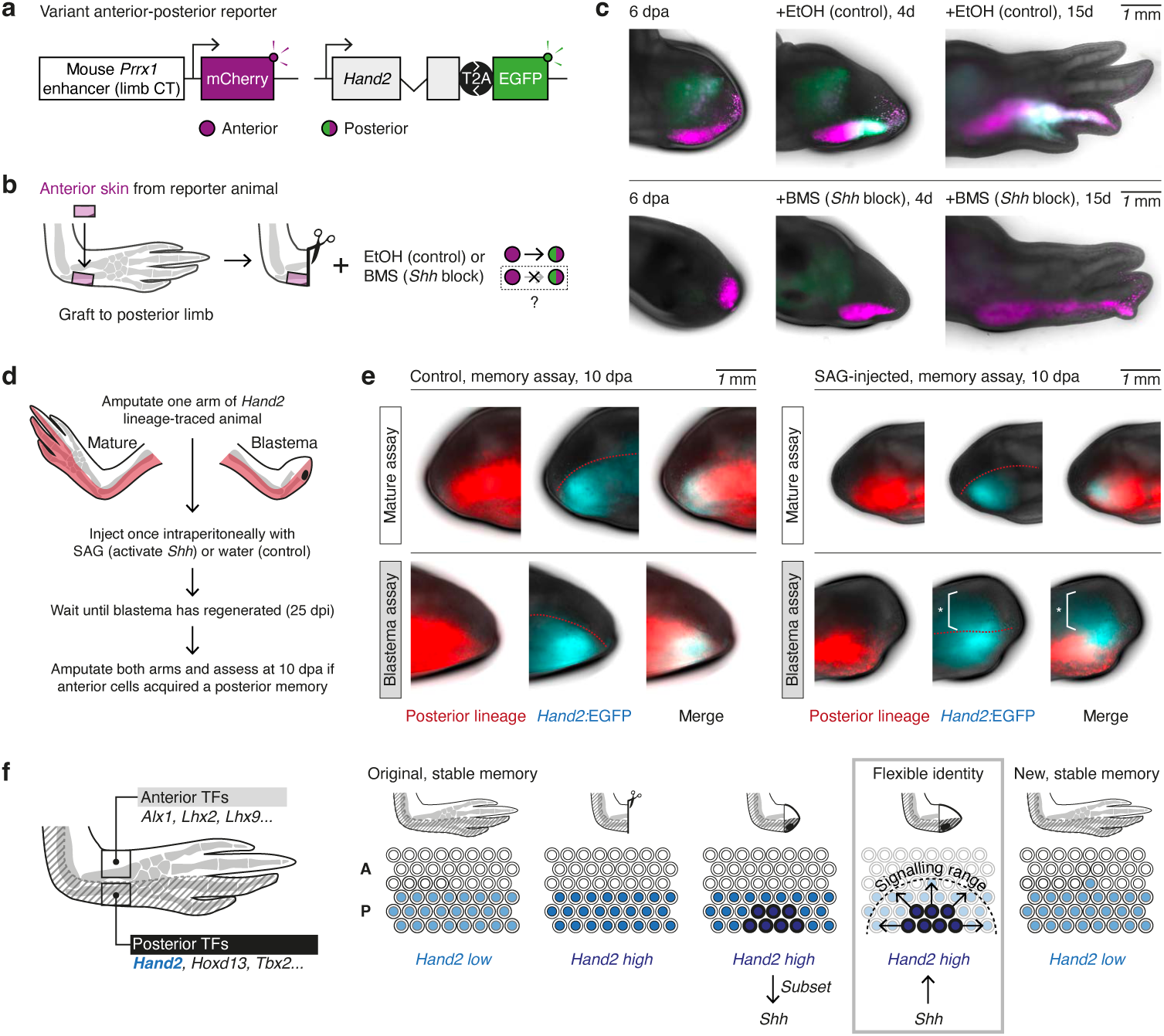
*Shh* signalling reprogrammes anterior cells with a posterior memory. **a,** A second double transgenic axolotl to label anterior and posterior cells. Prrx1>mCherry (magenta) labels all connective tissue cells, enabling them to be tracked continuously after transplantation. Hand2:EGFP (green) labels posterior cells only. **b,** Experimental strategy to test if Shh signalling is necessary for posteriorisation of anterior cells. Anterior limb skin was transplanted from the double reporter animal in (a) to the posterior limb of an unlabelled host. 4 days later, the limb was amputated through the transplant. At 6 dpa, the animal was injected intraperitoneally with 1 mM BMS-833923 (test) or ethanol (control). Hand2:EGFP expression (indicating posteriorisation) was tracked during regeneration. **c,** Result of the experiment described in (b). Blocking Shh signalling prevents upregulation of Hand2:EGFP in transplanted cells. n = 4 per condition. **d,** Experimental strategy to test if Shh signalling is sufficient to posteriorize anterior cells. Posterior cells were labelled with permanent mCherry by lineage tracing embryonic Hand2 cells as depicted in Fig. 2h. The right arm of each animal was amputated to generate a blastema and the left arm left intact. 8 days later, animals were injected intraperitoneally with 1.5 mM SAG (test) or water (control). Posteriorisation was assessed by upregulation of Hand2:EGFP reporter in anterior (mCherry-negative) cells after re-amputation. **e,** Results of experiment described in (d). Images depict 10 dpa of limbs previously treated with water (left) or SAG (right). In water-injected animals, Hand2 (cyan) is expressed within the originally posterior lineage (red), both in the mature and blastema assays (n = 4 limbs per assay). In SAG-injected animals, Hand2 becomes expressed in cells that originally had an anterior identity (asterisked region). This effect was seen in the blastema assay but not the mature assay (n = 5 limbs per assay). **f,** Model for transmission of anterior-posterior positional memory through regeneration. Positional memory is stable at steady state but is transiently flexible and subject to posteriorisation in the blastema. A: anterior. P: posterior.

### *Shh* signalling is sufficient to posteriorise positional memory during regeneration

Considering these findings, we next asked whether we could re-write positional memory in anterior cells by transiently inducing *Shh* signalling. When we delivered a baculovirus encoding *Shh* to anterior cells, we found that infected cells induced a ring of *Hand2:*EGFP expression in the surrounding blastema (**Extended Data Fig. 9a-b**). Thus, anterior blastema cells readily respond to an ectopic *Shh* source by inducing *Hand2.* During normal regeneration, the strongest *Hand2* signal is observed adjacent to the *Shh* signalling centre, indicating that posterior blastema cells similarly upregulate *Hand2* in response to *Shh* (**Fig. 2e,g**). Indeed, we found that continuously blocking *Shh* signalling during regeneration inhibited full upregulation of *Hand2* in posterior blastema cells (resulting in hypomorphic regeneration), while sparing the *Hand2* upregulation in the first 6 dpa that occurs prior to establishment of the *Shh* signalling centre (**Extended Data Fig. 9c)**. Next, we performed a lineage-traced experiment to test if transient *Shh* signalling can convert anterior cells into posterior memory cells *in vivo* (**Fig. 6d**). To this end, we prepared *Hand2* lineage tracing axolotls in which we could discriminate developmentally anterior (mCherry-) and developmentally posterior cells (mCherry+) and amputated the right limb from each animal to generate a blastema, leaving the left limb intact. Once the blastema had reached conical stage, we made a single intraperitoneal delivery of SAG (Smoothened agonist) to activate *Shh* signalling in both limbs. This approach allowed us to compare mature cells (left limb) and blastema cells (right limb) in one experiment. We found that anterior blastema cells acquired *Hand2:*EGFP expression after SAG injection (**Extended Data Fig. 9d-e**). Importantly, the positional memory of these anterior cells became stably posteriorized, as their descendants expressed *Hand2*:EGFP upon re-amputation 25 days after SAG injection (**Fig. 6e**, **Extended Data Fig. 9f**). We also validated that this effect was specific to blastema cells – anterior mature cells did not posteriorize at the same concentration of SAG (**Fig. 6e**, **Extended Data Fig. 9f**). In parallel experiments, we found that SAG-treatment of blastema cells resulted in ectopic *Shh* expression in the subsequent regeneration cycle (**Extended Data Fig. 9g**). In all, we conclude that transient *Shh* signalling is sufficient to posteriorize positional memory in anterior blastema cells.

## Discussion

How regenerative cells recall positional information to recreate the correct tissue part is a longstanding question in developmental biology and tissue engineering. Few studies have investigated changes to positional memory, whereby positional information is stably re-written in a way that that affects successive rounds of regeneration ^41^. Here, we identified a genetic circuit that maintains posterior identity in the axolotl limb and that alters positional memory when triggered in anterior cells. Our functional analyses support that *Hand2* is necessary and sufficient to induce *Shh* and, reciprocally, that *Shh* can induce *Hand2* expression in blastema cells. The positive feedback configuration between *Hand2* and *Shh*, which can sustain itself after an inducing trigger, provides a molecular explanation for how stable positional memory is created and maintained. The *Hand2-Shh* loop also explains why it is easy to posteriorize anterior cells but difficult to anteriorize posterior cells in the transplantation assay: anterior blastema cells readily activate the *Hand2-Shh* loop in response to exogenous *Shh* whereas posterior cells cannot easily disengage from the loop, supporting a ‘posterior-dominant’ encoding of positional memory.

Interestingly, *Hand2* was previously implicated in posterior identity in the regenerative zebrafish pectoral fin, although in this system, *Hand2* was not coupled to *Shh* and memory effects were not assayed ^29^. In the limb, our results indicate that *Shh* can alter positional memory in blastema cells, but not in mature cells. A similar conclusion was made for RA-induced proximalisation of axolotl limb cells, although the mechanism mediating this phenomenon and potential effects on positional memory have remained elusive ^42^. Elucidating the properties that enable blastema cells to alter their positional memory, while steady state cells remain fixed, will be an important next step. Notably, our conclusion that *Shh* can reprogram positional memory differs from previous claims made using ALM-based iterative transplantations ^13^. These authors used morphogenetic readouts but lacked reporter axolotl models for tracking anterior and posterior identity, making it challenging to locate reprogrammed cells. Regarding the dorsal-ventral axis, it has been suggested that ventral cells can dorsalise during regeneration by switching on *Lmx1b* ^9^. However, dorsal cell contamination could not be excluded in these experiments, as ventral cells were not labelled specifically. It will be interesting to test if the dorsal-ventral axis harbours hierarchical encoding of positional memory, similar to the anterior-posterior axis.

We also uncovered potentially novel features of limb development and regeneration that may shed light on evolutionary aspects of limb ontogenesis. We found that the *Shh* lineage is more widespread in axolotl limbs than in mouse limbs, in which *Shh* cells are mostly restricted to the posterior 2.5 digits of the hand (**Extended Data Fig. 3a**) ^43^. The expanded *Shh* lineage in axolotls is consistent with the known, more proximal location of *Shh* cells during development and could explain why the *Shh* mutant phenotype is more severe in axolotl limbs than in mouse ^20,37,44,45^. Our data support that expression level impacts on *Hand2*’s ability to induce *Shh*. We showed that ablating positional discontinuity through uniform expression of *Hand2* prevented limb outgrowth. It remains an open question whether this is an axolotl-specific phenotype arising from the unique configuration of anterior-posterior positional discontinuity in salamanders, or if a similar result can be achieved in other vertebrates. We found that the sole addition or removal of *Hand2* during development can alter the positional identity relevant for engineering accessory limbs post-embryonically.

We can now propose a model for how anterior-posterior memory is propagated through a limb regeneration cycle (**Fig. 6f**). At steady state, the anterior and posterior halves of the axolotl limb have fixed positional memories characterised by weak expression of embryonic patterning transcription factors. Of these, *Hand2* is necessary and sufficient to encode posterior identity and is responsible for the competence to express *Shh*. After amputation, *Hand2* expression rises significantly, first in a *Shh*-independent manner and, after *Shh* induction, in a *Shh*-dependent manner. *Hand2* induces the *Shh* signalling centre in a subset of posterior cells, possibly mediated by the first, smaller increase in its own expression level. Because blastema cells have a flexible positional memory, *Shh* emanating from the *Shh* signalling centre stimulates cells residing in the posterior half of the blastema to acquire a posterior memory (including *Hand2* expression), which they inherit into the regenerated limb. Thus, positional memory is refreshed during each regeneration cycle. If anterior cells experience *Shh* during this flexible window, they become reprogrammed into posterior memory cells (**Extended Data Fig. 10a-c**). As regeneration completes, *Hand2* expression declines but is weakly retained in posterior memory cells, while *Shh* expression is extinguished. An unexpected outcome of this model is posterior dominance: an overall tendency to posteriorize positional memory during regeneration. Important future directions will be to understand how *Shh* cells are induced as a subset of *Hand2* cells, and how the *Shh* signalling range is controlled to prevent over-posteriorisation.

Our work has implications for designing synthetic tissues for regenerative therapy or disease modelling ^3^. We showed that altering positional memory can stably change signalling centre outputs from regenerative cells. At the same time, positional memories are non-equivalently and hierarchically encoded, with a posterior dominance in the case of the limb. Elucidating genetic rules, such as these, for establishing and altering spatial information in adult cells creates opportunities to engineer synthetic tissues with stable yet contextually switchable functions. An exciting prospect would be to programme patient cells with synthetic cellular memories so that, in response to diverse injury or disease stimuli, these cells express pre-determined signalling outputs in spatial domains conducive to repair or regeneration.

## Methods

### Axolotl husbandry

Axolotls (*Ambystoma mexicanum*) were raised in individual aquaria in Vienna tap water and experiments were performed in accordance with local ethics guidelines and License GZ:51072/2019/16 issued by the Magistrate of Vienna (Genetically Modified Organism Office and MA58, City of Vienna, Austria). Axolotl matings were performed by the animal care team at the IMP. Axolotl surgeries, live imaging and tissue harvesting were performed under anaesthesia in 0.015% benzocaine (Merck E-1501) diluted in Vienna tap water, with benzocaine preparation as described in ^46^. All limb amputations were performed through the middle of the lower arm (zeugopod), unless indicated otherwise. Axolotl sizes are reported in cm, measured from snout to tail.

### Axolotl genome and transcriptome reference

Genome assembly AmexG_v6.0-DD and transcriptome assembly AmexT_v47 ^47^.

### Isolation of anterior and posterior dermal cells for RNA-sequencing

Axolotl embryos of genotype tgSceI(Mmu.*Prrx1:TFPnls-T2A-ERT2-Cre-ERT2; Caggs:loxP-GFP -loxP-mCherry*)^Etnka^ were treated with 4-OHT (Merck H7904) as described in ^48^ to permanently label connective tissue cells with mCherry, then raised individually until 12 cm. Skin (harbouring dermal connective tissue cells) was harvested from the lower arms (zeugopods) then dissected into anterior and posterior halves, leaving a gap in between. 2 replicates were prepared for anterior and 3 for posterior, with each replicate deriving from 8 axolotls (16 lower arms). Anterior and posterior samples were dissociated into single cell suspensions using Liberase TM enzyme (Merck 5401119001) as described in ^49^, with the following modifications: enzymatic digestion was performed for 50 mins and the cells were filtered through a 50 μm Filcon filter (BD Biosciences 340630). mCherry-positive cells were purified from each replicate by FACS (FACSAria III Cell Sorter, BD Biosciences) using a 100 μm low pressure nozzle and collected into separate tubes of cold amphibian culture medium ^50^. Each replicate was pelleted at 300 *rcf* for 4 mins at 4 °C, then re-suspended in 500 ul of TRIzol reagent (Thermo Fisher Scientific 15596026). RNA was extracted according to the manufacturer’s protocol and stored at −70 °C until required.

### QuantSeq library preparation and RNA-sequencing

Libraries for RNA-sequencing were prepared using QuantSeq 3’ mRNA-Seq Library Prep Kit FWD (Lexogen) using 20.25 ng of input RNA per sample. Input was 4.5 ul of input RNA plus 0.5 ul of ERCC RNA Spike-In Mix (Thermo Fisher Scientific 4456740) pre-diluted 1:10,000 in water. Samples were multiplexed for sequencing, using i7 indices 7023 (CACACT, anterior rep 1), 7025 (TTTATG, anterior rep 2), 7022 (GGAGGT, posterior rep 1), 7024 (CCGCAA, posterior rep 2) and 7026 (AACGCC, posterior rep 3). Each replicate was sequenced to a depth of 120 M reads, in SE 100 mode, distributed over 3 lanes of a HiSeq 2500 with v4 chemistry (Illumina). Sequencing was performed by the Next Generation Sequencing Facility at Vienna BioCenter Core Facilities (VBCF), member of the Vienna BioCenter (VBC), Austria.

### Gene expression analysis

Adaptor sequences were trimmed from the raw sequencing reads with Trimmomatic (version 0.39) ^51^, using parameters: ILLUMINACLIP:Adapters.fa:2:30:7 SLIDINGWINDOW:4:20 MINLEN:40 in single-end mode. Trimmed sequenced reads were mapped to axolotl genome AmexG_v6.0-DD with HISAT2 ^52^, using parameters: --no-unal –summary-file Output.log -k 5 –very-sensitive -x DBGenome -U Reads.fq.gz > Alignment.sam. featureCounts ^53^ was used to generate a read counts table. Differential expression analysis was performed on 2 anterior replicates and 3 posterior replicates using R version 4.1.2 and DESeq2 ^54^ version 1.34.0 with an FDR cutoff of *p* < 0.01. Volcano plots were generated using ggplot2 version 3.3.6 ^55^. Heatmaps were generated with the pheatmap package version 1.0.12 (R. Kolde). Gene Ontology analysis was performed with the topGO package version 2.46.0 (Alexa A, Rahnenfuhrer J) using parameters: ontology = “BP”, geneSelectionFun = topDiffGenes, annot = annFUN.org, mapping = “org.Hs.eg.db”. To calculate significant GO terms, Fisher’s exact test was performed with the “elim” algorithm. To facilitate interpretation of the differential expression results, we generated a custom gene nomenclature derived from the AmexT_v47 transcriptome. We concatenated each axolotl gene identifier with the gene symbol for the direct human homologue where available or, if not available, the closest homologue from the NCBI non-redundant database.

### Axolotl transgenesis

Plasmids for axolotl transgenesis were assembled by Gibson Assembly, amplified using Plasmid Maxi Kits (Qiagen 12163) and verified by Sanger sequencing before egg injection. One-cell stage axolotl eggs were surface sterilised twice x 5 mins with ∼0.004% Sodium hypochlorite solution (Honeywell 71696) diluted in Vienna tap water, then washed well with fresh tap water. The following steps were performed as described in ^46^. Eggs were de-jellied using sharp forceps in 20% Ficoll (Merck GE17-300-05)/1X MMR/Pen-Strep (Merck P0781) solution, then held in 10% Ficoll/1X MMR/Pen-Strep solution until microinjection. For microinjections, borosilicate glass capillary needles with filament (Harvard Apparatus, GC100F-15) were pulled using a Flaming/Brown Micropipette Puller P-97 (Sutter Instrument Co.) with settings *P* = 500; heat = 530 (Ramp test + 30); pull = 100; velocity = 120; time = 150. 5 nl of appropriate injection mix was injected into each de-jellied egg, delivered in 2 x 2.5 nl shots. Egg injections were performed at an Olympus SZX10 microscope using a PV830 pneumatic Picopump (World Precision Instruments) with settings: Vacuum Eject, Regulator 25, Range 100 ms, Timed, Duration 10-0. Injected eggs were transferred to 5% Ficoll/0.1X MMR/Pen-Strep solution overnight. The following morning, healthy eggs were transferred to individual wells of a 24-well multiwell plate (Thermo Fisher Scientific 142475) filled with 0.1X MMR/Pen-Strep solution. Embryos were screened for fluorescent transgene expression at embryo stage 42 using an AXIOzoom V16 widefield microscope (Zeiss). Axolotl lines are named according to the convention established in ^56^.

The following axolotl lines were generated by ‘random insertion’ I-SceI meganuclease-mediated transgenesis: (1) “ZRS>TFP” (tgSceI(*ZRS:TFPnls-T2A-ERT2-Cre-ERT2*)^Etnka^), (2) “Prrx1>mCherry” (tgSceI(*Mmu.Prrx1:mCherry*)^Etnka^), (3) “Prrx1>mCherry-*Hand2*” (tgSceI(*Mmu.Prrx1:mCherry-Hand2*)^Etnka^). Injection mix was prepared according to ^57^: Transgene plasmid 1 μg, I-SceI enzyme (NEB R0694) 5 units, CutSmart buffer (NEB) 1X, Water to 10 μl. In Prrx1>mCherry-*Hand2*, we fused mCherry to the N-terminus of *Hand2,* connected by a Glycine-Serine rich linker of the sequence SGGGGSGGGGS. In ZRS>TFP, the following CMV minimal promoter was used.

>CMV minimal promoter (56 bp) GGCGTGTACGGTGGGAGGTCTATATAAGCAGAGCTGGTTTAGTGAACCGTCAGA TC

The following axolotl lines were generated by NHEJ-mediated CRISPR/Cas9 knock-in: (1) “*Hand2*:EGFP” (tm(*Hand2^t/+^:Hand2-T2A-EGFP*)^Etnka^) (2) “*Hand2* lineage tracer” (tm(*Hand2^t/+^:Hand2-P2A-EGFP-T2A-ERT2-Cre-ERT2*)^Etnka^), (3) “*Alx4*:mCherry” (tm(*Alx4^t/+^:Alx4-T2A-mCherry*)^Etnka^). We followed the protocol in ^58^, using the following injection mix: Cas9-NLS protein 5 μg, gRNA 4 μg, targeting construct 0.5 μg, Cas9 buffer 1X, Water to 10 μl. Cas9-NLS protein and buffer were synthesized by Vienna Biocenter Core Facilities.

The following transgenic axolotls were published previously: tgSceI(*Caggs:loxP-GFP-dead(Stop)-loxP-mCherry*)^Etnka (ref 8)^, tgSceI(*Caggs:loxP-GFP-loxP-mCherry*)^Etnka (ref 48)^, tgSceI(Mmu.*Prrx1:TFPnls-T2A-ERT2-Cre-ERT2*)^Etnka (ref 48)^.

### Further details on generating 3’ knock-in axolotls by NHEJ-mediated CRISPR/Cas9

We generated and characterised efficient sgRNAs targeting the last intron of *Hand2* or *Alx4* following the protocol in ^58^. sgRNAs were produced by assembly and *in vitro* transcription of a synthesised DNA template. The following forward oligos, harbouring the sgRNA target sequence in the *Hand2/Alx4* intron (lower case) plus T7 promoter (underlined), were ordered at 100 μM concentration from Merck. They were PCR amplified with the universal oligo_reverse, also at 100 μM, then purified to generate DNA templates for *in vitro* transcription.

>*Hand2* sgRNA oligo_forward GAAAT<u>TAATACGACTCACTATAGG</u>atgctgtcctctaaaccgGTTTTAGAGCTAGAAATAGC

>*Alx4* sgRNA oligo_forward GAAAT<u>TAATACGACTCACTATAGG</u>ttcactactggtaaatacGTTTTAGAGCTAGAAATAGC

>Universal oligo_reverse AAAAGCACCGACTCGGTGCCACTTTTTCAAGTTGATAACGGACTAGCCTTATTTTAACTTGCTATTTCTAGCTCTA AAAC

*In vitro* transcription was performed using a MEGAscript T7 transcription kit (Thermo Fisher Scientific AM1334) and 500 ng of purified template in a 20 μl reaction. The transcription reaction was performed overnight at 37 °C. DNA template was removed by adding 1 μl of TURBO DNase (Thermo Fisher Scientific AM2238) for 15 mins. RNA was purified through LiCl precipitation. Purified sgRNA was re-suspended in water at a concentration of 1 μg/μl and stored at −70 °C.

Knock-in constructs were generated by PCR amplification and Gibson Assembly of the following components: (1) ∼400 bp of the last intron (harbouring the sgRNA target sequence) plus the complete last exon of *Hand2/Alx4* minus the stop codon, (2) the transgenes to be knocked in (e.g. T2A-EGFP) plus stop codon, (3) Poly(A) sequence from SV40 (*Hand2* knock-ins) or rabbit β-globin (*Alx4* knock-in).

>*Hand2* last intron plus <u>last exon minus stop codon</u> (650 bp) GGCCGCGGACATTAGGCGACGTAAAGAAAGGCCCATCGCAGCCGCGGCCTGTATTtTCGCGGATAATGCCTGCGCCGCGTCTGGAGGGGCAGATATAATCCCCAGCTCCACGGCAGCCCTTCAGATGTGGCGATTGCCTCGGTTTAGAGGACAGCATTTACATAGCTTTCAGGTGAACTTGAGTATGAATCGCAATCACTCGTGTTGTCTTTCTCTCTCTCTGTGTATCCCCCTCCCCCTCTCTCTTTTTATATATATATATATATATATATTGCaGTTTCGCCTACAACTGTGGCCCTGTCTGTCTGCTAAAAAGGGGGGAATTGGCAAGT GCGTGTTGCTGAAGGCTGTAGTGCGGTGTGTGTGCGTGTATATATGTTACGTAGAGATACATAGATACATATCCGTGTTACGTGTTACGAATTCGTGCGTGTGTGTGTGC GCGATTATCCGCGTTGGTTGTGAACACATGTTTGGGTCTGCAGCAAATCAACATTCAATTGTGAGATATTGAGTTCTCTTTGCTTTTGTCTCCCTTCCCGCTCTCTTGCCA G<u>AATGAACTCTTGAAAAGTACCGTCGGCAGCAACGACAAGAAGAGCAAGGGCA</u> <u>GGACTGGCTGGCCTCAGCACGTCTGGGCCTTGGAGCTCAAGCAG</u>

> *Alx4* last intron plus <u>last exon minus stop codon</u> (679 bp) GACATGTAGGGGCAATCTGAAGTCCCACTCAAAGCCCACCTAGAACCGTCCCTGCTCAGCTGGGGGAAGGCAGAATCAAATTTTGTGGAAGGCAGTCCTGTAACTCGCACCCAGAACTCTACAGCCTGTACACTGAAATATAATCAAATGGTGTTGATAATTCACAATGTGATTCACTACTGGTAAATACCGGTAACACTGAACCGCTGAGCGACATCATACAACATATTTCAAATTGGTATTAATTGTATAATGTTCCTATACTCGTCTCTTGCTGTAAATCTTATTTATTGCCTCCAGCCCTCCAAATAGTGCCACTTTCTCATTCC TTGTCTACTTTTGTCTTCTCCTGTTACAG<u>ATCCAGAACCCAACATGGATTGGAAAC</u> <u>AACAGCGGGGGCTCTCCGGTGGCAGCCTGTGTGGTCCCCTGTGACACCGTCCCAT</u> <u>CCTGCATGTCTCCTCATGGCCACCCCCATGCAAGTGGAGGTGTTTCTGAATTCCT</u> <u>GAGCGTGCCTAGCTCAGGAAGCCACATGGGTCAGGCACACATGGGTAACCTCTT</u> <u>TGGCACTGCTGGGCTCAGCACAGGCATCAATGGCTACGACCTCAACGTGGAGCC</u> <u>AGACCGCAAGACCTCCAGCATCGCAGCCCTGCGGATGAAGGCCAAGGAGCACAG</u> <u>TGCCGCCATCTCCTGGGCCACA</u>

### Codon alteration

The following sequences were codon altered to enable them to be distinguished from endogenously expressed mRNA: axolotl *Hand2* in Prrx1>mCherry-*Hand2* and axolotl *Shh* in BV-*Shh* baculovirus. An axolotl codon usage table was generated using the first transcript isoform from each gene annotated in axolotl transcriptome assembly AmexT_v47. Optimizer (http://genomes.urv.es/OPTIMIZER/) ^59^ was used with method “Guided random” to alter codons while still reflecting axolotl codon usage.

>Codon-altered axolotl *Hand2* ORF (645 bp) ATGAGCCTGGTGGGCGGTTTCCCACACCACCATCCTGGCGTGCATCACCACCATGAGGGCTACCCCTTTTCGGCCGCCGCAGCAACGGGGAGATGCCACGAAGACTCGCCATACTTTCATGGTTGGCTTATCGGTCATCCGGAGCTCTCGCCTCCCGATTATGGTCCAGGAGCACCCTACAGTCCTGAATATGGAGGGGGGGGCGGCCTTGAACTATGCGGGCCTGGGGGCGCGCCAGGGGGAGGAGCCGGAGCGCTTCTCTCAACTAGACCTGTGAAGCGGCGAGGCACCGCTAATAGGAAGGAGCGGCGGAGAACCCAAAGCATCAACAGTGCTTTCGCTGAGCTCCGGGAATGTATCCCGAATGTGCCAGCCGACACGAAGTTGTCAAAGATCAAAACTTTGCGTCTAGCCACTTCTTATATCGCCTACCTGATGGATTTGCTTGCCAAGGATGAGCAGTCTGAAGCCGAAGCTTTCCGGGCAGATCTGAAACAGAGGGGAGGGGGTGGGGCTGAGTGTAAGGAAGATAAAAGAAAGAAGGAGTTGAATGAATTGCTGAAGTCCACAGTCGGGAGTAACGACAAGAAATCCAAGGGTCGCACCGGTTGGCCACAGCATGTGTGGGCGCTAGAGCTCAAGCAG

>Codon-altered axolotl *Shh* ORF (1269 bp) ATGCGTCTCCTCCTTCGCCGGCTACTGCTGGGTACCTTGGTTTGGGCACTGCTAGTGCCCAGCGGCCTGACTTGCGGCCCGGGGCGTGGTATCGGTAAAAGGAGACAGCCTAAAAAACTGACACCCCTCGCGTACAAGCAGTTTATCCCCAACGTCGCGGAGAAGACACTGGGAGCATCTGGACGTTATGAGGGGAAGATCACTAGGAACTCTGACCGTTTCAAGGAGCTCACTCCTAATTACAACCCCGACATCATTTTTAAGGACGAGGAG AATACAGGAGCTGACCGACTGATGACTCAGAGGTGCAAAGACAAACTGAATGCCCTGGCTATTAGCGTAATGAATCAGTGGCCGGGCGTGAAACTGCGGGTGACGGAA GGCTGGGATGAAGATGGTCATCACAGTGAGGAGAGTCTGCATTACGAGGGCCGAGCCGTGGATATCACAACCTCTGACCGTGACAGGTCTAAGTATGGAATGCTGGCACGTCTGGCCGTGGAGGCAGGCTTTGATTGGGTCTACTTCGAGTCCAAGGCCCACATACATTGCAGCGTGAAGGCGGAGAACAGTGTGGCAGCCAAGTCGGGAGGATGTTTTCCGGCCAGTGCTAAGGTTACACTGGAACATGGCGTTACGAGACCAGTGAAGGATCTGCGACCCGGAGACCGTGTGCTAGCAGCAGATGGACAAGGTCGACTGGTTTATAGCGACTTTCTTATGTTTCTCGACAAAGAAGAGGCAGTGACAAAGGTCTTTTACGTCATTGAGACGGAGAGACCAAGGCAGAGGCTAAGGTTGACAGCAGCCCACCTCCTGTTCGCCGCAAGGCATCCCGCAAACTCATCTAGCTCCACCGGGTTCCAAAGTATCTTCGCATCAAGGGTTCGACCTGGGCACCGGGTGCTTACTGTCGACCAGGAAGGACGGGGGCTTCAGGAGGCTACTGTCACTCGCGTGTACCTGGAGGAGGGTGCCGGAGCCTACGCCCCCGTTACCAGTCATGGAACCGTTGTGATTGACAAGGTACTCGCCAGTTGCTACGCAGTGATCGAGGAGCATTCCTGGGCCCACTGGGCTTTTGCCCCTCTGCGACTTGGCTACGGCATACTGAGCATCTTTTCCCCTCAAGATTACAGCCCACATAGTCCCCCCGCGCCTAGCCAGAAAGAAGGCGTGCATTGGTACTCAGAAATCCTGTATCATATAGGGACATGGGTGCTGCATAGCGACACTATTCACCCCTGGGGCATGGCCGCCAAGTCGAGT

### Generation of *Hand2* CRISPant axolotls

The following mix was microinjected into fertilised one-cell stage eggs, following the protocol in ^58^: Cas9-NLS protein 5 μg, sgRNA1 2 μg, sgRNA2 2 μg, Cas9 buffer 1X, Water to 10 μl. The target sequences for sgRNA1 and sgRNA2 flank the *Hand2* translational start. Control mix contained all components apart from the sgRNAs.

>*Hand2* CRISPant sgRNA1 GCGGCCCCTGGGAGGCCC

>*Hand2* CRISPant sgRNA 2 ACCCCAGGGTGGTGGTGA

### Estimation of indel frequency in *Hand2* CRISPants

Genomic DNA was individually extracted from *Hand2* CRISPant or control sibling limbs. Amputations were performed through the middle of the lower arm and each offcut was harvested into 50 μl of 50 mM NaOH. Harvested tissue was heated to 95 °C for 12 mins, then cooled to 4 °C in a thermocycler. 5 μl of 1 M Tris, pH8 was added, then the extracted DNA was stored at 4 °C until genotyping. For genotyping, 1 μl of extracted DNA was PCR amplified using KAPA HiFi HotStart ReadyMix (Roche 07958927001) and primers that generate a ∼750 bp amplicon surrounding the *Hand2* translational start. 30 ng of PCR-amplified and purified DNA was Sanger sequenced using Hand2_sequencing primer at the IMP/IMBA Molecular Biology Service. Indel frequency was estimated from Sanger sequencing results using the ICE Analysis Tool (Synthego, https://ice.synthego.com/).

>*Hand2*_genotyping_F GAAGTAGCAGGGATGGACGAG

>*Hand2*_genotyping_R AAGGCGCTGTTGATGCTCT

>*Hand2*_sequencing CACAGGCCAGGACTTCAAGAA

### Determination of axolotl ZRS enhancer

The axolotl ZRS enhancer was determined by multiple species alignment of the following genome sequences using mVISTA ^60^ and PipMaker ^61^.

Axolotl (*Ambystoma mexicanum*) assembly AmexG_v6.0-DD chr2p:694366863-694689506

Human (*Homo sapiens*) assembly hg38 chr7:156769228-156790956

Mouse (*Mus musculus*) assembly mm10 chr5:29292950-29323801

Chick (*Gallus gallus*) assembly Gal6 chr2:8538956-8559114

Fugu (*Takifugu rubripes*) assembly fr3 chr10:5739579-5747090

>Axolotl ZRS enhancer, AmexG_v6.0-DD chr2p:694613160-694613980 (821 bp) ACCTTAATATCCATCTTTGCATTTGAAGTTGTTGCATAAAATGTACCACGAGCGACAGCAACATCCTGACTAATTAGCCAAATTACCCAGACATCCCTCCAAAAAAGCCGCGAAACAGAGAGCATGTCTGTCGGATTAAAAGGTTGTAACTCCTAAAACATCAAACGGAGCGCCAGATAATAAAAGCCAATCGTACAGAAATTTGAGGTAACTTCCTTGCTTAATTAATTAGCTAGGCCAGTTGGAGCGAGGAGGCCAACGCGGGCGCGTAGAACGCCCATAAAGCTGAACAACTCGACAGCACAAAAGTGGAGAAACAAAGATTTTTTAATATGCGTCTATCCTGTGTCACAGTTTGAAATTGTCCTGGTTTATGTCCCTTTTGGCAAAGTTACAATAAAAGTGACCCTGTACTGTATTTTATGGCCAGACGACTTTTCGTTTTGTTCCCGGTGACTAATTTGACTCAGGCCCCCATCTTGAATAGACACAGAAAGGGGCCGGGGGAATGAGGCTGTCTGTCTCGCTTGGGTTTCATTGCATTTTTTCATTATTCGGGCTCGTTTTTCGCCACAGATCATCCATAAATTGTTGGAAATGAGTGATTAAGGAAGTGCTGCTTAATGTTAGTAGCACACATTCTTTGTGCGTTTCACCCTCCCGCCCCCTCCATTTTGTGGGTGAGAGGAAATCAAGTAATGCAGAAACAATAAGGAAGCCTCCTGCTGGGAACCTTTCAAGGAAATGTAACCTGCATACTGTTTTGATCTCGGTGTTCCTTTCAGAGTATGCCGCGATGTTTCAACAGCTATTTTCATGTG

### Genetic lineage tracing (ZRS / *Hand2*)

For lineage tracing, either ZRS>TFP axolotls or *Hand2* lineage tracing axolotls were mated with loxP-mCherry fate mapping axolotl of genotype: tgSceI(*Caggs:loxP-GFP-dead(Stop)-loxP-mCherry*)^Etnka^ ^(ref^ ^8)^. To induce Cre/loxP recombination during development, stage 42 ZRS or *Hand2* lineage tracing embryos were bathed overnight dark in 500 ml of 2 μM 4-OHT, as described in the water-based method of ^46^. To induce Cre/loxP recombination in *Hand2* cells of the mature limb, 7 cm axolotls were bathed individually and overnight dark in 100 ml of 5 μM 4-OHT. After treatment, axolotls were transferred to tap water and allowed to recover for one week. The same 5 μM 4-OHT treatment and one week recovery was repeated twice more, for a total of three treatments. Animals were screened for Cre/loxP recombination and mCherry expression using an AXIOzoom V16 widefield microscope (Zeiss).

### Surgical depletion of embryonic *Shh* cells

ZRS lineage traced axolotls were prepared by treating stage 42 embryos with 2 μM 4-OHT, as described in the section “Genetic lineage tracing”. After growing to 6 cm size, the left arm of each axolotl was depleted for ZRS lineage cells by using microscissors to excise tissue posterior to the ulna in the lower arm. Successful depletion was confirmed by imaging loss of mCherry fluorescence using an AXIOzoom V16 widefield microscope (Zeiss). The right arm of each animal was treated as a control and depleted for an equivalent amount of tissue anterior to the radius instead. 2 days post-surgery, each arm was amputated through the distal part of the lower arm, distal to the depleted region. Images were acquired every few days post-amputation to assess onset of ZRS>TFP expression in depleted *versus* control limbs. mCherry depletion efficiency was estimated by comparing the area of mCherry-positive tissue in the blastema in widefield images acquired from control and depleted animals.

### Tissue sections: preparation, staining and imaging

Samples were fixed overnight in 4% PFA (paraformaldehyde), pH 7.4, at 6 °C. The following morning, samples were washed well with cold PBS then equilibrated with the following solutions at 6 °C for one overnight each: (1) 20% sucrose/PBS, (2) 30% sucrose/PBS, (3) 1:1 mix of 30% sucrose/PBS and Tissue-Tek O.C.T. compound (Sakura). Samples were mounted in Tissue-Tek O.C.T. compound, frozen on dry ice and stored at −70 °C until sectioning. Cryosections of 16 μm thickness were prepared using a Cryostar NX70 (Thermo Fisher Scientific) and collected on Superfrost Plus adhesion microscope slides (Epredia J1800AMNZ). Slides were stored at −20 °C until required. For staining, slides were brought to room temperature then washed well with PBS to remove O.C.T. before proceeding to the following steps. *DAPI staining only*: slides were incubated with DAPI 1:1,000 in PBS + 0.2% Triton X-100 for 1 hour at room temperature, then washed well with PBS + 0.2% Triton X-100 before mounting. *Staining with anti-Prrx1 antibody*: slides were blocked for 30 mins at room temperature with PBS + 0.2% Triton X-100 + 1% normal goat serum (NGS), then incubated overnight at 6 °C with rabbit anti-Prrx1 antibody (^48^) diluted 1:500 in PBS + 0.2% Triton X-100 + 0.1% NGS. The following day, slides were washed well with PBS + 0.2% Triton X-100 then incubated for 2 hours at room temperature with Alexa 647-conjugated anti-rabbit secondary antibody diluted 1:500 in PBS + 0.2% Triton X-100. Slides were washed well with PBS + 0.2% Triton X-100 before mounting. *For HCR in situ hybridisation*: slides were stained according to the HCR RNA-FISH protocol for fixed frozen tissue sections (Molecular Instruments), omitting post-fixation and Proteinase K treatment. Probe hybridisation buffer, wash buffer and amplification buffer were purchased from Molecular Instruments. Samples were mounted in Abberior Mount liquid antifade mounting media (Abberior) for imaging. Images were acquired with a LSM980 AxioObserver inverted confocal microscope with ZEN software (Zeiss), plus AiryScan 2 for HCR experiments only.

### Wholemount samples: preparation, HCR staining, tissue clearing and imaging

Samples were fixed overnight in 4% PFA (paraformaldehyde), pH 7.4, at 6 °C. The following morning, samples were washed well with cold PBS then dehydrated progressively through ice cold 25% methanol/PBS, 50% methanol/PBS, 75% methanol/PBS and 100% methanol (30 mins each). Samples were kept for one overnight in 100% methanol at −20 °C. The following day, samples were re-hydrated progressively through ice cold 75% methanol/PBS, 50% methanol/PBS, 25% methanol/PBS and PBS (30 mins each). Samples were washed twice more with cold PBS then stained for *Shh* transcripts using the HCR RNA-FISH protocol for whole-mount zebrafish embryos and larvae (Molecular Instruments), starting at the section “Detection Stage”. After completion of the HCR protocol, samples were stained overnight at 6 °C in DAPI 5 mg/ml (Sigma) diluted 1:1,000 in 5x SSC + 0.1% Tween 20. The following day, samples were washed well with 5x SSC + 0.1% Tween 20 then optically cleared overnight at room temperature on an aerial rotator in Ce3D solution Refractive Index 1.50, prepared as described in ^62^. Images were acquired in Ce3D solution using a LightSheet.Z1 microscope with ZEN software (Zeiss) and custom imaging chamber as described in ^63^.

### HCR probe design and detection

HCR probes targeting axolotl *Shh* mRNA, probe hybridisation buffer, wash buffer, detection hairpins and amplification buffer were purchased from Molecular Instruments. Sequences unique to *Shh* mRNA were identified by BLAST alignment against axolotl transcriptome assembly Amex.T_v47 ^47^. Sequences were considered not unique if they exhibited homology to non-target transcripts at more than 36 out of 50 nucleotides. *Shh* HCR signal was detected using B5 hairpins conjugated to Alexa-647 fluorophore. HCR probes targeting TFP mRNA were purchased at the 50 pmol scale from IDT (Integrated DNA Technologies, oPools), suspended in water and stored at −20 °C. TFP HCR signal was detected using B1 hairpins conjugated to Alexa-546 fluorophore.

### Quantification of *Hand2:*EGFP expression during regeneration

*For whole-tissue measurements:* 4.5 cm axolotls harbouring *Hand2*:EGFP were amputated through the middle of the lower arm and imaged throughout regeneration with identical acquisition settings using an AXIOzoom V16 widefield microscope (Zeiss). Longitudinal imaging of 6 limbs was performed on 0, 2, 4, 7, 10, 15, 21, 28 and 39 dpa. Mean EGFP fluorescence intensity was measured in Fiji software ^64^ by manually drawing a region of interest in the EGFP-positive region of the blastema. For 0, 2 and 4 dpa, no/little blastema had formed so measurements were instead taken from 500 μm behind the amputation plane. The 500 μm source zone for lower arm regeneration was established in ^65^.

*For single cell measurements*: The *Hand2*:EGFP intensity of mature arm cells, 7 dpa blastema cells and 14 dpa blastema cells were compared by flow cytometry. Lower arm tissue was harvested from 6 cm *Hand2*:EGFP axolotls. The entire lower arm was taken for mature measurements. Blastemas were generated by amputating through the middle of the lower arm 7 or 14 days prior to flow cytometry. Harvested tissues were dissociated into single cell suspensions using Liberase TM enzyme (Merck 05401127001) as described in ^49^, with the following modifications: dissociation was performed for 55 mins (mature sample) or 45 mins (blastema samples) and the cells were filtered through a 70 μm MACS SmartStrainer (Miltenyi Biotec 130-098-462). Cells were analysed by FACS (FACSAria III Cell Sorter, BD Biosciences) using a 100 μm low pressure nozzle. Mean fluorescence intensities were quantified using FLOWJO software (BD Biosciences).

### Accessory Limb Model (ALM)

ALM was performed on the upper arm as described in ^66^. For *Hand2* CRISPant ALMs, donor axolotls (F0 *Hand2* CRISPant) were 9-10 cm and host axolotls (GFP-expressing controls) were 13-14 cm. 1 mm x 1 mm donor skin grafts were transplanted distal to the deviated nerve on host animals. Donor grafts were taken from *Hand2* CRISPants deemed to have a high mutation rate as judged by regeneration of a hypomorphic spike after a previous lower arm amputation. As controls, skin grafts were taken from sibling axolotls injected with Cas9 but no sgRNA. *Hand2* CRISPant ALM and control grafts were performed on opposite arms of each host axolotl. 58 days after surgery, ALMs were harvested and fixed for skeletal staining with Alcian Blue / Alizarin Red. *Hand2* misexpression ALMs were performed similarly. Donor axolotls (strong Prrx1>*Hand2*) were 5 cm and host axolotls (unlabelled controls) were 8 cm. As controls, skin grafts were taken from weak Prrx1>*Hand2* siblings. ALM formation was deemed to have been completed by 34 days post-surgery.

### Alcian Blue / Alizarin Red staining

Skeletal staining of fixed accessory limbs was performed as described in ^67^. Stained limbs were imaged in 70% Ethanol/water using an AxioCam ERc 5s colour camera (Carl Zeiss Microimaging) mounted on an Olympus SZX10 microscope. Alcian blue (A3157), Alizarin red (A5533) and Trypsin (85450C) were purchased from Merck.

### Cell transplantations by injection of FACSorted cells

Upper and lower arms were harvested from 4 cm double reporter axolotls (*Alx4:mCherry*_*Hand2:EGFP*) and dissociated into single cell suspensions using Liberase TM enzyme (Merck 05401127001) as described in ^49^, with the following modifications: dissociation was performed for 55 mins and the cells were filtered through a 70 μm MACS SmartStrainer (Miltenyi Biotec 130-098-462). Anterior cells (mCherry-positive plus EGFP-negative) or posterior cells (EGFP-positive) were purified by FACS (FACSAria III Cell Sorter, BD Biosciences) using a 100 μm low pressure nozzle and collected into separate tubes of amphibian culture medium ^50^. Pelleting and injection of FACSorted cells into the arms of 4 cm unlabelled sibling axolotls was performed as described in ^49^, using a Nanoject II injector (Drummond Scientific Company). Anterior cells were injected into the posterior lower arm, while posterior cells were injected into the anterior lower arm. 9,000 cells were injected per experiment. Host arms were imaged at 2 days post-injection (dpi) to confirm successful transplantation using an AXIOzoom V16 widefield microscope (Zeiss). At 15 dpi, host arms were amputated distal to the transplant. Regenerating blastemas were imaged at 6, 8, 10, 15 and 26 dpa until the limb was considered fully regenerated. At 26 dpa, a second amputation was performed through the regenerated part of the limb to generate a second blastema and test if cells had altered their positional memory. This second blastema was harvested at 8 dpa and fixed and processed for wholemount HCR staining against *Shh* transcripts.

### Dilution and storage of BMS-833923 and SAG

BMS-833923 (Cayman Chemical 16240) was dissolved to 10 mM in ethanol and stored as single-use aliquots at −70 °C. InSolution Smoothened Agonist (SAG, Merck 566661) for intraperitoneal injections was purchased at 10 mM in water and stored as single-use aliquots at −20 °C. SAG for bathing experiments was purchased from Merck (566660), diluted to 40 uM and stored as single-use aliquots at −20 °C.

### Skin transplantation plus intraperitoneal delivery of BMS-833923

Skin (∼1 mm x 1 mm) was transplanted from the anterior lower leg of 7 cm double reporter axolotls (*Prrx1>mCherry_Hand2:EGFP*) to the posterior lower arm of 8 cm unlabelled *d/d* hosts, maintaining dorsal-ventral and proximal-distal directionality. 4 days after transplantation, the host arm was amputated through the distal third of the transplant and blastema outgrowth was monitored every 1-2 days using an AXIOzoom V16 widefield microscope (Zeiss). At 6 dpa (conical blastema stage), test axolotls were injected intraperitoneally with 25 μl of BMS-833923 diluted to 1 mM in water. Control axolotls instead received the appropriate dilution of ethanol in water injection. Injection mix contained Fast Green dye (Thermo Fisher Scientific) for visual contrast. Blastemas were further imaged at 4 and 15 days post-injection to assess for changes in positional identity.

### Assessing the effect of blocking *Shh* signalling on *Hand2*:EGFP expression

7 cm *Hand2*:EGFP axolotls were amputated at the top of the lower arm. Every 3 days from 0 dpa until 21 dpa, test axolotls were injected intraperitoneally with 20 μl of BMS-833923 diluted to 1 mM in water, while control axolotls were instead injected with 20 μl of water. Injection mix contained Fast Green dye (Thermo Fisher Scientific) for visual contrast. Blastemas were imaged every 3 days using an AXIOzoom V16 widefield microscope (Zeiss). Mean *Hand2*:EGFP fluorescence was quantified from manually defined regions of interest in the posterior blastema.

### SAG positional memory experiment (mature vs blastema assay)

*Hand2* lineage traced axolotls were prepared by treating stage 42 embryos with 2 μM 4-OHT, as described in the section “Genetic lineage tracing”. After growing to 8 cm size, the right arm of each axolotl was amputated through the middle of the lower arm (blastema assay). The left arm was left intact (mature assay). At 8 dpa, test axolotls were injected intraperitoneally with 20 μl of SAG diluted to 1.5 mM in water. Control axolotls instead received a water injection. Injection mix contained Fast Green dye (Merck F7258) for visual contrast. Both the blastema limb and mature limb were imaged every few days until 25 dpi using an AXIOzoom V16 widefield microscope (Zeiss). At 25 dpi, both limbs were amputated through the hand plate region to assay for effects on positional memory by expression of *Hand2*:EGFP.

### SAG positional memory experiment (*Shh* HCR assay)

5 cm *Hand2:*EGFP axolotls were amputated through both lower limbs, then bathed in water (control) or 10 nM SAG (test) for the first 21 days of regeneration. Bathing volume was 40 ml, and solution was prepared and exchanged daily, according to the protocol of ^13^. Regeneration was deemed to be complete at 30 dpa. Axolotls were raised for a further 30 days in water, to ensure complete washout of SAG from test animals. Subsequently, axolotls were re-amputated through the hand plate region to generate a new blastema in the reprogrammed part of the limb. At 9 dpa, the new blastemas were harvested and fixed for wholemount HCR staining against *Shh* mRNA, tissue clearing and light sheet imaging.

### Baculovirus production and injection

Pseudotyped baculovirus was produced as described in ^68^. BV-mCherry, a control baculovirus to misexpress mCherry, was published previously as *ch*BV ^68^. The cytomegalovirus immediate-early promoter (pCMV) drives expression of mCherry in infected cells. BV-*Shh*, to misexpress axolotl *Shh*, was generated for this study. pCMV drives expression of nuclear-localised mCherry – T2A – axolotl *Shh*. Co-translational cleavage in the T2A sequence releases full length axolotl *Shh* protein. Axolotl *Shh* was codon-altered to enable distinction of virally expressed mRNA from endogenous axolotl *Shh* mRNA.

BV-mCherry or BV-*Shh* were injected into the anterior lower arm of 4 cm *Hand2*:EGFP axolotls. Injection mix contained Fast Green dye (Merck F7258) for visual contrast. 18 days post-infection, limbs were amputated through the middle of the lower arm. The regenerating blastema was imaged every few days using an AXIOzoom V16 widefield microscope (Zeiss). At 11 dpa, blastemas were harvested for fixation, wholemount tissue clearing and imaging using a LightSheet.Z1 microscope (Zeiss).

### Image analysis

Microscope images were analysed using ZEN software (Zeiss) or Fiji software ^64^.

### Statistical analysis and data representation

Statistical analyses and graph plotting was performed using Prism software (GraphPad). Data were tested for assumptions of normality and equality of variance to determine the appropriate statistical tests to perform. No data were excluded. Mean values are reported ± standard deviation (SD). Statistical significance was defined as *p* < 0.05. All figures were assembled in Adobe Illustrator.

## End notes

## Acknowledgments

We are grateful to Tanaka laboratory members for project discussions (Pietro Tardivo), assistance with RNA-sequencing library preparation (Marco Leyva), bioinformatics (Francisco Falcon and Sergej Nowoshilow) and baculovirus production (Helena Okulski). We thank Christina Lilliehook (Life Science Editors), Osvaldo Chara (University of Nottingham) and Elad Bassat, Katharina Lust and Wouter Masselink (Tanaka laboratory) for critical manuscript feedback. We thank the animal care team for excellent axolotl care (Magdalena Blaschek, Emina Silic, Victoria Szilagyi, Andrea Lentz-Koblenc, Erika Kiligan, Elisabeth Zöllner, Tamara Torrecilla Lobos, Sandra Faltin and Wilfried Auer). We thank the BioOptics facility and the Molecular Biology Service at the IMP/IMBA Core Facilities for expert support. We thank the Next Generation Sequencing facility at the Vienna Biocenter Core Facilities for RNA-sequencing.

This work was funded by HFSP (Human Frontier Science Program) fellowship LT000785/2019-L to L.O., FFG (Die Österreichische Forschungsförderungsgesellschaft) FEMtech Praktika Studentship to S.A.P. and ERC (European Research Council) Advanced Grant 742046 (RegGeneMems) to E.M.T. For the purpose of open access, the author has applied a CC BY public copyright licence to any Author Accepted Manuscript version arising from this submission.

## Author Contributions

L.O. and E.M.T conceived the project and secured funding. L.O. performed and analysed all experiments, with support from S.A.P. L.O. and Y.T-S. microinjected axolotl eggs to generate transgenic axolotls. L.O. and E.M.T. wrote the manuscript. All authors approved the manuscript.

## Materials and Correspondence

Correspondence and requests for materials should be addressed to L.O. and E.M.T..

## References

1. Rinn, J. L. et al. A dermal HOX transcriptional program regulates site-specific epidermal fate. Genes Dev. 22, 303–307 (2008).

2. Chang, H. Y. et al. Diversity, topographic differentiation, and positional memory in human fibroblasts. PNAS 99, 12877–12882 (2002).

3. Otsuki, L. & Tanaka, E. M. Positional Memory in Vertebrate Regeneration: A Century’s Insights from the Salamander Limb. Csh Perspect Biol 14, a040899 (2021).

4. Kragl, M. et al. Cells keep a memory of their tissue origin during axolotl limb regeneration. Nature 460, 60–65 (2009).

5. Nacu, E. et al. Connective tissue cells, but not muscle cells, are involved in establishing the proximo-distal outcome of limb regeneration in the axolotl. Development 140, 513–518 (2013).

6. Silva, S. M. da, Gates, P. B. & Brockes, J. P. The newt ortholog of CD59 is implicated in proximodistal identity during amphibian limb regeneration. Dev. Cell 3, 547–555 (2002).

7. Oliveira, C. R. et al. Tig1 regulates proximo-distal identity during salamander limb regeneration. Nat. Commun. 13, 1141 (2022).

8. Kawaguchi, A. et al. Chromatin states at homeoprotein loci distinguish axolotl limb segments prior to regeneration. Biorxiv 2022.11.14.516253 (2022) doi:10.1101/2022.11.14.516253.

9. Satoh, A. & Makanae, A. Conservation of Position-Specific Gene Expression in Axolotl Limb Skin. jzoo 31, 6–13 (2014).

10. Yamamoto, S., Kashimoto, R., Furukawa, S., Ohashi, A. & Satoh, A. Lmx1b activation in axolotl limb regeneration. Dev. Dyn. 251, 1509–1523 (2022).

11. Laufer, E., Nelson, C. E., Johnson, R. L., Morgan, B. A. & Tabin, C. Sonic hedgehog and Fgf-4 act through a signaling cascade and feedback loop to integrate growth and patterning of the developing limb bud. Cell 79, 993–1003 (1994).

12. Niswander, L., Jeffrey, S., Martin, G. R. & Tickle, C. A positive feedback loop coordinates growth and patterning in the vertebrate limb. Nature 371, 609–612 (1994).

13. Nacu, E., Gromberg, E., Oliveira, C. R., Drechsel, D. & Tanaka, E. M. FGF8 and SHH substitute for anterior-posterior tissue interactions to induce limb regeneration. Nature 533, 407–410 (2016).

14. Tank, P. W. & Holder, N. The effect of healing time on the proximodistal organization of double-half forelimb regenerates in the axolotl, Ambystoma mexicanum. Dev Biol 66, 72–85 (1978).

15. Tank, P. W. The failure of double half forelimbs to undergo distal transformation following amputation in the axolotl, Ambystoma mexicanum. Journal of Experimental Zoology 204, 325–336 (1978).

16. Endo, T., Bryant, S. V. & Gardiner, D. M. A stepwise model system for limb regeneration. Dev Biol 270, 135–145 (2004).

17. Purushothaman, S., Elewa, A. & Seifert, A. W. Fgf-signaling is compartmentalized within the mesenchyme and controls proliferation during salamander limb development. Elife 8, 507 (2019).

18. Christensen, R. N., Weinstein, M. & Tassava, R. A. Expression of fibroblast growth factors 4, 8, and 10 in limbs, flanks, and blastemas of Ambystoma. Dev. Dyn. 223, 193–203 (2002).

19. Han, M. J., An, J. Y. & Kim, W. S. Expression patterns of Fgf-8 during development and limb regeneration of the axolotl. Dev. Dyn. 220, 40–48 (2001).

20. Torok, M. A., Gardiner, D. M., Belmonte, J. C. I. & Bryant, S. V. Sonic hedgehog (shh) expression in developing and regenerating axolotl limbs. J Exp Zool 284, 197–206 (1999).

21. Sagai, T., Hosoya, M., Mizushina, Y., Tamura, M. & Shiroishi, T. Elimination of a long-range cis-regulatory module causes complete loss of limb-specific Shh expression and truncation of the mouse limb. Development 132, 797–803 (2005).

22. Lettice, L. A. et al. A long-range Shh enhancer regulates expression in the developing limb and fin and is associated with preaxial polydactyly. Hum. Mol. Genet. 12, 1725–1735 (2003).

23. Carlson, B. M. Morphogenetic Interactions Between Rotated Skin Cuffs and Underlying Stump Tissues in Regenerating Axolotl Forelimbs. Dev Biol 39, 263–285 (1974).

24. Iwata, R., Makanae, A. & Satoh, A. Stability and plasticity of positional memory during limb regeneration in Ambystoma mexicanum. Dev. Dyn. 249, 342–353 (2020).

25. Yelon, D. et al. The bHLH transcription factor hand2 plays parallel roles in zebrafish heart and pectoral fin development. Development 127, 2573–2582 (2000).

26. Fernandez-Teran, M. et al. Role of dHAND in the anterior-posterior polarization of the limb bud: implications for the Sonic hedgehog pathway. Development 127, 2133–2142 (2000).

27. Charité, J., McFadden, D. G. & Olson, E. N. The bHLH transcription factor dHAND controls Sonic hedgehog expression and establishment of the zone of polarizing activity during limb development. Development 127, 2461–2470 (2000).

28. Slack, M. J. Morphogenetic properties of the skin in axolotl limb regeneration. J Embryol Exp Morphol 58, 265–288 (1980).

29. Nachtrab, G., Kikuchi, K., Tornini, V. A. & Poss, K. D. Transcriptional components of anteroposterior positional information during zebrafish fin regeneration. Development 140, 3754–3764 (2013).

30. Zúñiga, A., Haramis, A. P., McMahon, A. P. & Zeller, R. Signal relay by BMP antagonism controls the SHH/FGF4 feedback loop in vertebrate limb buds. Nature 401, 598– 602 (1999).

31. Capdevila, J., Tsukui, T., Esteban, C. R., Zappavigna, V. & Belmonte, J. C. I. Control of vertebrate limb outgrowth by the proximal factor Meis2 and distal antagonism of BMPs by Gremlin. Mol Cell 4, 839–849 (1999).

32. Galli, A. et al. Distinct roles of Hand2 in initiating polarity and posterior Shh expression during the onset of mouse limb bud development. PLoS Genet. 6, e1000901 (2010).

33. Osterwalder, M. et al. HAND2 targets define a network of transcriptional regulators that compartmentalize the early limb bud mesenchyme. Dev. Cell 31, 345–357 (2014).

34. Donnelly, M. L. L. et al. Analysis of the aphthovirus 2A/2B polyprotein ‘cleavage’ mechanism indicates not a proteolytic reaction, but a novel translational effect: a putative ribosomal ‘skip.’ J. Gen. Virol. 82, 1013–1025 (2001).

35. Felipe, P. de, Hughes, L. E., Ryan, M. D. & Brown, J. D. Co-translational, Intraribosomal Cleavage of Polypeptides by the Foot-and-mouth Disease Virus 2A Peptide*. J. Biol. Chem. 278, 11441–11448 (2003).

36. Srivastava, D. et al. Regulation of cardiac mesodermal and neural crest development by the bHLH transcription factor, dHAND. Nat Genet 16, 154–160 (1997).

37. Purushothaman, S., Aviña, B. B. L. & Seifert, A. W. Sonic hedgehog is Essential for Proximal-Distal Outgrowth of the Limb Bud in Salamanders. Frontiers Cell Dev Biology 10, 797352 (2022).

38. Logan, M. et al. Expression of Cre recombinase in the developing mouse limb bud driven by a Prxl enhancer. genesis 33, 77–80 (2002).

39. Welscher, P. te, Fernandez-Teran, M., Ros, M. A. & Zeller, R. Mutual genetic antagonism involving GLI3 and dHAND prepatterns the vertebrate limb bud mesenchyme prior to SHH signaling. Genes Dev. 16, 421–426 (2002).

40. Rollman-Dinsmore, C. & Bryant, S. V. Pattern regulation between hind- and forelimbs after blastema exchanges and skin grafts in Notophthalmus viridescens. J Exp Zool 223, 51– 56 (1982).

41. Wang, Y.-T. et al. Genetic Reprogramming of Positional Memory in a Regenerating Appendage. Current Biology 29, 4193–4207.e4 (2019).

42. Maden, M. The effect of vitamin A on the regenerating axolotl limb. J Embryol Exp Morphol 77, 273–295 (1983).

43. Harfe, B. D. et al. Evidence for an expansion-based temporal Shh gradient in specifying vertebrate digit identities. Cell 118, 517–528 (2004).

44. Kraus, P., Fraidenraich, D. & Loomis, C. A. Some distal limb structures develop in mice lacking Sonic hedgehog signaling. Mech. Dev. 100, 45–58 (2001).

45. Chiang, C. et al. Manifestation of the limb prepattern: limb development in the absence of sonic hedgehog function. Dev Biol 236, 421–435 (2001).

46. Khattak, S. et al. Optimized axolotl (Ambystoma mexicanum) husbandry, breeding, metamorphosis, transgenesis and tamoxifen-mediated recombination. Nat. Protoc. 9, 529– 540 (2014).

47. Schloissnig, S. et al. The giant axolotl genome uncovers the evolution, scaling, and transcriptional control of complex gene loci. Proc. Natl. Acad. Sci. U.S.A. 118, (2021).

48. Gerber, T. et al. Single-cell analysis uncovers convergence of cell identities during axolotl limb regeneration. Science 362, eaaq0681 (2018).

49. Lin, T.-Y. et al. Fibroblast dedifferentiation as a determinant of successful regeneration. Dev. Cell 56, 1541–1551.e6 (2021).

50. Denis, J.-F., Sader, F., Ferretti, P. & Roy, S. Culture and transfection of axolotl cells. Methods Mol. Biol. (Clifton, NJ) 1290, 187–96 (2015).

51. Bolger, A. M., Lohse, M. & Usadel, B. Trimmomatic: a flexible trimmer for Illumina sequence data. Bioinformatics 30, 2114–2120 (2014).

52. Kim, D., Paggi, J. M., Park, C., Bennett, C. & Salzberg, S. L. Graph-based genome alignment and genotyping with HISAT2 and HISAT-genotype. Nat. Biotechnol. 37, 907–915 (2019).

53. Liao, Y., Smyth, G. K. & Shi, W. featureCounts: an efficient general purpose program for assigning sequence reads to genomic features. Bioinformatics 30, 923–930 (2014).

54. Love, M. I., Huber, W. & Anders, S. Moderated estimation of fold change and dispersion for RNA-seq data with DESeq2. Genome Biol. 15, 550 (2014).

55. Wickham, H. ggplot2, Elegant Graphics for Data Analysis. (2016) doi:10.1007/978-3-319-24277-4.

56. Nowoshilow, S., Fei, J.-F., Voss, R. S., Tanaka, E. M. & Murawala, P. Gene and transgenics nomenclature for the laboratory axolotl - Ambystoma mexicanum. Dev. Dyn. (2021) doi:10.1002/dvdy.351.

57. Sobkow, L., Epperlein, H.-H., Herklotz, S., Straube, W. L. & Tanaka, E. M. A germline GFP transgenic axolotl and its use to track cell fate: Dual origin of the fin mesenchyme during development and the fate of blood cells during regeneration. Dev. Biol. 290, 386–397 (2006).

58. Fei, J.-F. et al. Application and optimization of CRISPR-Cas9-mediated genome engineering in axolotl (Ambystoma mexicanum). Nat Protoc 13, 2908–2943 (2018).

59. Puigbò, P., Guzmán, E., Romeu, A. & Garcia-Vallvé, S. OPTIMIZER: a web server for optimizing the codon usage of DNA sequences. Nucleic Acids Res. 35, W126–W131 (2007).

60. Frazer, K. A., Pachter, L., Poliakov, A., Rubin, E. M. & Dubchak, I. VISTA: computational tools for comparative genomics. Nucleic Acids Res. 32, W273–W279 (2004).

61. Schwartz, S. et al. PipMaker—A Web Server for Aligning Two Genomic DNA Sequences. Genome Res. 10, 577–586 (2000).

62. Li, W., Germain, R. N. & Gerner, M. Y. High-dimensional cell-level analysis of tissues with Ce3D multiplex volume imaging. Nat. Protoc. 14, 1708–1733 (2019).

63. Glotzer, G. L., Tardivo, P. & Tanaka, E. M. Canonical Wnt signaling and the regulation of divergent mesenchymal Fgf8 expression in axolotl limb development and regeneration. eLife 11, e79762 (2022).

64. Schindelin, J., et al. Fiji: an open-source platform for biological-image analysis. Nat. Methods 9, 676–682 (2012).

65. Currie, J. D. et al. Live Imaging of Axolotl Digit Regeneration Reveals Spatiotemporal Choreography of Diverse Connective Tissue Progenitor Pools. Dev. Cell 39, 411–423 (2016).

66. Endo, T., Gardiner, D. M., Makanae, A. & Satoh, A. The accessory limb model: an alternative experimental system of limb regeneration. Methods Mol. Biol. (Clifton, NJ) 1290, 101–13 (2015).

67. Taniguchi, Y. et al. The posterior neural plate in axolotl gives rise to neural tube or turns anteriorly to form somites of the tail and posterior trunk. Dev. Biol. 422, 155–170 (2017).

68. Oliveira, C. R. et al. Pseudotyped baculovirus is an effective gene expression tool for studying molecular function during axolotl limb regeneration. Dev. Biol. 433, 262–275 (2018).

