## Extended Figures for "Molecular basis for positional memory and its reprogrammability in limb regeneration"

#### Extended Data Fig. 1: Expression domains of transgenic axolotls generated in this study.

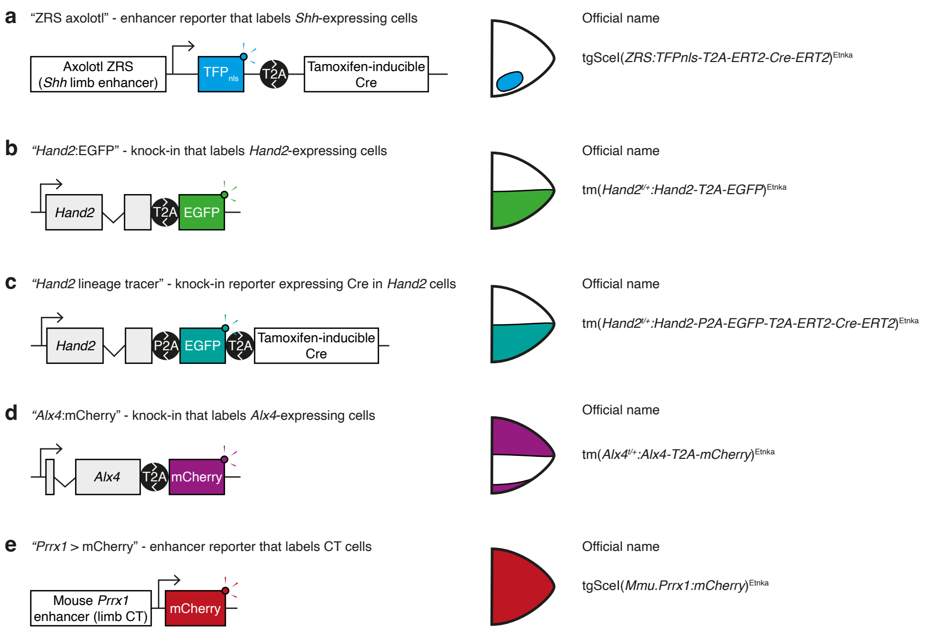


**Extended Data Fig. 1: Expression domains of transgenic reporters generated in this study.**
**a-e,** The constructs used to generate transgenic axolotls for this study (left), their expression domains in the limb bud / blastema mesenchyme (centre) and desgination according to official nomenclature (right). In the expression domain schematics, anterior is up and posterior is down. Only mesenchyme is depicted.

### Extended Data Fig. 2: Characterisation of ZRS transgenic axolotl.

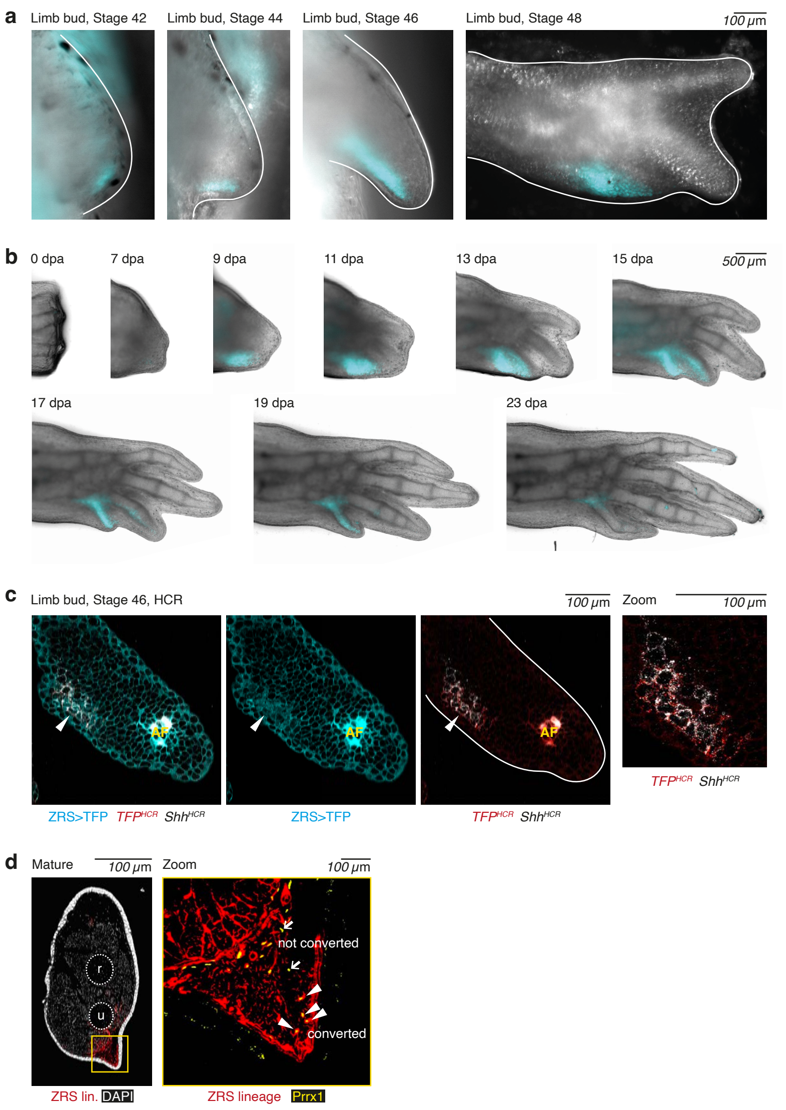


**Extended Data Fig. 2: Characterisation of ZRS transgenic axolotl.** **a,** Shh cells express ZRS>TFP (cyan) during limb development. As commonly seen in transgenic reporter animals, some perdurance of TFP protein is observed past the known Shh expression window. **b,** Regenerative Shh cells also express ZRS>TFP (cyan), but downregulate the reporter as regeneration completes. **c,** Single section confocal image of a Stage 46 ZRS>TFP limb bud. TFP-positive cells (cyan) are stained with HCR in situ hybridisation against TFP mRNA (red) and endogenous Shh mRNA (white). TFP fluorescence was not preserved well in this protocol. 81.8 ± 10.4 % of cells that expressed TFP mRNA expressed Shh mRNA. Reciprocally, 93.4 ± 8.8 % of cells that expressed Shh mRNA expressed TFP mRNA. n = 8 limb buds, with a total of 182 TFP-positive cells and 154 Shh-expressing cells quantified. AF: autofluorescence. **d,** Estimation of Cre/loxP labelling efficiency in the ZRS lineage tracing experiment. Confocal image of a cross-section of a lower arm harbouring recombined Shh lineage cells (red), with nuclei co-stained using DAPI (white). The region in the yellow box is magnified in the panel to the right. Antibody staining against Prrx1 (yellow) labels connective tissue cells. Arrows indicate examples of recombined and non-recombined Prrx1+ cells. mCherry labelling efficiency was estimated as the percentage of Prrx1+ cells posterior to the ulna bone that were also mCherry+. r: radius (anterior bone), u: ulna (posterior bone).

#### Extended Data Fig. 3: Characterisation of the axolotl ZRS lineage.

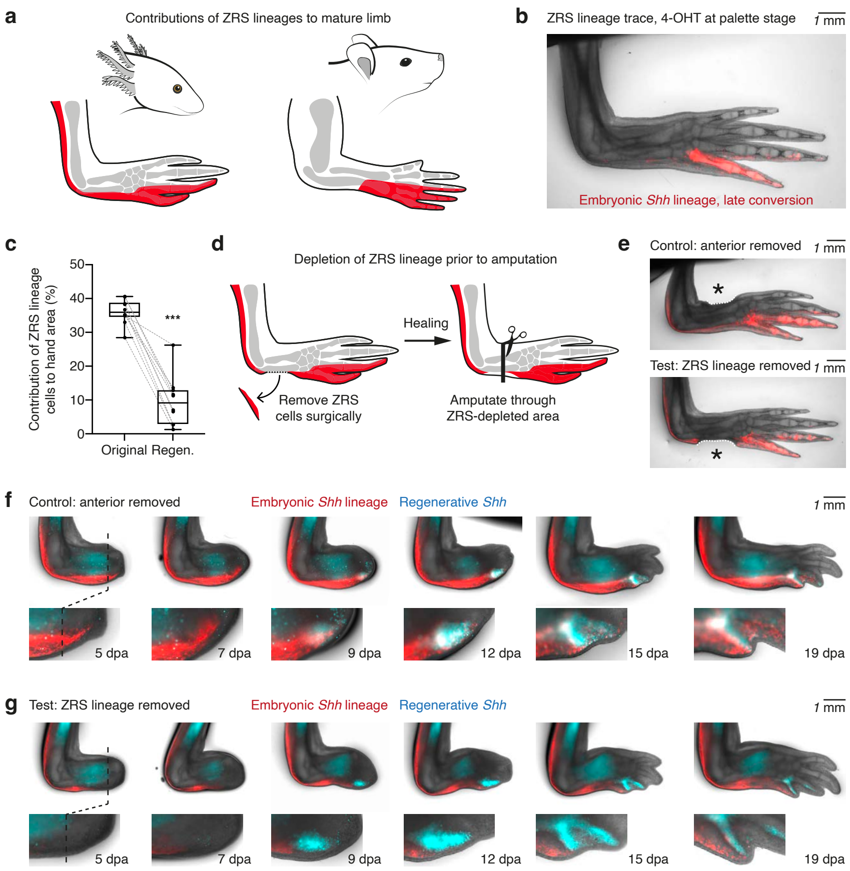


**Extended Data Fig. 3: Characterisation of the axolotl ZRS lineage.**
**a,** The contributions of embryonic Shh cells to the forelimb in axolotls (left, this study) or in mice (right, data from Harfe et al. 2004). **b,** Treating ZRS>TFP axolotls with 4-OHT at Stage 47 (instead of Stage 42 as in the other experiments) results in only the distal subset of the Shh lineage becoming labelled (n = 8 limbs). **c,** Box plot depicting the depletion of embryonic Shh cells during regeneration. The area of the hand derived from embryonic Shh cells (mCherry+) was compared in the same limbs prior to amputation (original) and after regeneration (regen.). Data from the same limb are connected by a dotted line. ***: p = 1.11e-6, paired two-tailed t-test, n = 10 limbs per condition. **d,** Strategy to deplete embryonic Shh cells prior to amputation. Micro-scissors and forceps were used to surgically excise mCherry+ tissue from the posterior of test limbs. A control surgery (removal of anterior tissue) was performed on the opposite arm of the same animal. 2 days later, both limbs were amputated through the lower arm. **e,** Control and test limbs immediately prior to amputation. Surgery sites are marked with asterisks. **f,** Regeneration time series of control limbs. Top row: overview. Bottom row: zoom-in of blastema region. The amputation plane is indicated with a dotted line. Regenerative Shh cells (cyan) appeared weakly at 7 dpa and strongly by 9 dpa. 6 out of 6 limbs expressed TFP at 7 dpa. **g,** Regeneration time series of test (Shh lineage-depleted) limbs. Limbs were amputated at the dotted line. Regenerative Shh cells (cyan) appeared with the same timing as in control limbs. Subsequent tissue morphogenesis also proceeded with similar timing to control limbs, although at later stages, proximally surviving Shh lineage cells had entered the blastema by cell migration. 6 out of 6 limbs expressed TFP at 7 dpa.

#### Extended Data Fig. 4: Gene expression in anterior and posterior dermal cells.

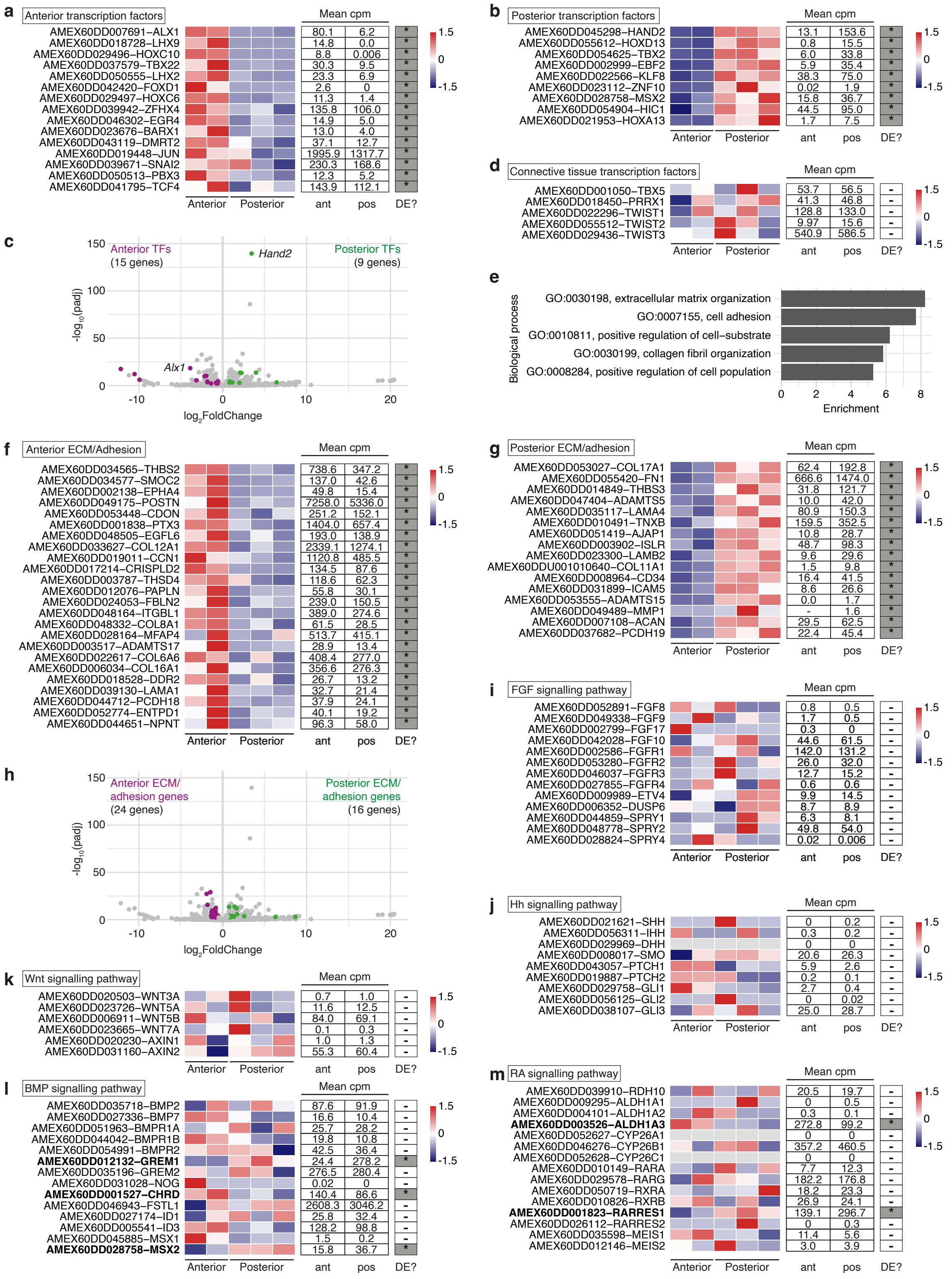


**Extended Data Fig. 4: Gene expression in anterior and posterior dermal cells.**
**a-b,** Heatmaps depicting the relative expression of transcription factors enriched in anterior cells (a) or posterior cells (b). Genes are ordered by decreasing statistical significance. Heatmap scores are normalised by row. cpm: counts per million. ant and pos: mean cpm in anterior or posterior samples respectively. An asterisk and dark grey box in the DE? column indicates differential expression. Gene nomenclature: AMEX[….] is the axolotl gene identifier and the gene symbol of the human homologue is given in suffix. **c,** Volcano plot highlighting the transcription factors differentially expressed in anterior cells (magenta) or in posterior cells (green). The most statistically significant transcription factor in each population is labelled. **d,** Heatmap of general connective tissue and forelimb identity genes. Depiction as in (a-b). **e,** Top 5 Gene Ontology (GO) terms describing the differentially expressed genes. **f-g,** Heatmaps depicting the relative expression of genes belonging to the GO term categories 'extracellular matrix' or 'cell adhesion'. Genes enriched in anterior cells (f) or posterior cells (g). Depiction as in (a-b). **h,** Volcano plot highlighting differentially expressed genes belonging to the GO term categories 'extracellular matrix' or 'cell adhesion'. **i-m,** Heatmaps for genes in selected functional categories. Heatmap depiction is as for (a-b), except that genes are not ordered by significance value. Bold gene names highlight genes with differential expression in anterior or posterior cells.

#### Extended Data Fig. 5: *Hand2* lineage tracing in embryo and mature limb.

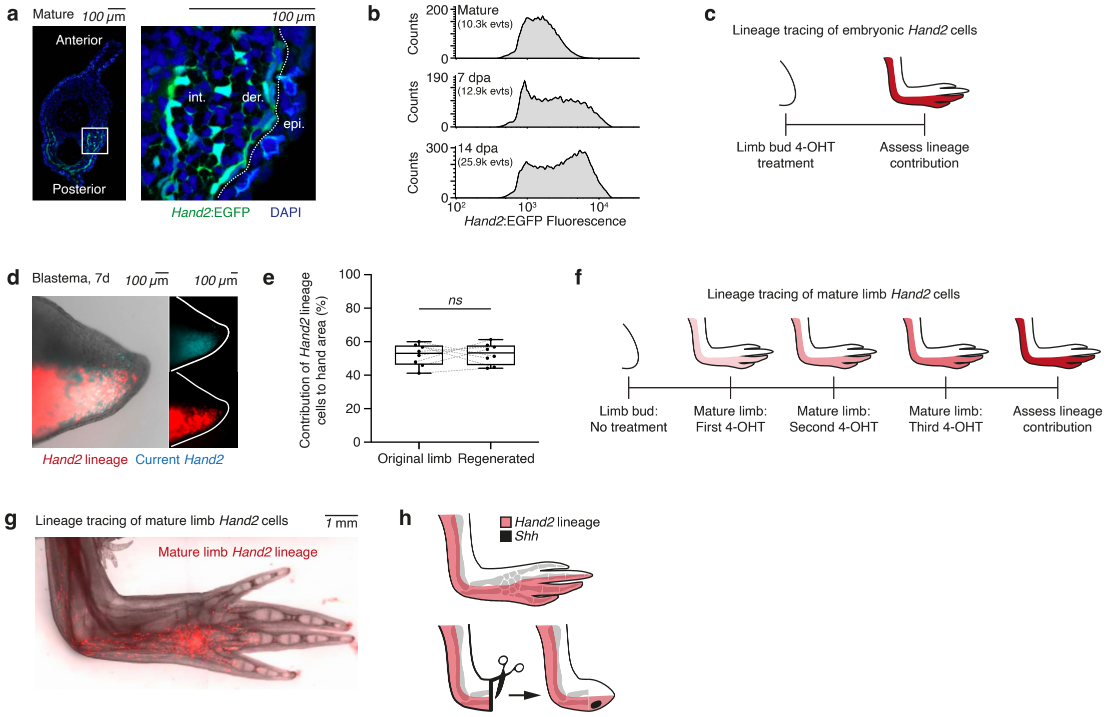


**Extended Data Fig. 5: *Hand2* lineage tracing in embryo and mature limb.**

**a,** Single transverse section through a Hand2:EGFP mature limb, with nuclei stained by DAPI (blue). The region in the white box is magnified in the panel to the right. Dotted line indicates the border between epidermis (epi.) and dermal cells (der.). Interstitial connective tissue cells (int.) reside deeper in the limb. Both dermal and interstitial cells in the posterior limb express Hand2:EGFP (green). **b,** Hand2:EGFP fluorescence intensity per cell in the mature limb (top), 7 dpa blastema (middle) and 14 dpa blastema (bottom), measured by flow cytometry. Fluorescence intensity is displayed in log scale. Only EGFP+ events (evts) are displayed. **c,** Strategy to lineage trace embryonic Hand2 cells into the mature limb. 4-OHT treatment was performed on Stage 42 embryos. **d,** 7 dpa blastema from a Hand2 lineage-traced limb. Regenerative Hand2 expression (cyan) and embryonic Hand2 lineage (red) overlap in widefield images. **e,** The area of the hand derived from embryonic Hand2 cells (mCherry+) was compared in the same limbs prior to amputation (original limb) and after regeneration (regenerated). Paired data (before and after amputation of the same limb) are connected by a dotted line. ns: not significant, p = 0.82, paired two-tailed t-test, n = 8 limbs per condition. **f,** Strategy to lineage trace Hand2 cells from the mature limb. 7 cm axolotls were treated three times with 4 uM 4-OHT at weekly intervals to induce labelling. **g,** Labelling of Hand2 cells after the treatment described in (f). **h,** Schematic depicting the contributions of Hand2 lineage cells to the mature limb and to the regenerative Shh signalling centre.

#### Extended Data Fig. 6: Characterisation of *Hand2* CRISPant phenotypes during development, regeneration and ALM.

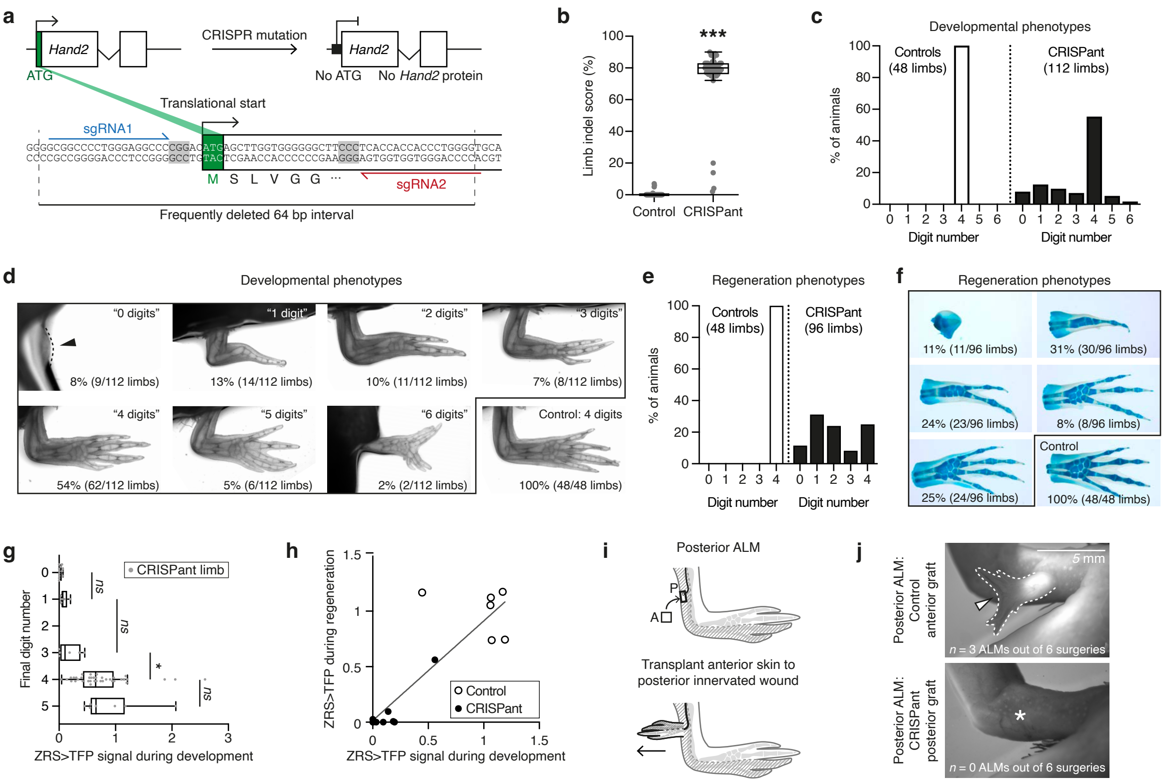

**Extended Data Fig. 6: Characterisation of *Hand2* CRISPant phenotypes during development, regeneration and ALM.**
**a,** Experimental strategy to mutate Hand2. Two sgRNAs targeting sequences flanking the Hand2 translational start codon (green) were co-injected with Cas9 protein into fertilised axolotl eggs. Shaded sequences highlight the PAM sequences for these sgRNAs. CRISPR/Cas9-mediated cutting removes the translational start from Hand2. **b,** Genomic DNA was extracted from control or Hand2 CRISPant larvae and sequenced to determine the frequency of indels. Each dot represents one limb. **c,** Bar chart summarising the range of digit number phenotypes observed in control or Hand2 CRISPant limbs at the end of development. **d,** Examples of each of the digit number phenotypes quantified in (c). **e,** Bar chart depicting digit numbers of amputated and regenerated control or Hand2 CRISPant limbs. **f,** Examples of each of the digit number phenotypes quantified in (e). Limbs were stained with Alcian Blue to highlight cartilage. **g,** Box plot comparing ZRS>TFP signal in Hand2 CRISPant limb buds (x-axis; '1' is the mean signal in control limb buds) with the eventual number of digits generated by the end of limb development (y-axis). Each dot represents 1 limb, n = 68. ns: not significant, $p > 0.05$. *: p = 2.34e-2, Kruskal-Wallis test followed by Dunn's post-hoc test. **h,** Correlation analysis between ZRS>TFP signal in the limb bud (x-axis) and in the blastema after amputation of the same limb (y-axis). Each dot represents one limb. There is a direct positive correlation between limb bud TFP and blastema TFP (Spearman's rank test, p = 2.40e-3, n = 14 limbs). **i,** Schematic depicting the posterior ALM. A patch of anterior limb skin transplanted to an innervated posterior wound elicits an accessory limb. **j,** Posterior ALM performed with control anterior skin (top) or Hand2 CRISPant posterior skin (bottom). 3 out of 6 control grafts elicited ALMs (arrowed). 0 out of 6 Hand2 CRISPant grafts elicited ALMs (asterisked).

#### Extended Data Fig. 7: Characterisation of limbs misexpressing *Hand2*.

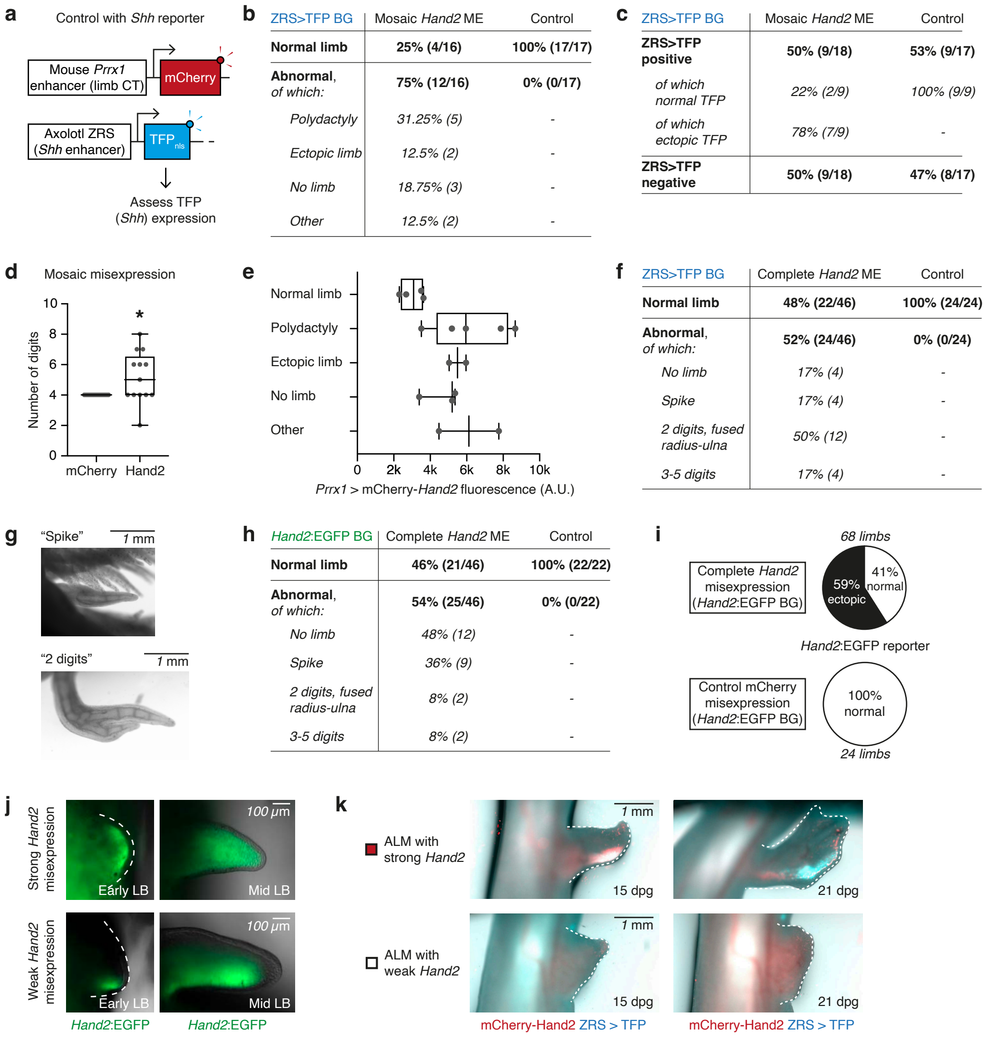


**b**

| ZRS>TFP BG | Mosaic *Hand2* ME | Control |
|---|---|---|
| **Normal limb** | **25% (4/16)** | **100% (17/17)** |
| **Abnormal**, *of which:* | **75% (12/16)** | **0% (0/17)** |
| *Polydactyly* | *31.25% (5)* | - |
| *Ectopic limb* | *12.5% (2)* | - |
| *No limb* | *18.75% (3)* | - |
| *Other* | *12.5% (2)* | - |

**c**

| ZRS>TFP BG | Mosaic *Hand2* ME | Control |
|---|---|---|
| **ZRS>TFP positive** | **50% (9/18)** | **53% (9/17)** |
| *of which normal TFP* | *22% (2/9)* | *100% (9/9)* |
| *of which ectopic TFP* | *78% (7/9)* | - |
| **ZRS>TFP negative** | **50% (9/18)** | **47% (8/17)** |

**f**

| ZRS>TFP BG | Complete *Hand2* ME | Control |
|---|---|---|
| **Normal limb** | **48% (22/46)** | **100% (24/24)** |
| **Abnormal**, *of which:* | **52% (24/46)** | **0% (0/24)** |
| *No limb* | *17% (4)* | - |
| *Spike* | *17% (4)* | - |
| *2 digits, fused radius-ulna* | *50% (12)* | - |
| *3-5 digits* | *17% (4)* | - |

**h**

| *Hand2*:EGFP BG | Complete *Hand2* ME | Control |
|---|---|---|
| **Normal limb** | **46% (21/46)** | **100% (22/22)** |
| **Abnormal**, *of which:* | **54% (25/46)** | **0% (0/22)** |
| *No limb* | *48% (12)* | - |
| *Spike* | *36% (9)* | - |
| *2 digits, fused radius-ulna* | *8% (2)* | - |
| *3-5 digits* | *8% (2)* | - |

**Extended Data Fig. 7: Characterisation of limbs misexpressing *Hand2*.**
**a,** Control axolotl for Hand2 misexpression experiments. **b,** Table summarising the phenotypes arising from mosaic Hand2 misexpression (ME) in the ZRS>TFP genetic background (BG). **c,** Table summarising the effects of mosaic Hand2 misexpression on ZRS>TFP expression. **d,** Box plot comparing the number of digits in mosaic Hand2 misexpression limbs (Hand2) with control limbs (mCherry). Each dot represents one limb. *: p = 0.03, Kolmogorov-Smirnov test, n = 17 (controls) and n = 13 (Hand2). **e,** Box plot comparing the level of mosaic mCherry-Hand2 misexpression (x-axis) with the resulting limb phenotype (y-axis). Each dot represents one limb, n = 16 in total. No stastistically significant differences were observed (Kruskal-Wallis test), likely due to the low n numbers in some categories (see (b)). **f,** Table summarising the effects of complete Hand2 misexpression on digit number (in the ZRS>TFP genetic background). **g,** Examples of the "spike" and "2 digits, fused radius-ulna" phenotypes quantified in (f). **h,** Table summarising the effects of complete Hand2 misexpression on digit number (in the Hand2:EGFP genetic background). **i,** Pie chart summarising the effect of complete Hand2 misexpression on Hand2:EGFP expression. **j,** Comparison of Hand2:EGFP expression (green) in axolotls misexpressing strong mCherry-Hand2 (top) or weak mCherry-Hand2 (bottom). **k,** Anterior ALM was performed with anteriorly located skin grafts misexpressing strong Hand2 (top) or weak Hand2 (bottom). All strong Hand2 ALMs induced ZRS>TFP (cyan, n = 8 out of 8 surgeries at 21 dpg) but weak Hand2 grafts did not (n = 0 out of 8 surgeries).

#### Extended Data Fig. 8: Characterisation of anterior-posterior positional memory switch.

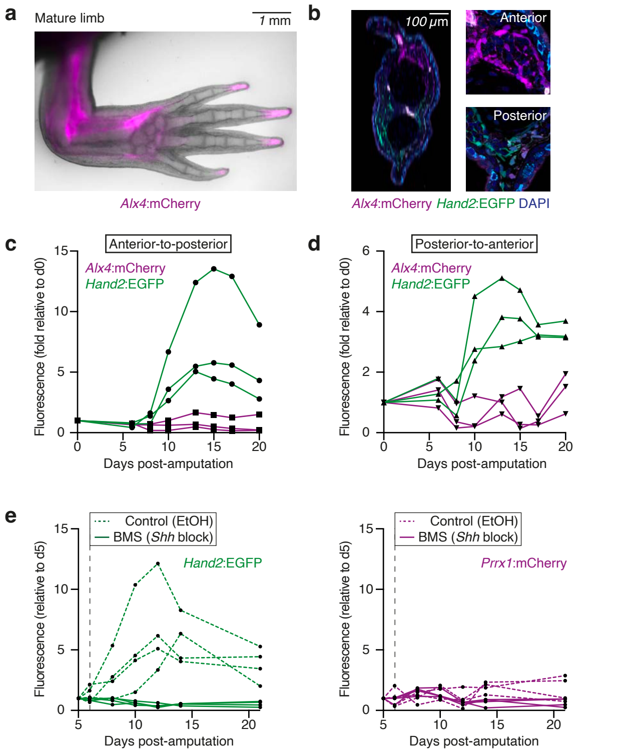


**Extended Data Fig. 8: Characterisation of anterior-posterior positional memory switch.**
**a,** Mature limb of an Alx4:mCherry_Hand2:EGFP double knock-in axolotl, depicting Alx4:mCherry fluorescence (magenta) only. **b,** Cross-section through the lower arm of an Alx4:mCherry_Hand2:EGFP mature limb, with nuclei stained using DAPI (blue). Insets show magnified regions of anterior and posterior tissue. **c,** Quantification of mCherry and EGFP fluorescence intensities in anterior-to-posterior transplantations after amputation. Intensities were measured in the blastema area and are normalised to the 0 dpa values. Each line represents data from 1 limb. n = 3. **d,** Quantification of mCherry and EGFP fluorescence intensities in posterior-to-anterior transplantations after amputation. Intensities were measured in the blastema area and are normalised to the 0 dpa values. Each line represents data from 1 limb. n = 3. **e,** Fluorescence intensity measurements of Hand2:EGFP (left) and Prrx1>mCherry (right) in anterior-to-posterior transplantations in the presence or absence of BMS-833923 (Shh pathway inhibitor). Animals were injected with one dose of BMS-833923 or ethanol (EtOH) at 6 dpa (dotted vertical line). Fluorescence intensities were measured in the blastema area and are normalised to the 5 dpa values. Each line represents data from 1 limb. n = 4 per condition.

**Extended Data Fig. 9: *Shh* signalling can induce a posterior memory in anterior blastema cells.**

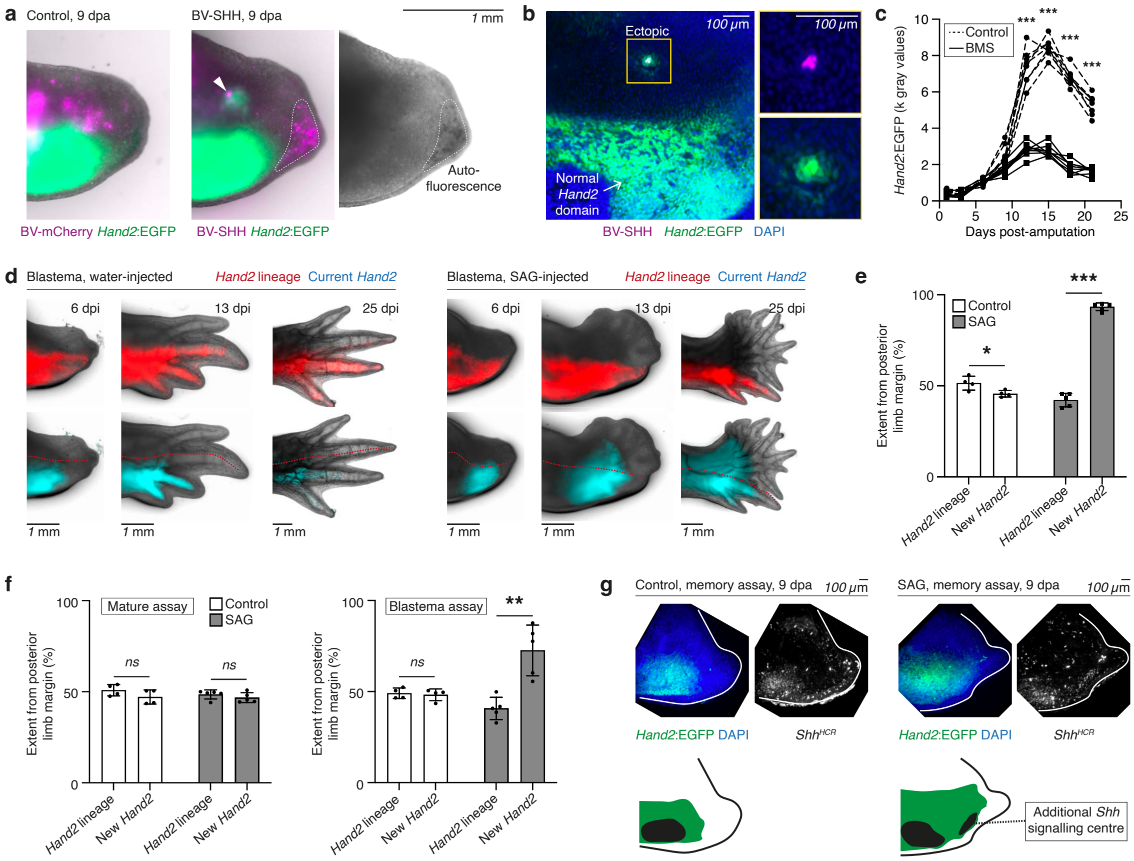


**Extended Data Fig. 9: *Shh* signalling can induce a posterior memory in anterior blastema cells.**
**a,** Mature limbs of Hand2:EGFP axolotls were injected with baculoviruses (BV) encoding Prrx1>mCherry (BV-mCherry, left) or Prrx1>mCherry-T2A-Shh (BV-Shh, right). 9 days after amputation, ectopic anterior Hand2:EGFP was observed around Shh-misexpressing cells (7 out of 10 blastemas). In the BV-SHH image, magenta signal in the distal epidermis is autofluorescence, as confirmed by chromatic densities in the brightfield image. **b,** Wholemount imaging of the BV-Shh blastema depicted in (a), stained for nuclei by DAPI (blue). Hand2:EGFP (green) is expressed in a ring around Shh-misexpressing cells (magenta). The yellow boxed region is magnified in the panels to the right. **c,** Hand2 expression during regeneration of axolotls injected with BMS or water (control). Mean EGFP fluorescence in the posterior blastema was measured from widefield images. There is no significant difference between BMS and control samples prior to 9 dpa. From 12 dpa, BMS-treated blastemas have significantly lower Hand2:EGFP fluorescence than controls. ***: $p < 1e-4$, 2-way ANOVA plus post-hoc testing, $F(7, 98) = 178.0$. n = 8 blastemas per condition. **d,** SAG treatment of Hand2 lineage traced animals. Images depict the blastema assay only, as the 'current Hand2' reporter (cyan) was not bright enough to observe in the mature assay. Regenerative Hand2 (cyan) is expressed within the posterior domain (red) in control blastemas (left). In SAG-treated blastemas (right), Hand2 becomes expressed in originally anterior cells (not red). **e,** Quantification of the experiment in (d). The relative extents of the posterior lineage (mCherry, 'Hand2 lineage') and regenerative Hand2 domain (EGFP, 'New Hand2') were measured from the posterior margin in 13 dpi blastemas. *: p = 4.42e-2, ***: p = 3.09e-8, 2-way ANOVA plus post-hoc testing, $F(1, 7) = 458.4$. n = 4 blastemas (control) or 5 blastemas (SAG). **f,** Bar charts summarising the results of the positional memory alteration assay. The percentage of the blastema positive for mCherry (Hand2 lineage) and EGFP (new Hand2) was quantified from the posterior edge. **: p = 5.60e-3, paired t-test, n = 5 per condition. **g,** SAG-treated animals form ectopic Shh signalling centres after amputation in 4 out of 6 cases. In this experiment, SAG was delivered by bathing regenerating axolotls continuously in 10 nM SAG for 21 days following amputation. After washout of the drug, limbs were re-amputated and Shh expression assessed in the new blastema by HCR.

#### Extended Data Fig. 10: Interpretation of positional memory alterations by *Shh* signalling.

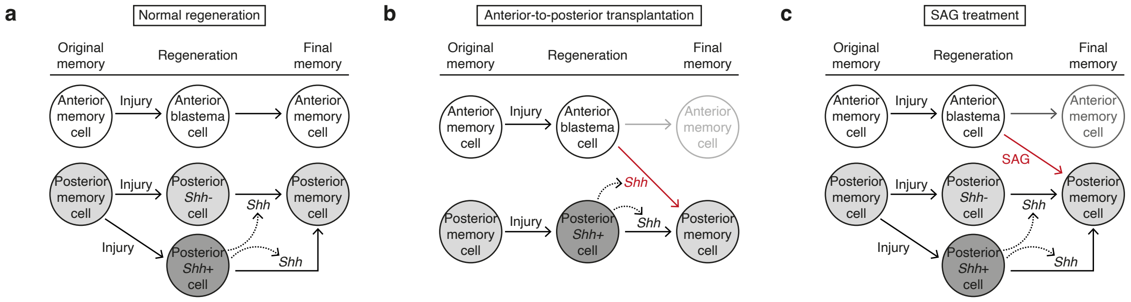


**Extended Data Fig. 10: Interpretation of positional memory alterations by *Shh* signalling.**
**a,** Broad concordance of original anterior/posterior memory and regenerated anterior/posterior memory during normal limb regeneration. It is important that Shh-expressing cells arise in the most posterior blastema to prevent undesired posteriorisation of anterior blastema cells (which are receptive to Shh signal). **b,** When anterior cells are transplanted to the posterior side, they come into the proximity of endogenous Shh-expressing posterior cells. Shh signalling posteriorises anterior blastema cells for the next round of regeneration (red arrow). **c,** When blastemas are treated with SAG, Shh signalling is activated in anterior blastema cells regardless of proximity to endogenous Shh-expressing posterior cells. These cells become posteriorised for the next round of regeneration (red arrow).
