## Extended Tables for "Molecular basis for positional memory and its reprogrammability in limb regeneration"

**Extended Data Table 1. Top Posterior Genes ordered by increasing p-value.**

| Gene ID and Symbol | baseMean | log2FoldChange | lfcSE | stat | pvalue | padj |
|---|---|---|---|---|---|---|
| AMEX60DD045298-HAND2 | 4912.45172 | 3.443650825 | 0.1345577 | 25.5923729 | 1.85E-144 | 2.94E-140 |
| AMEX60DD012132-GREM1 | 8540.91092 | 3.25094095 | 0.16105461 | 20.1853329 | 1.32E-90 | 1.05E-86 |
| AMEX60DD029166-EMP1 | 2292.65241 | 2.373880007 | 0.18425358 | 12.8837663 | 5.56E-38 | 2.94E-34 |
| AMEX60DD038184-KRT19 | 6905.60604 | 1.738820823 | 0.1546723 | 11.2419665 | 2.54E-29 | 5.03E-26 |
| AMEX60DD011094-TNFAIP2 | 1467.04171 | 2.146399177 | 0.20348551 | 10.5481674 | 5.18E-26 | 9.13E-23 |
| AMEX60DD029590-IGFBP5 | 17772.9286 | 0.994750364 | 0.10649902 | 9.34046525 | 9.59E-21 | 1.17E-17 |
| AMEX60DD032648 | 791.110044 | 4.352161621 | 0.47045477 | 9.25096718 | 2.22E-20 | 2.35E-17 |
| AMEX60DD024268-N/A | 168445.733 | 0.8466196 | 0.09465729 | 8.94405074 | 3.75E-19 | 3.13E-16 |
| AMEX60DD007136-CAPG | 19666.906 | 1.080803126 | 0.12251761 | 8.82161439 | 1.13E-18 | 8.95E-16 |
| AMEX60DD053027-COL17A1 | 7154.99358 | 1.425025615 | 0.16854798 | 8.45471779 | 2.80E-17 | 1.85E-14 |
| AMEX60DD055612-HOXD13 | 470.189221 | 3.95867925 | 0.47104571 | 8.40402349 | 4.31E-17 | 2.74E-14 |
| AMEX60DD054625-TBX2 | 1088.27978 | 2.088070732 | 0.24868584 | 8.39642002 | 4.60E-17 | 2.81E-14 |
| AMEX60DD055420-FN1 | 57753.2051 | 0.75250987 | 0.09000369 | 8.36087779 | 6.23E-17 | 3.66E-14 |
| AMEX60DD002999-EBF2 | 1155.34492 | 2.23459173 | 0.26894851 | 8.30862283 | 9.68E-17 | 5.49E-14 |
| AMEX60DD005352-ANXA2 | 17616.5041 | 1.0993795 | 0.13251059 | 8.29654086 | 1.07E-16 | 5.87E-14 |
| AMEX60DD049637-PRSS23 | 1977.57231 | 1.909873615 | 0.23254149 | 8.21304458 | 2.16E-16 | 1.14E-13 |
| AMEX60DD042697-EFNA5 | 7842.57822 | 1.185005537 | 0.14595543 | 8.11895493 | 4.70E-16 | 2.41E-13 |
| AMEX60DD003755-LOC102059433 | 92008.1817 | 0.921383041 | 0.11366008 | 8.10647875 | 5.21E-16 | 2.58E-13 |
| AMEX60DD001823-RARRES1 | 11906.6236 | 0.982061175 | 0.12133343 | 8.09390453 | 5.78E-16 | 2.78E-13 |
| AMEX60DD014849-THBS3 | 4237.65202 | 1.653942957 | 0.20915441 | 7.90776031 | 2.62E-15 | 1.09E-12 |
| AMEX60DD027994 | 113499.74 | 1.05339233 | 0.1346031 | 7.82591435 | 5.04E-15 | 2.05E-12 |
| AMEX60DD040289-ANGPT1 | 1774.05709 | 1.916553777 | 0.24740123 | 7.74674317 | 9.43E-15 | 3.74E-12 |
| AMEX60DD029823-KRT5 | 6762.94759 | 1.575341882 | 0.20539076 | 7.66997456 | 1.72E-14 | 6.66E-12 |
| AMEX60DD009815-VAT1 | 20353.9 | 0.920852663 | 0.12307946 | 7.4817736 | 7.33E-14 | 2.59E-11 |
| AMEX60DD008884-CYP2A13 | 391.923488 | 5.468803172 | 0.73686733 | 7.42169311 | 1.16E-13 | 3.82E-11 |
| AMEX60DD024244-MST1 | 5721.35214 | 1.086943706 | 0.14685424 | 7.40151375 | 1.35E-13 | 4.36E-11 |
| AMEX60DD051700-LOC113941412 | 9142.54214 | 0.90685493 | 0.12269317 | 7.391242 | 1.45E-13 | 4.62E-11 |
| AMEX60DD008883-CYP2A13 | 2647.77993 | 1.900175408 | 0.25803488 | 7.36402539 | 1.78E-13 | 5.55E-11 |
| AMEX60DD054004 | 676.216936 | 2.906676704 | 0.40708413 | 7.14023585 | 9.32E-13 | 2.64E-10 |
| AMEX60DD008885 | 303.142951 | 3.46334521 | 0.49137534 | 7.0482682 | 1.81E-12 | 4.96E-10 |
| AMEX60DD026748-MFAP5 | 52508.2854 | 1.044464086 | 0.14936079 | 6.99289331 | 2.69E-12 | 7.12E-10 |
| AMEX60DD014975-S100A10 | 21912.5135 | 0.729445133 | 0.10483862 | 6.95779023 | 3.46E-12 | 8.99E-10 |
| AMEX60DD018697-F13B | 511.524358 | 2.691494518 | 0.39525561 | 6.80950364 | 9.79E-12 | 2.47E-09 |
| AMEX60DD045295-SCRG1 | 1491.27378 | 1.563050079 | 0.23031511 | 6.7865719 | 1.15E-11 | 2.80E-09 |
| AMEX60DD011307-FBLN5 | 5318.32318 | 0.96420854 | 0.14351243 | 6.71864137 | 1.83E-11 | 4.41E-09 |
| AMEX60DD024084-SEMA3G | 3682.07295 | 1.517732593 | 0.23918877 | 6.34533372 | 2.22E-10 | 4.89E-08 |
| AMEX60DD036862-PRSS23 | 5859.64087 | 1.220094313 | 0.19286452 | 6.3261729 | 2.51E-10 | 5.46E-08 |
| AMEX60DD049867-SFRP2 | 3185.2241 | 1.089299519 | 0.17556608 | 6.2044986 | 5.49E-10 | 1.18E-07 |
| AMEX60DD026746-A2ML1 | 399.218006 | 2.673499615 | 0.4401543 | 6.07400545 | 1.25E-09 | 2.57E-07 |
| AMEX60DD008878-TAFA2 | 459.617751 | 2.850442859 | 0.47641067 | 5.98316337 | 2.19E-09 | 4.34E-07 |
| AMEX60DD052276-LOC112113698 | 831.697183 | 3.308852757 | 0.55646381 | 5.94621374 | 2.74E-09 | 5.38E-07 |
| AMEX60DD041315 | 5275.55951 | 0.840379842 | 0.14339704 | 5.86051037 | 4.61E-09 | 8.72E-07 |
| AMEX60DD007579 | 1366.51285 | 1.512658467 | 0.25862411 | 5.84886885 | 4.95E-09 | 9.13E-07 |
| AMEX60DD028151 | 23.8815995 | 20.44548793 | 3.51407136 | 5.81817665 | 5.95E-09 | 1.07E-06 |
| AMEX60DD056319-PRKAG3 | 178.029898 | 4.333120688 | 0.74908398 | 5.78455926 | 7.27E-09 | 1.25E-06 |
| AMEX60DD047404-ADAMTS5 | 1426.64432 | 1.72120583 | 0.29923357 | 5.75204791 | 8.82E-09 | 1.47E-06 |
| AMEX60DD047262-LOC115092556 | 22.6394389 | 20.3694219 | 3.5461032 | 5.74417064 | 9.24E-09 | 1.53E-06 |
| AMEX60DD003298 | 21.7283406 | 20.31478881 | 3.58273938 | 5.67018324 | 1.43E-08 | 2.31E-06 |
| AMEX60DD043733-DR999-PMT147[illegible] | 4857.66764 | 0.799258154 | 0.14122918 | 5.65929891 | 1.52E-08 | 2.44E-06 |
| AMEX60DD005684-UMOD | 4678.88763 | 1.14571914 | 0.20355439 | 5.62856526 | 1.82E-08 | 2.88E-06 |
| AMEX60DD056780-LOC113575404 | 21.1410881 | 20.27658353 | 3.60347276 | 5.62695624 | 1.83E-08 | 2.88E-06 |
| AMEX60DDU001005517 | 20.7495865 | 20.24363125 | 3.61781963 | 5.59553359 | 2.20E-08 | 3.39E-06 |
| AMEX60DD037078-FGF16 | 897.00511 | 1.524575089 | 0.27258547 | 5.59301674 | 2.23E-08 | 3.41E-06 |

| | | | | | | |
|---|---|---|---|---|---|---|
| AMEX60DD014985-LOC115077151 | 533.277855 | 2.262911089 | 0.40543602 | 5.58142582 | 2.39E-08 | 3.54E-06 |
| AMEX60DD020691-XB5721655.S | 20.2180507 | 20.21282943 | 3.6314951 | 5.56598009 | 2.61E-08 | 3.80E-06 |
| AMEX60DD007948-GCC1 | 26.0348585 | 19.03906012 | 3.45451454 | 5.51135621 | 3.56E-08 | 5.09E-06 |
| AMEX60DD043026-CFC1 | 18.8661991 | 20.11911469 | 3.68696606 | 5.45682123 | 4.85E-08 | 6.75E-06 |
| AMEX60DD034915 | 18.6077635 | 20.10024268 | 3.69802416 | 5.43540059 | 5.47E-08 | 7.48E-06 |
| AMEX60DD016049-LGI1 | 27.4907473 | 18.51978609 | 3.41251927 | 5.42701291 | 5.73E-08 | 7.76E-06 |
| AMEX60DD017064-LOC102363937 | 2582.01916 | 1.41888493 | 0.26156575 | 5.42458236 | 5.81E-08 | 7.76E-06 |
| AMEX60DD035993-KCNG3 | 955.30407 | 1.787752341 | 0.33011777 | 5.41549869 | 6.11E-08 | 8.08E-06 |
| AMEX60DD036742-ZNF658 | 18.0710011 | 19.95575475 | 3.72245686 | 5.36090961 | 8.28E-08 | 1.09E-05 |
| AMEX60DD010473-PSMB7 | 2914.49491 | 0.848630583 | 0.16064405 | 5.28267687 | 1.27E-07 | 1.62E-05 |
| AMEX60DD012162-XELAEV-180399 | 2321.51545 | 0.967789936 | 0.18583962 | 5.20766209 | 1.91E-07 | 2.33E-05 |
| AMEX60DD016367-LOC115080300 | 3085.24811 | 1.114752851 | 0.21425951 | 5.20281633 | 1.96E-07 | 2.38E-05 |
| AMEX60DD037492-GPC4 | 17754.2114 | 0.641214652 | 0.12330135 | 5.20038633 | 1.99E-07 | 2.39E-05 |
| AMEX60DD035117-LAMA4 | 6227.24234 | 0.653641319 | 0.12664745 | 5.16110928 | 2.45E-07 | 2.91E-05 |
| AMEX60DD024521-CYP2F1 | 8848.63873 | 0.884755836 | 0.17153739 | 5.15780179 | 2.50E-07 | 2.94E-05 |
| AMEX60DD053266-RGS10 | 1594.04541 | 1.549312447 | 0.30269826 | 5.1183394 | 3.08E-07 | 3.57E-05 |
| AMEX60DD000084-ADGRD1 | 2128.2854 | 1.720598828 | 0.3364647 | 5.11375735 | 3.16E-07 | 3.63E-05 |
| AMEX60DD004644-FUT3 | 708.766354 | 1.361808709 | 0.26728308 | 5.09500534 | 3.49E-07 | 3.98E-05 |
| AMEX60DD011997-CRIP1 | 8382.10982 | 1.087300684 | 0.2135917 | 5.09055682 | 3.57E-07 | 4.05E-05 |
| AMEX60DD022566-KLF8 | 3091.12695 | 0.829169326 | 0.16373253 | 5.0641698 | 4.10E-07 | 4.52E-05 |
| AMEX60DD049312-SHISA2 | 282.551001 | 2.676956723 | 0.52909383 | 5.05951224 | 4.20E-07 | 4.60E-05 |
| AMEX60DD023968 | 1171.25506 | 2.25543714 | 0.4484712 | 5.02916824 | 4.93E-07 | 5.35E-05 |
| AMEX60DD012216-CFL2 | 4150.24472 | 0.872454606 | 0.17366677 | 5.023728 | 5.07E-07 | 5.47E-05 |
| AMEX60DD024630 | 18797.8784 | 0.764373742 | 0.15314661 | 4.99112424 | 6.00E-07 | 6.27E-05 |
| AMEX60DD025131-CD248 | 27581.3376 | 0.672160279 | 0.13515564 | 4.9732316 | 6.58E-07 | 6.74E-05 |
| AMEX60DD038192-KRT19 | 269.742062 | 2.343686642 | 0.47703343 | 4.91304483 | 8.97E-07 | 8.95E-05 |
| AMEX60DD020532-COL21A1 | 5635.87481 | 1.103256949 | 0.22609583 | 4.87959888 | 1.06E-06 | 0.0001035 |
| AMEX60DD020053-ELOVL1 | 24724.5452 | 0.646217362 | 0.13246828 | 4.87827998 | 1.07E-06 | 0.00010356 |
| AMEX60DD010491-TNXB | 14399.8852 | 1.025301077 | 0.21312368 | 4.81082658 | 1.50E-06 | 0.00014369 |
| AMEX60DD025178-CAPN1 | 24099.4861 | 0.617751977 | 0.12844025 | 4.80964469 | 1.51E-06 | 0.00014369 |
| AMEX60DD051419-AJAP1 | 1082.95382 | 1.098173741 | 0.22843592 | 4.80736022 | 1.53E-06 | 0.00014448 |
| AMEX60DD012436 | 50.9836263 | 8.730046356 | 1.82071981 | 4.7948324 | 1.63E-06 | 0.0001529 |
| AMEX60DD010096-KRT17 | 58.0803328 | 8.918620481 | 1.86295816 | 4.78734343 | 1.69E-06 | 0.00015686 |
| AMEX60DD003902-ISLR | 3994.21959 | 0.772821134 | 0.1623958 | 4.75887383 | 1.95E-06 | 0.00017913 |
| AMEX60DD008914-WNT2B | 656.307747 | 1.341689731 | 0.2819699 | 4.75827287 | 1.95E-06 | 0.00017913 |
| AMEX60DD022420-NEBL | 601.81601 | 1.821173488 | 0.38369077 | 4.74646161 | 2.07E-06 | 0.00018882 |
| AMEX60DD006238-SEMA3E | 1343.18817 | 1.466383395 | 0.31004872 | 4.72952566 | 2.25E-06 | 0.0002041 |
| AMEX60DD031445 | 66.6121528 | 9.116311274 | 1.93321357 | 4.71562553 | 2.41E-06 | 0.0002173 |
| AMEX60DD052522-LIPA | 4000.27662 | 0.769597516 | 0.1635635 | 4.70519121 | 2.54E-06 | 0.00022742 |
| AMEX60DD032718-SCARA5 | 5418.09963 | 0.883727043 | 0.18798859 | 4.70096107 | 2.59E-06 | 0.00023088 |
| AMEX60DD023300-LAMB2 | 1063.00385 | 1.22505409 | 0.26074329 | 4.69831489 | 2.62E-06 | 0.00023258 |
| AMEX60DDU001010640-COL11A1 | 326.320566 | 2.5619916 | 0.54737983 | 4.68046401 | 2.86E-06 | 0.00024824 |
| AMEX60DD035574-AKAP12 | 28280.9047 | 0.629671183 | 0.13541942 | 4.64978482 | 3.32E-06 | 0.00028353 |
| AMEX60DD012996-ARHGEF15 | 1221.81176 | 1.238904835 | 0.26843367 | 4.61531092 | 3.93E-06 | 0.00032615 |
| AMEX60DD008964-CD34 | 1568.35589 | 1.032026831 | 0.22443686 | 4.59829481 | 4.26E-06 | 0.00035211 |
| AMEX60DD001909-IGSF10 | 12699.3477 | 0.657207551 | 0.14337059 | 4.58397737 | 4.56E-06 | 0.00037131 |
| AMEX60DD036951-ARHGAP20 | 565.684265 | 1.643421253 | 0.3588472 | 4.57972435 | 4.66E-06 | 0.00037701 |
| AMEX60DD046915 | 777.451906 | 1.293241916 | 0.28315516 | 4.56725543 | 4.94E-06 | 0.0003961 |
| AMEX60DD045638-UCHL1 | 2562.83405 | 1.038253685 | 0.22746932 | 4.564368 | 5.01E-06 | 0.00039957 |
| AMEX60DD010804-TAPBP | 3616.98147 | 0.639671154 | 0.14178038 | 4.5117042 | 6.43E-06 | 0.00050032 |
| AMEX60DD025422-AHNAK | 4013.32081 | 1.068788787 | 0.23795461 | 4.49156588 | 7.07E-06 | 0.00053947 |
| AMEX60DD025264-LOC106737871 | 165.445742 | 2.463044341 | 0.54915548 | 4.48514933 | 7.29E-06 | 0.00055331 |
| AMEX60DD023112-ZNF10 | 55.1665625 | 6.42850121 | 1.43641481 | 4.47537936 | 7.63E-06 | 0.00057646 |
| AMEX60DD028758-MSX2 | 1445.19736 | 1.158942658 | 0.26074724 | 4.44469765 | 8.80E-06 | 0.00065891 |
| AMEX60DD030050-TIMD4 | 2203.52501 | 1.184338252 | 0.26699838 | 4.43575068 | 9.18E-06 | 0.00068366 |
| AMEX60DD002023-PROS1 | 294.523367 | 2.445814515 | 0.55492966 | 4.40743159 | 1.05E-05 | 0.00077217 |

| | | | | | | |
|---|---|---|---|---|---|---|
| AMEX60DD031899-ICAM5 | 976.15079 | 1.352357644 | 0.30739378 | 4.39943072 | 1.09E-05 | 0.00079748 |
| AMEX60DDU001010337 | 41.202502 | 8.42307732 | 1.91651608 | 4.39499433 | 1.11E-05 | 0.00079994 |
| AMEX60DD008880-CYP2A6 | 77.7925056 | 7.879075857 | 1.79423343 | 4.39133265 | 1.13E-05 | 0.00080905 |
| AMEX60DD054904-HIC1 | 3755.37755 | 0.821676996 | 0.18721581 | 4.38892947 | 1.14E-05 | 0.00081435 |
| AMEX60DDU001006983-PARPI-001 | 2806.23633 | 0.748824389 | 0.17475163 | 4.2850782 | 1.83E-05 | 0.00125508 |
| AMEX60DD053555-ADAMTS15 | 49.7214696 | 6.279766739 | 1.48093962 | 4.24039349 | 2.23E-05 | 0.00152641 |
| AMEX60DD047987-SEPTIN10 | 4136.03006 | 0.595833858 | 0.14072707 | 4.23396751 | 2.30E-05 | 0.00154409 |
| AMEX60DD015002-S100A13 | 8207.27349 | 0.846534637 | 0.201839 | 4.19410844 | 2.74E-05 | 0.00181869 |
| AMEX60DD049489-MMP1 | 48.8004415 | 8.667396854 | 2.06802382 | 4.19114944 | 2.78E-05 | 0.00182021 |
| AMEX60DD021953-HOXA13 | 264.245694 | 1.994431727 | 0.4787357 | 4.1660393 | 3.10E-05 | 0.00200296 |
| AMEX60DD007108-ACAN | 2490.26505 | 0.994854433 | 0.23882297 | 4.1656564 | 3.10E-05 | 0.00200296 |
| AMEX60DD007343-TNKS1BP1 | 13478.3831 | 0.609428576 | 0.1463731 | 4.16352843 | 3.13E-05 | 0.00201354 |
| AMEX60DD004446-MUC5B | 7922.47533 | 2.407111469 | 0.57845508 | 4.16127642 | 3.16E-05 | 0.00201901 |
| AMEX60DD000921-OASL | 1914.17122 | 0.834003289 | 0.20043006 | 4.1610689 | 3.17E-05 | 0.00201901 |
| AMEX60DD031199-ABCA8 | 3706.2058 | 0.722546324 | 0.1749887 | 4.12910272 | 3.64E-05 | 0.00228456 |
| AMEX60DD012693-B3GNT9 | 2661.34638 | 0.882018331 | 0.21402135 | 4.12116986 | 3.77E-05 | 0.00235537 |
| AMEX60DD032906-CYP2J2 | 474.771301 | 1.453293329 | 0.35412864 | 4.10385714 | 4.06E-05 | 0.00251902 |
| AMEX60DD020640-IL4R | 2432.8198 | 0.739630972 | 0.18028007 | 4.10267742 | 4.08E-05 | 0.00252205 |
| AMEX60DD023548-CAV3 | 47.9478423 | 7.204512668 | 1.75664354 | 4.1012946 | 4.11E-05 | 0.00252733 |
| AMEX60DD029914-S100A11 | 781.158493 | 1.066999091 | 0.26182257 | 4.07527546 | 4.60E-05 | 0.00278408 |
| AMEX60DD029423-ZNF300 | 139.611398 | 2.597669876 | 0.63842764 | 4.06885558 | 4.72E-05 | 0.00285102 |
| AMEX60DD004552-SIGIRR | 2974.73032 | 0.895319438 | 0.2201321 | 4.06719156 | 4.76E-05 | 0.00286058 |
| AMEX60DD030522-ARHGDIB | 4581.52691 | 0.654441791 | 0.16153756 | 4.05132883 | 5.09E-05 | 0.00302723 |
| AMEX60DD001008-DTX1 | 64.0917588 | 5.072671872 | 1.2543175 | 4.04416896 | 5.25E-05 | 0.00308656 |
| AMEX60DD045942-AFAP1 | 1114.49433 | 1.549570254 | 0.38350951 | 4.04050021 | 5.33E-05 | 0.00312368 |
| AMEX60DD048288-F10 | 12126.2856 | 0.665655896 | 0.16509803 | 4.03188282 | 5.53E-05 | 0.00322857 |
| AMEX60DD037682-PCDH19 | 1854.51964 | 0.808726282 | 0.20248724 | 3.99396176 | 6.50E-05 | 0.00371902 |
| AMEX60DD010594-RGS3 | 62.7874553 | 4.346138459 | 1.09325296 | 3.9754189 | 7.03E-05 | 0.00398223 |
| AMEX60DD031409-C1QTNF1 | 494.890427 | 1.675935524 | 0.4293424 | 3.90349408 | 9.48E-05 | 0.00525589 |
| AMEX60DD028768-DRD1 | 105.655615 | 2.722356246 | 0.69878281 | 3.89585461 | 9.79E-05 | 0.00537379 |
| AMEX60DD022344-TAC1 | 49.7464985 | 5.288965856 | 1.35900782 | 3.8917847 | 9.95E-05 | 0.00544592 |
| AMEX60DD024603-SERTAD2 | 147.926746 | 2.30865129 | 0.59411531 | 3.88586401 | 0.00010197 | 0.00556121 |
| AMEX60DD026007-LOC115075658 | 4757.3707 | 0.648159288 | 0.16718047 | 3.87700373 | 0.00010575 | 0.00569481 |
| AMEX60DD047972-SCML2 | 122.263557 | 2.934457254 | 0.75830794 | 3.86974353 | 0.00010895 | 0.00582203 |
| AMEX60DDU001002672 | 32.7048533 | 8.090173738 | 2.09664326 | 3.8586315 | 0.00011402 | 0.00603749 |
| AMEX60DD034470-LOC101955705 | 4041.81194 | 0.752363311 | 0.19502928 | 3.8576941 | 0.00011446 | 0.00603749 |
| AMEX60DD004436-AP2A2 | 3856.30948 | 0.651074689 | 0.16877692 | 3.85760509 | 0.0001145 | 0.00603749 |
| AMEX60DD023020-PLA2G4C | 2119.15825 | 0.771228386 | 0.20006778 | 3.85483547 | 0.00011581 | 0.00608603 |
| AMEX60DD052038-CAPNS1 | 1766.21324 | 0.75208767 | 0.19525992 | 3.8517258 | 0.00011729 | 0.0061233 |
| AMEX60DD008176-CPM | 148.784668 | 2.266421047 | 0.59040313 | 3.83876868 | 0.00012365 | 0.00643441 |
| AMEX60DD005280-HARBI1 | 41.2018022 | 6.986080626 | 1.82265038 | 3.83292413 | 0.00012663 | 0.00656774 |
| AMEX60DD038318-DSP | 8704.70013 | 0.714319097 | 0.18712398 | 3.81735733 | 0.00013489 | 0.00692822 |
| AMEX60DD056032-STEAP3 | 1678.83628 | 0.729781976 | 0.19131897 | 3.81447779 | 0.00013647 | 0.0069421 |
| AMEX60DD041516-MTARC1 | 840.322064 | 1.129760047 | 0.29651488 | 3.81012941 | 0.00013889 | 0.00703671 |
| AMEX60DD026180-ASIC1 | 361.085135 | 1.815753594 | 0.4766316 | 3.80955356 | 0.00013922 | 0.00703671 |
| AMEX60DD039930-PI15 | 47.9763601 | 7.174200464 | 1.91003773 | 3.7560517 | 0.00017262 | 0.00853646 |
| AMEX60DD047161-PCP4 | 182.024808 | 3.207392341 | 0.85393963 | 3.75599426 | 0.00017266 | 0.00853646 |
| AMEX60DD034453-MRPS5 | 108.380525 | 2.748004099 | 0.73626123 | 3.7323765 | 0.00018968 | 0.00932024 |
| AMEX60DD013815-HSD11B1L | 144.580045 | 2.678793596 | 0.72068683 | 3.71700088 | 0.0002016 | 0.00973257 |
| AMEX60DD007394-CTSS | 1193.77061 | 1.760717315 | 0.47386671 | 3.71563832 | 0.00020269 | 0.00973257 |
| AMEX60DD042726 | 33.3604272 | 8.117815684 | 2.1854298 | 3.71451679 | 0.00020359 | 0.00973257 |

**Extended Data Table 2. Top Posterior Genes ordered by decreasing fold change over anterior.**

| Gene ID and Symbol | baseMean | log2FoldChange | lfcSE | stat | pvalue | padj |
|---|---|---|---|---|---|---|
| AMEX60DD028151 | 23.8815995 | 20.44548793 | 3.51407136 | 5.81817665 | 5.95E-09 | 1.07E-06 |
| AMEX60DD047262-LOC115092556 | 22.6394389 | 20.3694219 | 3.5461032 | 5.74417064 | 9.24E-09 | 1.53E-06 |
| AMEX60DD003298 | 21.7283406 | 20.31478881 | 3.58273938 | 5.67018324 | 1.43E-08 | 2.31E-06 |
| AMEX60DD056780-LOC113575404 | 21.1410881 | 20.27658353 | 3.60347276 | 5.62695624 | 1.83E-08 | 2.88E-06 |
| AMEX60DDU001005517 | 20.7495865 | 20.24363125 | 3.61781963 | 5.59553359 | 2.20E-08 | 3.39E-06 |
| AMEX60DD020691-XB5721655.S | 20.2180507 | 20.21282943 | 3.6314951 | 5.56598009 | 2.61E-08 | 3.80E-06 |
| AMEX60DD043026-CFC1 | 18.8661991 | 20.11911469 | 3.68696606 | 5.45682123 | 4.85E-08 | 6.75E-06 |
| AMEX60DD034915 | 18.6077635 | 20.10024268 | 3.69802416 | 5.43540059 | 5.47E-08 | 7.48E-06 |
| AMEX60DD036742-ZNF658 | 18.0710011 | 19.95575475 | 3.72245686 | 5.36090961 | 8.28E-08 | 1.09E-05 |
| AMEX60DD007948-GCC1 | 26.0348585 | 19.03906012 | 3.45451454 | 5.51135621 | 3.56E-08 | 5.09E-06 |
| AMEX60DD016049-LGI1 | 27.4907473 | 18.51978609 | 3.41251927 | 5.42701291 | 5.73E-08 | 7.76E-06 |
| AMEX60DD031445 | 66.6121528 | 9.116311274 | 1.93321357 | 4.71562553 | 2.41E-06 | 0.0002173 |
| AMEX60DD010096-KRT17 | 58.0803328 | 8.918620481 | 1.86295816 | 4.78734343 | 1.69E-06 | 0.00015686 |
| AMEX60DD012436 | 50.9836263 | 8.730046356 | 1.82071981 | 4.7948324 | 1.63E-06 | 0.0001529 |
| AMEX60DD049489-MMP1 | 48.8004415 | 8.667396854 | 2.06802382 | 4.19114944 | 2.78E-05 | 0.00182021 |
| AMEX60DDU001010337 | 41.202502 | 8.42307732 | 1.91651608 | 4.39499433 | 1.11E-05 | 0.00079994 |
| AMEX60DD042726 | 33.3604272 | 8.117815684 | 2.1854298 | 3.71451679 | 0.00020359 | 0.00973257 |
| AMEX60DDU001002672 | 32.7048533 | 8.090173738 | 2.09664326 | 3.8586315 | 0.00011402 | 0.00603749 |
| AMEX60DD008880-CYP2A6 | 77.7925056 | 7.879075857 | 1.79423343 | 4.39133265 | 1.13E-05 | 0.00080905 |
| AMEX60DD023548-CAV3 | 47.9478423 | 7.204512668 | 1.75664354 | 4.1012946 | 4.11E-05 | 0.00252733 |
| AMEX60DD039930-PI15 | 47.9763601 | 7.174200464 | 1.91003773 | 3.7560517 | 0.00017262 | 0.00853646 |
| AMEX60DD005280-HARBI1 | 41.2018022 | 6.986080626 | 1.82265038 | 3.83292413 | 0.00012663 | 0.00656774 |
| AMEX60DD023112-ZNF10 | 55.1665625 | 6.42850121 | 1.43641481 | 4.47537936 | 7.63E-06 | 0.00057646 |
| AMEX60DD053555-ADAMTS15 | 49.7214696 | 6.279766739 | 1.48093962 | 4.24039349 | 2.23E-05 | 0.00152641 |
| AMEX60DD008884-CYP2A13 | 391.923488 | 5.468803172 | 0.73686733 | 7.42169311 | 1.16E-13 | 3.82E-11 |
| AMEX60DD022344-TAC1 | 49.7464985 | 5.288965856 | 1.35900782 | 3.8917847 | 9.95E-05 | 0.00544592 |
| AMEX60DD001008-DTX1 | 64.0917588 | 5.072671872 | 1.2543175 | 4.04416896 | 5.25E-05 | 0.00308656 |
| AMEX60DD032648 | 791.110044 | 4.352161621 | 0.47045477 | 9.25096718 | 2.22E-20 | 2.35E-17 |
| AMEX60DD010594-RGS3 | 62.7874553 | 4.346138459 | 1.09325296 | 3.9754189 | 7.03E-05 | 0.00398223 |
| AMEX60DD056319-PRKAG3 | 178.029898 | 4.333120688 | 0.74908398 | 5.78455926 | 7.27E-09 | 1.25E-06 |
| AMEX60DD055612-HOXD13 | 470.189221 | 3.95867925 | 0.47104571 | 8.40402349 | 4.31E-17 | 2.74E-14 |
| AMEX60DD008885 | 303.142951 | 3.46334521 | 0.49137534 | 7.0482682 | 1.81E-12 | 4.96E-10 |
| AMEX60DD045298-HAND2 | 4912.45172 | 3.443650825 | 0.1345577 | 25.5923729 | 1.85E-144 | 2.94E-140 |
| AMEX60DD052276-LOC112113698 | 831.697183 | 3.308852757 | 0.55646381 | 5.94621374 | 2.74E-09 | 5.38E-07 |
| AMEX60DD012132-GREM1 | 8540.91092 | 3.25094095 | 0.16105461 | 20.1853329 | 1.32E-90 | 1.05E-86 |
| AMEX60DD047161-PCP4 | 182.024808 | 3.207392341 | 0.85393963 | 3.75599426 | 0.00017266 | 0.00853646 |
| AMEX60DD047972-SCML2 | 122.263557 | 2.934457254 | 0.75830794 | 3.86974353 | 0.00010895 | 0.00582203 |
| AMEX60DD054004 | 676.216936 | 2.906676704 | 0.40708413 | 7.14023585 | 9.32E-13 | 2.64E-10 |
| AMEX60DD008878-TAFA2 | 459.617751 | 2.850442859 | 0.47641067 | 5.98316337 | 2.19E-09 | 4.34E-07 |
| AMEX60DD034453-MRPS5 | 108.380525 | 2.748004099 | 0.73626123 | 3.7323765 | 0.00018968 | 0.00932024 |
| AMEX60DD028768-DRD1 | 105.655615 | 2.722356246 | 0.69878281 | 3.89585461 | 9.79E-05 | 0.00537379 |
| AMEX60DD018697-F13B | 511.524358 | 2.691494518 | 0.39525561 | 6.80950364 | 9.79E-12 | 2.47E-09 |
| AMEX60DD013815-HSD11B1L | 144.580045 | 2.678793596 | 0.72068683 | 3.71700088 | 0.0002016 | 0.00973257 |
| AMEX60DD049312-SHISA2 | 282.551001 | 2.676956723 | 0.52909383 | 5.05951224 | 4.20E-07 | 4.60E-05 |
| AMEX60DD026746-A2ML1 | 399.218006 | 2.673499615 | 0.4401543 | 6.07400545 | 1.25E-09 | 2.57E-07 |
| AMEX60DD029423-ZNF300 | 139.611398 | 2.597669876 | 0.63842764 | 4.06885558 | 4.72E-05 | 0.00285102 |
| AMEX60DDU001010640-COL11A1 | 326.320566 | 2.5619916 | 0.54737983 | 4.68046401 | 2.86E-06 | 0.00024824 |
| AMEX60DD025264-LOC106737871 | 165.445742 | 2.463044341 | 0.54915548 | 4.48514933 | 7.29E-06 | 0.00055331 |
| AMEX60DD002023-PROS1 | 294.523367 | 2.445814515 | 0.55492966 | 4.40743159 | 1.05E-05 | 0.00077217 |
| AMEX60DD004446-MUC5B | 7922.47533 | 2.407111469 | 0.57845508 | 4.16127642 | 3.16E-05 | 0.00201901 |
| AMEX60DD029166-EMP1 | 2292.65241 | 2.373880007 | 0.18425358 | 12.8837663 | 5.56E-38 | 2.94E-34 |
| AMEX60DD038192-KRT19 | 269.742062 | 2.343686642 | 0.47703343 | 4.91304483 | 8.97E-07 | 8.95E-05 |
| AMEX60DD024603-SERTAD2 | 147.926746 | 2.30865129 | 0.59411531 | 3.88586401 | 0.00010197 | 0.00556121 |

| | | | | | | |
|---|---|---|---|---|---|---|
| AMEX60DD008176-CPM | 148.784668 | 2.266421047 | 0.59040313 | 3.83876868 | 0.00012365 | 0.00643441 |
| AMEX60DD014985-LOC115077151 | 533.277855 | 2.262911089 | 0.40543602 | 5.58142582 | 2.39E-08 | 3.54E-06 |
| AMEX60DD023968 | 1171.25506 | 2.25543714 | 0.4484712 | 5.02916824 | 4.93E-07 | 5.35E-05 |
| AMEX60DD002999-EBF2 | 1155.34492 | 2.23459173 | 0.26894851 | 8.30862283 | 9.68E-17 | 5.49E-14 |
| AMEX60DD011094-TNFAIP2 | 1467.04171 | 2.146399177 | 0.20348551 | 10.5481674 | 5.18E-26 | 9.13E-23 |
| AMEX60DD054625-TBX2 | 1088.27978 | 2.088070732 | 0.24868584 | 8.39642002 | 4.60E-17 | 2.81E-14 |
| AMEX60DD021953-HOXA13 | 264.245694 | 1.994431727 | 0.4787357 | 4.1660393 | 3.10E-05 | 0.00200296 |
| AMEX60DD040289-ANGPT1 | 1774.05709 | 1.916553777 | 0.24740123 | 7.74674317 | 9.43E-15 | 3.74E-12 |
| AMEX60DD049637-PRSS23 | 1977.57231 | 1.909873615 | 0.23254149 | 8.21304458 | 2.16E-16 | 1.14E-13 |
| AMEX60DD008883-CYP2A13 | 2647.77993 | 1.900175408 | 0.25803488 | 7.36402539 | 1.78E-13 | 5.55E-11 |
| AMEX60DD022420-NEBL | 601.81601 | 1.821173488 | 0.38369077 | 4.74646161 | 2.07E-06 | 0.00018882 |
| AMEX60DD026180-ASIC1 | 361.085135 | 1.815753594 | 0.4766316 | 3.80955356 | 0.00013922 | 0.00703671 |
| AMEX60DD035993-KCNG3 | 955.30407 | 1.787752341 | 0.33011777 | 5.41549869 | 6.11E-08 | 8.08E-06 |
| AMEX60DD007394-CTSS | 1193.77061 | 1.760717315 | 0.47386671 | 3.71563832 | 0.00020269 | 0.00973257 |
| AMEX60DD038184-KRT19 | 6905.60604 | 1.738820823 | 0.1546723 | 11.2419665 | 2.54E-29 | 5.03E-26 |
| AMEX60DD047404-ADAMTS5 | 1426.64432 | 1.72120583 | 0.29923357 | 5.75204791 | 8.82E-09 | 1.47E-06 |
| AMEX60DD000084-ADGRD1 | 2128.2854 | 1.720598828 | 0.3364647 | 5.11375735 | 3.16E-07 | 3.63E-05 |
| AMEX60DD031409-C1QTNF1 | 494.890427 | 1.675935524 | 0.4293424 | 3.90349408 | 9.48E-05 | 0.00525589 |
| AMEX60DD014849-THBS3 | 4237.65202 | 1.653942957 | 0.20915441 | 7.90776031 | 2.62E-15 | 1.09E-12 |
| AMEX60DD036951-ARHGAP20 | 565.684265 | 1.643421253 | 0.3588472 | 4.57972435 | 4.66E-06 | 0.00037701 |
| AMEX60DD029823-KRT5 | 6762.94759 | 1.575341882 | 0.20539076 | 7.66997456 | 1.72E-14 | 6.66E-12 |
| AMEX60DD045295-SCRG1 | 1491.27378 | 1.563050079 | 0.23031511 | 6.7865719 | 1.15E-11 | 2.80E-09 |
| AMEX60DD045942-AFAP1 | 1114.49433 | 1.549570254 | 0.38350951 | 4.04050021 | 5.33E-05 | 0.00312368 |
| AMEX60DD053266-RGS10 | 1594.04541 | 1.549312447 | 0.30269826 | 5.1183394 | 3.08E-07 | 3.57E-05 |
| AMEX60DD037078-FGF16 | 897.00511 | 1.524575089 | 0.27258547 | 5.59301674 | 2.23E-08 | 3.41E-06 |
| AMEX60DD024084-SEMA3G | 3682.07295 | 1.517732593 | 0.23918877 | 6.34533372 | 2.22E-10 | 4.89E-08 |
| AMEX60DD007579 | 1366.51285 | 1.512658467 | 0.25862411 | 5.84886885 | 4.95E-09 | 9.13E-07 |
| AMEX60DD006238-SEMA3E | 1343.18817 | 1.466383395 | 0.31004872 | 4.72952566 | 2.25E-06 | 0.0002041 |
| AMEX60DD032906-CYP2J2 | 474.771301 | 1.453293329 | 0.35412864 | 4.10385714 | 4.06E-05 | 0.00251902 |
| AMEX60DD053027-COL17A1 | 7154.99358 | 1.425025615 | 0.16854798 | 8.45471779 | 2.80E-17 | 1.85E-14 |
| AMEX60DD017064-LOC102363937 | 2582.01916 | 1.41888493 | 0.26156575 | 5.42458236 | 5.81E-08 | 7.76E-06 |
| AMEX60DD004644-FUT3 | 708.766354 | 1.361808709 | 0.26728308 | 5.09500534 | 3.49E-07 | 3.98E-05 |
| AMEX60DD031899-ICAM5 | 976.15079 | 1.352357644 | 0.30739378 | 4.39943072 | 1.09E-05 | 0.00079748 |
| AMEX60DD008914-WNT2B | 656.307747 | 1.341689731 | 0.2819699 | 4.75827287 | 1.95E-06 | 0.00017913 |
| AMEX60DD046915 | 777.451906 | 1.293241916 | 0.28315516 | 4.56725543 | 4.94E-06 | 0.0003961 |
| AMEX60DD012996-ARHGEF15 | 1221.81176 | 1.238904835 | 0.26843367 | 4.61531092 | 3.93E-06 | 0.00032615 |
| AMEX60DD023300-LAMB2 | 1063.00385 | 1.22505409 | 0.26074329 | 4.69831489 | 2.62E-06 | 0.00023258 |
| AMEX60DD036862-PRSS23 | 5859.64087 | 1.220094313 | 0.19286452 | 6.3261729 | 2.51E-10 | 5.46E-08 |
| AMEX60DD042697-EFNA5 | 7842.57822 | 1.185005537 | 0.14595543 | 8.11895493 | 4.70E-16 | 2.41E-13 |
| AMEX60DD030050-TIMD4 | 2203.52501 | 1.184338252 | 0.26699838 | 4.43575068 | 9.18E-06 | 0.00068366 |
| AMEX60DD028758-MSX2 | 1445.19736 | 1.158942658 | 0.26074724 | 4.44469765 | 8.80E-06 | 0.00065891 |
| AMEX60DD005684-UMOD | 4678.88763 | 1.14571914 | 0.20355439 | 5.62856526 | 1.82E-08 | 2.88E-06 |
| AMEX60DD041516-MTARC1 | 840.322064 | 1.129760047 | 0.29651488 | 3.81012941 | 0.00013889 | 0.00703671 |
| AMEX60DD016367-LOC115080300 | 3085.24811 | 1.114752851 | 0.21425951 | 5.20281633 | 1.96E-07 | 2.38E-05 |
| AMEX60DD020532-COL21A1 | 5635.87481 | 1.103256949 | 0.22609583 | 4.87959888 | 1.06E-06 | 0.0001035 |
| AMEX60DD005352-ANXA2 | 17616.5041 | 1.0993795 | 0.13251059 | 8.29654086 | 1.07E-16 | 5.87E-14 |
| AMEX60DD051419-AJAP1 | 1082.95382 | 1.098173741 | 0.22843592 | 4.80736022 | 1.53E-06 | 0.00014448 |
| AMEX60DD049867-SFRP2 | 3185.2241 | 1.089299519 | 0.17556608 | 6.2044986 | 5.49E-10 | 1.18E-07 |
| AMEX60DD011997-CRIP1 | 8382.10982 | 1.087300684 | 0.2135917 | 5.09055682 | 3.57E-07 | 4.05E-05 |
| AMEX60DD024244-MST1 | 5721.35214 | 1.086943706 | 0.14685424 | 7.40151375 | 1.35E-13 | 4.36E-11 |
| AMEX60DD007136-CAPG | 19666.906 | 1.080803126 | 0.12251761 | 8.82161439 | 1.13E-18 | 8.95E-16 |
| AMEX60DD025422-AHNAK | 4013.32081 | 1.068788787 | 0.23795461 | 4.49156588 | 7.07E-06 | 0.00053947 |
| AMEX60DD029914-S100A11 | 781.158493 | 1.066999091 | 0.26182257 | 4.07527546 | 4.60E-05 | 0.00278408 |
| AMEX60DD027994 | 113499.74 | 1.05339233 | 0.1346031 | 7.82591435 | 5.04E-15 | 2.05E-12 |
| AMEX60DD026748-MFAP5 | 52508.2854 | 1.044464086 | 0.14936079 | 6.99289331 | 2.69E-12 | 7.12E-10 |
| AMEX60DD045638-UCHL1 | 2562.83405 | 1.038253685 | 0.22746932 | 4.564368 | 5.01E-06 | 0.00039957 |

| | | | | | | |
|---|---|---|---|---|---|---|
| AMEX60DD008964-CD34 | 1568.35589 | 1.032026831 | 0.22443686 | 4.59829481 | 4.26E-06 | 0.00035211 |
| AMEX60DD010491-TNXB | 14399.8852 | 1.025301077 | 0.21312368 | 4.81082658 | 1.50E-06 | 0.00014369 |
| AMEX60DD007108-ACAN | 2490.26505 | 0.994854433 | 0.23882297 | 4.1656564 | 3.10E-05 | 0.00200296 |
| AMEX60DD029590-IGFBP5 | 17772.9286 | 0.994750364 | 0.10649902 | 9.34046525 | 9.59E-21 | 1.17E-17 |
| AMEX60DD001823-RARRES1 | 11906.6236 | 0.982061175 | 0.12133343 | 8.09390453 | 5.78E-16 | 2.78E-13 |
| AMEX60DD012162-XELAEV-1803999 | 2321.51545 | 0.967789936 | 0.18583962 | 5.20766209 | 1.91E-07 | 2.33E-05 |
| AMEX60DD011307-FBLN5 | 5318.32318 | 0.96420854 | 0.14351243 | 6.71864137 | 1.83E-11 | 4.41E-09 |
| AMEX60DD003755-LOC102059433 | 92008.1817 | 0.921383041 | 0.11366008 | 8.10647875 | 5.21E-16 | 2.58E-13 |
| AMEX60DD009815-VAT1 | 20353.9 | 0.920852663 | 0.12307946 | 7.4817736 | 7.33E-14 | 2.59E-11 |
| AMEX60DD051700-LOC113941412 | 9142.54214 | 0.90685493 | 0.12269317 | 7.391242 | 1.45E-13 | 4.62E-11 |
| AMEX60DD004552-SIGIRR | 2974.73032 | 0.895319438 | 0.2201321 | 4.06719156 | 4.76E-05 | 0.00286058 |
| AMEX60DD024521-CYP2F1 | 8848.63873 | 0.884755836 | 0.17153739 | 5.15780179 | 2.50E-07 | 2.94E-05 |
| AMEX60DD032718-SCARA5 | 5418.09963 | 0.883727043 | 0.18798859 | 4.70096107 | 2.59E-06 | 0.00023088 |
| AMEX60DD012693-B3GNT9 | 2661.34638 | 0.882018331 | 0.21402135 | 4.12116986 | 3.77E-05 | 0.00235537 |
| AMEX60DD012216-CFL2 | 4150.24472 | 0.872454606 | 0.17366677 | 5.023728 | 5.07E-07 | 5.47E-05 |
| AMEX60DD010473-PSMB7 | 2914.49491 | 0.848630583 | 0.16064405 | 5.28267687 | 1.27E-07 | 1.62E-05 |
| AMEX60DD024268-N/A | 168445.733 | 0.8466196 | 0.09465729 | 8.94405074 | 3.75E-19 | 3.13E-16 |
| AMEX60DD015002-S100A13 | 8207.27349 | 0.846534637 | 0.201839 | 4.19410844 | 2.74E-05 | 0.00181869 |
| AMEX60DD041315 | 5275.55951 | 0.840379842 | 0.14339704 | 5.86051037 | 4.61E-09 | 8.72E-07 |
| AMEX60DD000921-OASL | 1914.17122 | 0.834003289 | 0.20043006 | 4.1610689 | 3.17E-05 | 0.00201901 |
| AMEX60DD022566-KLF8 | 3091.12695 | 0.829169326 | 0.16373253 | 5.0641698 | 4.10E-07 | 4.52E-05 |
| AMEX60DD054904-HIC1 | 3755.37755 | 0.821676996 | 0.18721581 | 4.38892947 | 1.14E-05 | 0.00081435 |
| AMEX60DD037682-PCDH19 | 1854.51964 | 0.808726282 | 0.20248724 | 3.99396176 | 6.50E-05 | 0.00371902 |
| AMEX60DD043733-DR999-PMT1478 | 4857.66764 | 0.799258154 | 0.14122918 | 5.65929891 | 1.52E-08 | 2.44E-06 |
| AMEX60DD003902-ISLR | 3994.21959 | 0.772821134 | 0.1623958 | 4.75887383 | 1.95E-06 | 0.00017913 |
| AMEX60DD023020-PLA2G4C | 2119.15825 | 0.771228386 | 0.20006778 | 3.85483547 | 0.00011581 | 0.00608603 |
| AMEX60DD052522-LIPA | 4000.27662 | 0.769597516 | 0.1635635 | 4.70519121 | 2.54E-06 | 0.00022742 |
| AMEX60DD024630 | 18797.8784 | 0.764373742 | 0.15314661 | 4.99112424 | 6.00E-07 | 6.27E-05 |
| AMEX60DD055420-FN1 | 57753.2051 | 0.75250987 | 0.09000369 | 8.36087779 | 6.23E-17 | 3.66E-14 |
| AMEX60DD034470-LOC101955705 | 4041.81194 | 0.752363311 | 0.19502928 | 3.8576941 | 0.00011446 | 0.00603749 |
| AMEX60DD052038-CAPNS1 | 1766.21324 | 0.75208767 | 0.19525992 | 3.8517258 | 0.00011729 | 0.0061233 |
| AMEX60DDU001006983-PARPI-0014 | 2806.23633 | 0.748824389 | 0.17475163 | 4.2850782 | 1.83E-05 | 0.00125508 |
| AMEX60DD020640-IL4R | 2432.8198 | 0.739630972 | 0.18028007 | 4.10267742 | 4.08E-05 | 0.00252205 |
| AMEX60DD056032-STEAP3 | 1678.83628 | 0.729781976 | 0.19131897 | 3.81447779 | 0.00013647 | 0.0069421 |
| AMEX60DD014975-S100A10 | 21912.5135 | 0.729445133 | 0.10483862 | 6.95779023 | 3.46E-12 | 8.99E-10 |
| AMEX60DD031199-ABCA8 | 3706.2058 | 0.722546324 | 0.1749887 | 4.12910272 | 3.64E-05 | 0.00228456 |
| AMEX60DD038318-DSP | 8704.70013 | 0.714319097 | 0.18712398 | 3.81735733 | 0.00013489 | 0.00692822 |
| AMEX60DD025131-CD248 | 27581.3376 | 0.672160279 | 0.13515564 | 4.9732316 | 6.58E-07 | 6.74E-05 |
| AMEX60DD048288-F10 | 12126.2856 | 0.665655896 | 0.16509803 | 4.03188282 | 5.53E-05 | 0.00322857 |
| AMEX60DD001909-IGSF10 | 12699.3477 | 0.657207551 | 0.14337059 | 4.58397737 | 4.56E-06 | 0.00037131 |
| AMEX60DD030522-ARHGDIB | 4581.52691 | 0.654441791 | 0.16153756 | 4.05132883 | 5.09E-05 | 0.00302723 |
| AMEX60DD035117-LAMA4 | 6227.24234 | 0.653641319 | 0.12664745 | 5.16110928 | 2.45E-07 | 2.91E-05 |
| AMEX60DD004436-AP2A2 | 3856.30948 | 0.651074689 | 0.16877692 | 3.85760509 | 0.0001145 | 0.00603749 |
| AMEX60DD026007-LOC115075658 | 4757.3707 | 0.648159288 | 0.16718047 | 3.87700373 | 0.00010575 | 0.00569481 |
| AMEX60DD020053-ELOVL1 | 24724.5452 | 0.646217362 | 0.13246828 | 4.87827998 | 1.07E-06 | 0.00010356 |
| AMEX60DD037492-GPC4 | 17754.2114 | 0.641214652 | 0.12330135 | 5.20038633 | 1.99E-07 | 2.39E-05 |
| AMEX60DD010804-TAPBP | 3616.98147 | 0.639671154 | 0.14178038 | 4.5117042 | 6.43E-06 | 0.00050032 |
| AMEX60DD035574-AKAP12 | 28280.9047 | 0.629671183 | 0.13541942 | 4.64978482 | 3.32E-06 | 0.00028353 |
| AMEX60DD025178-CAPN1 | 24099.4861 | 0.617751977 | 0.12844025 | 4.80964469 | 1.51E-06 | 0.00014369 |
| AMEX60DD007343-TNKS1BP1 | 13478.3831 | 0.609428576 | 0.1463731 | 4.16352843 | 3.13E-05 | 0.00201354 |
| AMEX60DD047987-SEPTIN10 | 4136.03006 | 0.595833858 | 0.14072707 | 4.23396751 | 2.30E-05 | 0.00154409 |

**Extended Data Table 3. Top Anterior Genes ordered by increasing p-value.**

| Gene ID and Symbol | baseMean | log2FoldChange | lfcSE | stat | pvalue | padj |
|---|---|---|---|---|---|---|
| AMEX60DD040394-ENPP2 | 4769.4996 | -2.507817369 | 0.19723487 | -12.714879 | 4.89E-37 | 1.94E-33 |
| AMEX60DD034565-THBS2 | 27875.7261 | -1.471111044 | 0.12247635 | -12.011388 | 3.10E-33 | 9.83E-30 |
| AMEX60DD034577-SMOC2 | 4361.98956 | -1.922237066 | 0.16503035 | -11.647779 | 2.36E-31 | 6.23E-28 |
| AMEX60DD045837-C1QTNF7 | 2086.61487 | -3.058125714 | 0.26442664 | -11.56512 | 6.19E-31 | 1.40E-27 |
| AMEX60DD014035-CPAMD8 | 15019.7225 | -1.237477495 | 0.11861955 | -10.432324 | 1.77E-25 | 2.80E-22 |
| AMEX60DD007691-ALX1 | 1951.28752 | -3.879454882 | 0.40006001 | -9.6971823 | 3.10E-22 | 4.47E-19 |
| AMEX60DD018728-LHX9 | 310.656084 | -12.15935289 | 1.2850391 | -9.4622435 | 3.01E-21 | 3.99E-18 |
| AMEX60DD014152-CRLF1 | 4087.29505 | -1.237293188 | 0.1332404 | -9.2861712 | 1.60E-20 | 1.81E-17 |
| AMEX60DD019499-PLPP3 | 18098.7828 | -1.31367337 | 0.14406571 | -9.1185707 | 7.61E-20 | 7.55E-17 |
| AMEX60DD003526-ALDH1A3 | 8943.15973 | -1.604185396 | 0.17837179 | -8.993493 | 2.39E-19 | 2.24E-16 |
| AMEX60DD002138-EPHA4 | 1559.31708 | -1.822645064 | 0.20313111 | -8.972752 | 2.89E-19 | 2.55E-16 |
| AMEX60DD018273-PRDX6 | 34403.2575 | -1.573974711 | 0.17982707 | -8.7527127 | 2.08E-18 | 1.57E-15 |
| AMEX60DD017769-SLC6A2 | 1062.99576 | -2.838572615 | 0.33145193 | -8.5640553 | 1.09E-17 | 7.86E-15 |
| AMEX60DD019770-LOC115099908 | 1390.74803 | -2.210020809 | 0.25941041 | -8.5193991 | 1.60E-17 | 1.11E-14 |
| AMEX60DD049175-POSTN | 328298.719 | -0.828029956 | 0.10267285 | -8.0647409 | 7.34E-16 | 3.43E-13 |
| AMEX60DD005869-RSPO1 | 9175.68926 | -1.487338743 | 0.18557085 | -8.0149375 | 1.10E-15 | 5.00E-13 |
| AMEX60DD029496-HOXC10 | 193.701276 | -10.51461044 | 1.32334737 | -7.9454652 | 1.93E-15 | 8.53E-13 |
| AMEX60DD053448-CDON | 10079.3829 | -0.848501068 | 0.10704207 | -7.9267996 | 2.25E-15 | 9.65E-13 |
| AMEX60DD036390-MDGA1 | 1168.69929 | -2.027373767 | 0.26660386 | -7.6044426 | 2.86E-14 | 1.08E-11 |
| AMEX60DD015718-CRABP2 | 16671.4165 | -1.104369399 | 0.145927 | -7.5679578 | 3.79E-14 | 1.40E-11 |
| AMEX60DD048418-LOC115076842 | 158.065401 | -11.18475886 | 1.48058405 | -7.5542883 | 4.21E-14 | 1.52E-11 |
| AMEX60DD000888-MN1 | 761.370421 | -2.026388821 | 0.27135362 | -7.4677052 | 8.16E-14 | 2.82E-11 |
| AMEX60DD030673-NTN1 | 4615.27492 | -1.214084683 | 0.1635444 | -7.4235786 | 1.14E-13 | 3.82E-11 |
| AMEX60DD037579-TBX22 | 981.691962 | -1.940600339 | 0.26462578 | -7.3333759 | 2.24E-13 | 6.85E-11 |
| AMEX60DD050555-LHX2 | 763.615828 | -2.165270948 | 0.30083957 | -7.1974273 | 6.14E-13 | 1.84E-10 |
| AMEX60DD001838-PTX3 | 50987.6973 | -1.248917624 | 0.17394136 | -7.1801074 | 6.97E-13 | 2.05E-10 |
| AMEX60DD052816-CRTAC1 | 4611.44867 | -1.15509378 | 0.16171993 | -7.1425568 | 9.16E-13 | 2.64E-10 |
| AMEX60DD046939-RABL3 | 2967.96611 | -1.087722657 | 0.15241696 | -7.1364934 | 9.57E-13 | 2.67E-10 |
| AMEX60DD048505-EGFL6 | 8627.65393 | -0.864325484 | 0.12270735 | -7.0437955 | 1.87E-12 | 5.03E-10 |
| AMEX60DD036008-PKDCC | 2041.34381 | -1.283665815 | 0.18735552 | -6.8514973 | 7.31E-12 | 1.87E-09 |
| AMEX60DD033627-COL12A1 | 90200.8947 | -1.114222582 | 0.16412379 | -6.7889159 | 1.13E-11 | 2.80E-09 |
| AMEX60DD019011-CCN1 | 38850.2766 | -1.33115672 | 0.20137848 | -6.6102232 | 3.84E-11 | 9.09E-09 |
| AMEX60DD017214-CRISPLD2 | 5668.74564 | -0.926187592 | 0.14267374 | -6.4916472 | 8.49E-11 | 1.98E-08 |
| AMEX60DD001004-EMID1 | 7308.09216 | -1.264861261 | 0.19686039 | -6.4251689 | 1.32E-10 | 3.03E-08 |
| AMEX60DD035799-ISM1 | 9466.91442 | -0.779163331 | 0.1217125 | -6.4016706 | 1.54E-10 | 3.48E-08 |
| AMEX60DD001527-CHRD | 5748.75414 | -0.918562679 | 0.14438121 | -6.3620652 | 1.99E-10 | 4.45E-08 |
| AMEX60DD029025-CLK4 | 3622.54533 | -0.971096464 | 0.15803843 | -6.1446855 | 8.01E-10 | 1.70E-07 |
| AMEX60DD028693-VSTM2L | 2602.59124 | -1.884959595 | 0.30977347 | -6.0849614 | 1.17E-09 | 2.43E-07 |
| AMEX60DD013358-PER1 | 389.257763 | -2.233125923 | 0.36879543 | -6.0551888 | 1.40E-09 | 2.85E-07 |
| AMEX60DD007033-FAM180A | 2308.91906 | -1.322991273 | 0.22096547 | -5.9873215 | 2.13E-09 | 4.29E-07 |
| AMEX60DD035184-SRD5A2 | 5667.73939 | -1.143950113 | 0.19300763 | -5.9269685 | 3.09E-09 | 5.97E-07 |
| AMEX60DD018240-PAPPA2 | 532.556007 | -3.694423657 | 0.6252155 | -5.9090404 | 3.44E-09 | 6.58E-07 |
| AMEX60DD042420-FOXD1 | 65.8278902 | -9.920178428 | 1.69602903 | -5.8490617 | 4.94E-09 | 9.13E-07 |
| AMEX60DD046940-HGD | 13281.7243 | -0.821919948 | 0.14089492 | -5.833567 | 5.43E-09 | 9.90E-07 |
| AMEX60DD003787-THSD4 | 4528.18517 | -1.156400564 | 0.19913241 | -5.807194 | 6.35E-09 | 1.13E-06 |
| AMEX60DD043210-PCSK5 | 4999.43042 | -0.881173123 | 0.15183104 | -5.8036428 | 6.49E-09 | 1.14E-06 |
| AMEX60DD012076-PAPLN | 2140.7147 | -1.015133212 | 0.17516707 | -5.7952286 | 6.82E-09 | 1.19E-06 |
| AMEX60DD002008-PID1 | 3766.15586 | -1.03417846 | 0.17899906 | -5.7775636 | 7.58E-09 | 1.29E-06 |
| AMEX60DD042580-NR2F1 | 866.38496 | -1.615003099 | 0.27963255 | -5.7754475 | 7.67E-09 | 1.30E-06 |
| AMEX60DD024053-FBLN2 | 9726.5494 | -0.793794924 | 0.13937634 | -5.6953349 | 1.23E-08 | 2.01E-06 |
| AMEX60DDU001005418-SEMA3F | 6149.82378 | -0.712649484 | 0.12687116 | -5.6171116 | 1.94E-08 | 3.02E-06 |
| AMEX60DD029497-HOXC6 | 304.845403 | -3.133002508 | 0.5607923 | -5.5867431 | 2.31E-08 | 3.50E-06 |

| | | | | | | |
|---|---|---|---|---|---|---|
| AMEX60DD028577 | 1321.32344 | -1.262353305 | 0.2260491 | -5.5844209 | 2.34E-08 | 3.51E-06 |
| AMEX60DD048164-ITGBL1 | 17043.675 | -0.800340444 | 0.14372803 | -5.5684369 | 2.57E-08 | 3.78E-06 |
| AMEX60DD035020-SLC16A10 | 1933.92486 | -0.987392117 | 0.1789146 | -5.51879 | 3.41E-08 | 4.92E-06 |
| AMEX60DD050768-KRT18 | 48.5623484 | -9.481845879 | 1.72734717 | -5.4892531 | 4.04E-08 | 5.72E-06 |
| AMEX60DD017420-CD82 | 3323.35702 | -0.804100117 | 0.14682791 | -5.4764802 | 4.34E-08 | 6.09E-06 |
| AMEX60DD013889-OCLN | 3555.05707 | -1.315658782 | 0.24180464 | -5.440999 | 5.30E-08 | 7.31E-06 |
| AMEX60DD000424-E1301-TTI00391 | 142.185063 | -3.420087065 | 0.63052619 | -5.4241792 | 5.82E-08 | 7.76E-06 |
| AMEX60DD043399-BNC2 | 2331.18423 | -1.044435736 | 0.19505282 | -5.3546304 | 8.57E-08 | 1.12E-05 |
| AMEX60DD021193-CHST12 | 644.718904 | -1.569407653 | 0.29570127 | -5.3074092 | 1.11E-07 | 1.43E-05 |
| AMEX60DD051000-LMO4 | 5762.20934 | -1.171535729 | 0.22139688 | -5.2915638 | 1.21E-07 | 1.55E-05 |
| AMEX60DD027622-R3HDML | 1110.89122 | -1.206491732 | 0.2284927 | -5.28022 | 1.29E-07 | 1.63E-05 |
| AMEX60DD015688-LRRC71 | 43.6647423 | -9.328383525 | 1.77900959 | -5.2435825 | 1.57E-07 | 1.94E-05 |
| AMEX60DD043069-DAPK1 | 1267.48303 | -1.814804171 | 0.35242194 | -5.1495209 | 2.61E-07 | 3.05E-05 |
| AMEX60DD033076-GDF6 | 451.802642 | -1.690135526 | 0.33233548 | -5.0856308 | 3.66E-07 | 4.12E-05 |
| AMEX60DD039942-ZFHX4 | 6203.76701 | -0.590178723 | 0.11607079 | -5.0846445 | 3.68E-07 | 4.12E-05 |
| AMEX60DD045225-CPE | 940.961677 | -1.238757914 | 0.24386402 | -5.0797075 | 3.78E-07 | 4.20E-05 |
| AMEX60DD003916-SEMA7A | 39.9199504 | -9.199190674 | 1.8333513 | -5.0176912 | 5.23E-07 | 5.57E-05 |
| AMEX60DD048332-COL8A1 | 2203.62174 | -1.379899396 | 0.27529397 | -5.0124578 | 5.37E-07 | 5.69E-05 |
| AMEX60DD024186-GNAI2 | 5249.54012 | -0.689900313 | 0.13811002 | -4.9952953 | 5.87E-07 | 6.17E-05 |
| AMEX60DD016542-NTN1 | 9019.96556 | -1.143902186 | 0.2300058 | -4.9733624 | 6.58E-07 | 6.74E-05 |
| AMEX60DD022752-ZNF385D | 2930.91358 | -0.828870742 | 0.16666848 | -4.9731703 | 6.59E-07 | 6.74E-05 |
| AMEX60DD028164-MFAP4 | 23910.4664 | -0.59668289 | 0.12008112 | -4.9689984 | 6.73E-07 | 6.85E-05 |
| AMEX60DD046302-EGR4 | 488.999052 | -1.833383622 | 0.37090205 | -4.9430399 | 7.69E-07 | 7.78E-05 |
| AMEX60DD007681-NTS | 400.055312 | -3.250143164 | 0.65830161 | -4.9371642 | 7.93E-07 | 7.96E-05 |
| AMEX60DD045587-CORIN | 3252.86717 | -0.851020218 | 0.17393292 | -4.8928071 | 9.94E-07 | 9.86E-05 |
| AMEX60DD018972-AK5 | 4221.21616 | -1.184540473 | 0.24242398 | -4.8862348 | 1.03E-06 | 0.00010132 |
| AMEX60DD007057-LOC101478214 | 37.8503253 | -9.122433971 | 1.86809388 | -4.8832845 | 1.04E-06 | 0.00010221 |
| AMEX60DD026347 | 39.4811536 | -9.182677298 | 1.88971521 | -4.8592916 | 1.18E-06 | 0.00011332 |
| AMEX60DD023676-BARX1 | 410.684217 | -1.839653003 | 0.38388289 | -4.7922245 | 1.65E-06 | 0.00015399 |
| AMEX60DD026060-CSPG5 | 2224.3562 | -0.885631228 | 0.18869574 | -4.6934351 | 2.69E-06 | 0.00023688 |
| AMEX60DD018752-RGS5 | 1035.56562 | -1.383512198 | 0.29521559 | -4.6864469 | 2.78E-06 | 0.00024326 |
| AMEX60DD037204-GRIA3 | 790.004432 | -1.360161105 | 0.29027702 | -4.6857348 | 2.79E-06 | 0.00024326 |
| AMEX60DD016897-CADM3 | 4319.30758 | -0.711943772 | 0.15249352 | -4.6686821 | 3.03E-06 | 0.00026147 |
| AMEX60DD043119-DMRT2 | 1247.36929 | -1.822852985 | 0.39247225 | -4.6445398 | 3.41E-06 | 0.00028927 |
| AMEX60DD003517-ADAMTS17 | 1045.75699 | -1.335682523 | 0.28797684 | -4.6381596 | 3.52E-06 | 0.00029676 |
| AMEX60DD022617-COL6A6 | 17219.8576 | -0.831659369 | 0.18108035 | -4.5927644 | 4.37E-06 | 0.0003597 |
| AMEX60DD054129-DHRS11 | 328.620937 | -1.874988479 | 0.40834482 | -4.5916794 | 4.40E-06 | 0.00035971 |
| AMEX60DD034102-SMPDL3A | 2517.66135 | -0.825621653 | 0.18055393 | -4.572715 | 4.81E-06 | 0.00038787 |
| AMEX60DD006034-COL16A1 | 16251.1998 | -0.647877618 | 0.14254907 | -4.5449446 | 5.49E-06 | 0.00043605 |
| AMEX60DD009964-NGFR | 1543.70786 | -1.094511995 | 0.24171553 | -4.5280996 | 5.95E-06 | 0.00046994 |
| AMEX60DD015640-LOC108800614 | 4669.04 | -0.831837939 | 0.18379678 | -4.5258571 | 6.02E-06 | 0.0004726 |
| AMEX60DD056801-ZNF420 | 32.3832013 | -8.897418932 | 1.96818225 | -4.5206276 | 6.17E-06 | 0.00048205 |
| AMEX60DD048805-EDNRB | 331.205438 | -2.849205465 | 0.6329879 | -4.5012005 | 6.76E-06 | 0.00052059 |
| AMEX60DD002519-ADGRA2 | 5082.33277 | -0.719104604 | 0.16101319 | -4.4661223 | 7.97E-06 | 0.00059911 |
| AMEX60DD017526-CDH13 | 1493.00083 | -1.228331768 | 0.27933656 | -4.3973183 | 1.10E-05 | 0.00079994 |
| AMEX60DD033519-ASRGL1 | 32.2083188 | -8.888896686 | 2.0226045 | -4.3947775 | 1.11E-05 | 0.00079994 |
| AMEX60DD006827-PTHLH | 1151.51399 | -1.209192588 | 0.27910629 | -4.3323732 | 1.48E-05 | 0.00104051 |
| AMEX60DD018528-DDR2 | 981.344926 | -1.160555347 | 0.26913361 | -4.3121903 | 1.62E-05 | 0.00113516 |
| AMEX60DD030355-SLIT3 | 10377.1244 | -0.59325914 | 0.13777085 | -4.3061297 | 1.66E-05 | 0.00116156 |
| AMEX60DD051919-N/A | 3885.96342 | -0.884248561 | 0.20566803 | -4.2993973 | 1.71E-05 | 0.00119216 |
| AMEX60DD019448-JUN | 85018.0985 | -0.777870771 | 0.18096785 | -4.2983921 | 1.72E-05 | 0.00119235 |
| AMEX60DD039130-LAMA1 | 1395.8691 | -1.013952815 | 0.23619376 | -4.2928856 | 1.76E-05 | 0.001217 |
| AMEX60DD011360-HSP90AA1 | 2014.63393 | -0.769427947 | 0.18154458 | -4.2382316 | 2.25E-05 | 0.00153457 |
| AMEX60DD044712-PCDH18 | 1579.45515 | -0.875039803 | 0.20652432 | -4.2369819 | 2.27E-05 | 0.00153653 |
| AMEX60DD003511-SYNM | 2585.07366 | -0.759752302 | 0.17938107 | -4.2354096 | 2.28E-05 | 0.00154074 |
| AMEX60DD045814-QDPR | 5034.3828 | -0.883040666 | 0.20974769 | -4.2100138 | 2.55E-05 | 0.00170283 |

| | | | | | | |
|---|---|---|---|---|---|---|
| AMEX60DD039128-ARHGAP28 | 2557.39667 | -1.013536462 | 0.24173303 | -4.1927926 | 2.76E-05 | 0.00181869 |
| AMEX60DD021967-TRIL | 6508.84744 | -0.635746031 | 0.15164689 | -4.1922788 | 2.76E-05 | 0.00181869 |
| AMEX60DD029989-KANSL2 | 224.640898 | -1.89816043 | 0.4566657 | -4.1565645 | 3.23E-05 | 0.00205097 |
| AMEX60DD044974-HHIP | 3556.94877 | -0.629027157 | 0.15208636 | -4.1359868 | 3.53E-05 | 0.00222592 |
| AMEX60DD052774-ENTPD1 | 1480.50416 | -1.313277764 | 0.32042521 | -4.098547 | 4.16E-05 | 0.00254764 |
| AMEX60DD014782-SEMA6D | 3359.55838 | -0.687062715 | 0.16783829 | -4.0935993 | 4.25E-05 | 0.00259264 |
| AMEX60DD021181-LFNG | 1989.70677 | -0.892786002 | 0.21882736 | -4.0798646 | 4.51E-05 | 0.00274015 |
| AMEX60DD016957-TAT | 4247.50604 | -0.668534429 | 0.16496624 | -4.052553 | 5.07E-05 | 0.00302275 |
| AMEX60DD020537-PTH1R | 699.065941 | -1.217992768 | 0.30098825 | -4.0466455 | 5.20E-05 | 0.00306982 |
| AMEX60DD028631-EMILIN3 | 2851.59946 | -1.053869784 | 0.26045192 | -4.0463122 | 5.20E-05 | 0.00306982 |
| AMEX60DD029855-GPD1 | 1180.26165 | -0.929474987 | 0.23064544 | -4.0298866 | 5.58E-05 | 0.00324418 |
| AMEX60DD052157-PLAU | 5025.12539 | -0.859293262 | 0.2143311 | -4.0091861 | 6.09E-05 | 0.00352918 |
| AMEX60DD034707-RSPO3 | 713.414507 | -1.822443452 | 0.45636827 | -3.9933614 | 6.51E-05 | 0.00371902 |
| AMEX60DD051456-GPR153 | 723.04829 | -1.116441745 | 0.28000231 | -3.9872591 | 6.68E-05 | 0.00380227 |
| AMEX60DD044651-NPNT | 3918.23697 | -0.856487101 | 0.21568147 | -3.9710741 | 7.15E-05 | 0.00404114 |
| AMEX60DD004316-SYT12 | 251.935609 | -1.889732494 | 0.47708122 | -3.9610289 | 7.46E-05 | 0.00420005 |
| AMEX60DD015721-LOC112544205 | 905.429401 | -1.322652487 | 0.336464 | -3.9310372 | 8.46E-05 | 0.00474337 |
| AMEX60DD050932-C8G | 239.582315 | -1.808472556 | 0.46310505 | -3.9051022 | 9.42E-05 | 0.00524498 |
| AMEX60DD014684-ANGPTL4 | 1202.50933 | -1.162037443 | 0.2982507 | -3.8961767 | 9.77E-05 | 0.00537379 |
| AMEX60DD055987-GPR39 | 28.336892 | -7.733423979 | 1.9919098 | -3.8824168 | 0.00010342 | 0.00562135 |
| AMEX60DD007682-RASSF9 | 273.834058 | -1.960485902 | 0.50516385 | -3.8808911 | 0.00010407 | 0.00563742 |
| AMEX60DD039042-TMEFF1 | 4427.68491 | -0.630547781 | 0.16264763 | -3.8767721 | 0.00010585 | 0.00569481 |
| AMEX60DD040413-HAS2 | 4701.91051 | -1.049252588 | 0.2707715 | -3.8750481 | 0.0001066 | 0.0057159 |
| AMEX60DD039671-SNAI2 | 10120.1447 | -0.603165745 | 0.15592335 | -3.8683478 | 0.00010958 | 0.0058358 |
| AMEX60DD004221-MEGF11 | 184.682903 | -2.813602036 | 0.730291 | -3.8527136 | 0.00011682 | 0.00611877 |
| AMEX60DD014711-LPL | 4363.87236 | -0.660440841 | 0.17246571 | -3.8294038 | 0.00012845 | 0.00661914 |
| AMEX60DD018185 | 27.1049154 | -8.6410721 | 2.26469861 | -3.8155506 | 0.00013588 | 0.0069421 |
| AMEX60DD050513-PBX3 | 432.094299 | -1.390330937 | 0.36443131 | -3.81507 | 0.00013614 | 0.0069421 |
| AMEX60DDU001000022-UGCG | 807.980095 | -0.998202089 | 0.26275435 | -3.7989936 | 0.00014529 | 0.00732005 |
| AMEX60DD047799-CRACDL | 482.1324 | -1.487876639 | 0.39209386 | -3.7946951 | 0.00014783 | 0.00742447 |
| AMEX60DD045563-RASL11B | 4764.80976 | -0.805750598 | 0.21293248 | -3.7840662 | 0.00015429 | 0.00772456 |
| AMEX60DD023996-LOC108704014 | 207.43385 | -2.142533559 | 0.56777423 | -3.7735661 | 0.00016093 | 0.00803185 |
| AMEX60DD051354-MMEL1 | 1191.7879 | -0.865166156 | 0.23021363 | -3.7581014 | 0.00017121 | 0.00851798 |
| AMEX60DD039021-GABBR2 | 715.811348 | -1.556425631 | 0.41719608 | -3.7306813 | 0.00019096 | 0.00935422 |
| AMEX60DD002821-GFRA2 | 1891.9653 | -0.697374597 | 0.18707811 | -3.7277188 | 0.00019322 | 0.00940677 |
| AMEX60DD041795-TCF4 | 6743.41714 | -0.722564022 | 0.19416941 | -3.7213073 | 0.00019819 | 0.00961939 |
| AMEX60DD053102-ADRA2A | 180.85026 | -2.56100687 | 0.6890752 | -3.7165855 | 0.00020193 | 0.00973257 |
| AMEX60DD034416-SGK1 | 4322.89952 | -0.775530841 | 0.20876617 | -3.7148301 | 0.00020334 | 0.00973257 |

**Extended Data Table 4. Top Anterior Genes ordered by decreasing fold change below posterior.**

| Gene ID and Symbol | baseMean | log2FoldChange | lfcSE | stat | pvalue | padj |
|---|---|---|---|---|---|---|
| AMEX60DD018728-LHX9 | 310.656084 | -12.15935289 | 1.2850391 | -9.4622435 | 3.01E-21 | 3.99E-18 |
| AMEX60DD048418-LOC115076842 | 158.065401 | -11.18475886 | 1.48058405 | -7.5542883 | 4.21E-14 | 1.52E-11 |
| AMEX60DD029496-HOXC10 | 193.701276 | -10.51461044 | 1.32334737 | -7.9454652 | 1.93E-15 | 8.53E-13 |
| AMEX60DD042420-FOXD1 | 65.8278902 | -9.920178428 | 1.69602903 | -5.8490617 | 4.94E-09 | 9.13E-07 |
| AMEX60DD050768-KRT18 | 48.5623484 | -9.481845879 | 1.72734717 | -5.4892531 | 4.04E-08 | 5.72E-06 |
| AMEX60DD015688-LRRC71 | 43.6647423 | -9.328383525 | 1.77900959 | -5.2435825 | 1.57E-07 | 1.94E-05 |
| AMEX60DD003916-SEMA7A | 39.9199504 | -9.199190674 | 1.8333513 | -5.0176912 | 5.23E-07 | 5.57E-05 |
| AMEX60DD026347 | 39.4811536 | -9.182677298 | 1.88971521 | -4.8592916 | 1.18E-06 | 0.00011332 |
| AMEX60DD007057-LOC101478214 | 37.8503253 | -9.122433971 | 1.86809388 | -4.8832845 | 1.04E-06 | 0.00010221 |
| AMEX60DD056801-ZNF420 | 32.3832013 | -8.897418932 | 1.96818225 | -4.5206276 | 6.17E-06 | 0.00048205 |
| AMEX60DD033519-ASRGL1 | 32.2083188 | -8.888896686 | 2.0226045 | -4.3947775 | 1.11E-05 | 0.00079994 |
| AMEX60DD018185 | 27.1049154 | -8.6410721 | 2.26469861 | -3.8155506 | 0.00013588 | 0.0069421 |
| AMEX60DD055987-GPR39 | 28.336892 | -7.733423979 | 1.9919098 | -3.8824168 | 0.00010342 | 0.00562135 |
| AMEX60DD007691-ALX1 | 1951.28752 | -3.879454882 | 0.40006001 | -9.6971823 | 3.10E-22 | 4.47E-19 |
| AMEX60DD018240-PAPPA2 | 532.556007 | -3.694423657 | 0.6252155 | -5.9090404 | 3.44E-09 | 6.58E-07 |
| AMEX60DD000424-E1301-TTI003913 | 142.185063 | -3.420087065 | 0.63052619 | -5.4241792 | 5.82E-08 | 7.76E-06 |
| AMEX60DD007681-NTS | 400.055312 | -3.250143164 | 0.65830161 | -4.9371642 | 7.93E-07 | 7.96E-05 |
| AMEX60DD029497-HOXC6 | 304.845403 | -3.133002508 | 0.5607923 | -5.5867431 | 2.31E-08 | 3.50E-06 |
| AMEX60DD045837-C1QTNF7 | 2086.61487 | -3.058125714 | 0.26442664 | -11.56512 | 6.19E-31 | 1.40E-27 |
| AMEX60DD048805-EDNRB | 331.205438 | -2.849205465 | 0.6329879 | -4.5012005 | 6.76E-06 | 0.00052059 |
| AMEX60DD017769-SLC6A2 | 1062.99576 | -2.838572615 | 0.33145193 | -8.5640553 | 1.09E-17 | 7.86E-15 |
| AMEX60DD004221-MEGF11 | 184.682903 | -2.813602036 | 0.730291 | -3.8527136 | 0.00011682 | 0.00611877 |
| AMEX60DD053102-ADRA2A | 180.85026 | -2.56100687 | 0.6890752 | -3.7165855 | 0.00020193 | 0.00973257 |
| AMEX60DD040394-ENPP2 | 4769.4996 | -2.507817369 | 0.19723487 | -12.714879 | 4.89E-37 | 1.94E-33 |
| AMEX60DD013358-PER1 | 389.257763 | -2.233125923 | 0.36879543 | -6.0551888 | 1.40E-09 | 2.85E-07 |
| AMEX60DD019770-LOC115099908 | 1390.74803 | -2.210020809 | 0.25941041 | -8.5193991 | 1.60E-17 | 1.11E-14 |
| AMEX60DD050555-LHX2 | 763.615828 | -2.165270948 | 0.30083957 | -7.1974273 | 6.14E-13 | 1.84E-10 |
| AMEX60DD023996-LOC108704014 | 207.43385 | -2.142533559 | 0.56777423 | -3.7735661 | 0.00016093 | 0.00803185 |
| AMEX60DD036390-MDGA1 | 1168.69929 | -2.027373767 | 0.26660386 | -7.6044426 | 2.86E-14 | 1.08E-11 |
| AMEX60DD000888-MN1 | 761.370421 | -2.026388821 | 0.27135362 | -7.4677052 | 8.16E-14 | 2.82E-11 |
| AMEX60DD007682-RASSF9 | 273.834058 | -1.960485902 | 0.50516385 | -3.8808911 | 0.00010407 | 0.00563742 |
| AMEX60DD037579-TBX22 | 981.691962 | -1.940600339 | 0.26462578 | -7.3333759 | 2.24E-13 | 6.85E-11 |
| AMEX60DD034577-SMOC2 | 4361.98956 | -1.922237066 | 0.16503035 | -11.647779 | 2.36E-31 | 6.23E-28 |
| AMEX60DD029989-KANSL2 | 224.640898 | -1.89816043 | 0.4566657 | -4.1565645 | 3.23E-05 | 0.00205097 |
| AMEX60DD004316-SYT12 | 251.935609 | -1.889732494 | 0.47708122 | -3.9610289 | 7.46E-05 | 0.00420005 |
| AMEX60DD028693-VSTM2L | 2602.59124 | -1.884959595 | 0.30977347 | -6.0849614 | 1.17E-09 | 2.43E-07 |
| AMEX60DD054129-DHRS11 | 328.620937 | -1.874988479 | 0.40834482 | -4.5916794 | 4.40E-06 | 0.00035971 |
| AMEX60DD023676-BARX1 | 410.684217 | -1.839653003 | 0.38388289 | -4.7922245 | 1.65E-06 | 0.00015399 |
| AMEX60DD046302-EGR4 | 488.999052 | -1.833383622 | 0.37090205 | -4.9430399 | 7.69E-07 | 7.78E-05 |
| AMEX60DD043119-DMRT2 | 1247.36929 | -1.822852985 | 0.39247225 | -4.6445398 | 3.41E-06 | 0.00028927 |
| AMEX60DD002138-EPHA4 | 1559.31708 | -1.822645064 | 0.20313111 | -8.972752 | 2.89E-19 | 2.55E-16 |
| AMEX60DD034707-RSPO3 | 713.414507 | -1.822443452 | 0.45636827 | -3.9933614 | 6.51E-05 | 0.00371902 |
| AMEX60DD043069-DAPK1 | 1267.48303 | -1.814804171 | 0.35242194 | -5.1495209 | 2.61E-07 | 3.05E-05 |
| AMEX60DD050932-C8G | 239.582315 | -1.808472556 | 0.46310505 | -3.9051022 | 9.42E-05 | 0.00524498 |
| AMEX60DD033076-GDF6 | 451.802642 | -1.690135526 | 0.33233548 | -5.0856308 | 3.66E-07 | 4.12E-05 |
| AMEX60DD042580-NR2F1 | 866.38496 | -1.615003099 | 0.27963255 | -5.7754475 | 7.67E-09 | 1.30E-06 |
| AMEX60DD003526-ALDH1A3 | 8943.15973 | -1.604185396 | 0.17837179 | -8.993493 | 2.39E-19 | 2.24E-16 |
| AMEX60DD018273-PRDX6 | 34403.2575 | -1.573974711 | 0.17982707 | -8.7527127 | 2.08E-18 | 1.57E-15 |
| AMEX60DD021193-CHST12 | 644.718904 | -1.569407653 | 0.29570127 | -5.3074092 | 1.11E-07 | 1.43E-05 |
| AMEX60DD039021-GABBR2 | 715.811348 | -1.556425631 | 0.41719608 | -3.7306813 | 0.00019096 | 0.00935422 |
| AMEX60DD047799-CRACDL | 482.1324 | -1.487876639 | 0.39209386 | -3.7946951 | 0.00014783 | 0.00742447 |
| AMEX60DD005869-RSPO1 | 9175.68926 | -1.487338743 | 0.18557085 | -8.0149375 | 1.10E-15 | 5.00E-13 |
| AMEX60DD034565-THBS2 | 27875.7261 | -1.471111044 | 0.12247635 | -12.011388 | 3.10E-33 | 9.83E-30 |

| | | | | | | |
|---|---|---|---|---|---|---|
| AMEX60DD050513-PBX3 | 432.094299 | -1.390330937 | 0.36443131 | -3.81507 | 0.00013614 | 0.0069421 |
| AMEX60DD018752-RGS5 | 1035.56562 | -1.383512198 | 0.29521559 | -4.6864469 | 2.78E-06 | 0.00024326 |
| AMEX60DD048332-COL8A1 | 2203.62174 | -1.379899396 | 0.27529397 | -5.0124578 | 5.37E-07 | 5.69E-05 |
| AMEX60DD037204-GRIA3 | 790.004432 | -1.360161105 | 0.29027702 | -4.6857348 | 2.79E-06 | 0.00024326 |
| AMEX60DD003517-ADAMTS17 | 1045.75699 | -1.335682523 | 0.28797684 | -4.6381596 | 3.52E-06 | 0.00029676 |
| AMEX60DD019011-CCN1 | 38850.2766 | -1.33115672 | 0.20137848 | -6.6102232 | 3.84E-11 | 9.09E-09 |
| AMEX60DD007033-FAM180A | 2308.91906 | -1.322991273 | 0.22096547 | -5.9873215 | 2.13E-09 | 4.29E-07 |
| AMEX60DD015721-LOC112544205 | 905.429401 | -1.322652487 | 0.336464 | -3.9310372 | 8.46E-05 | 0.00474337 |
| AMEX60DD013889-OCLN | 3555.05707 | -1.315658782 | 0.24180464 | -5.440999 | 5.30E-08 | 7.31E-06 |
| AMEX60DD019499-PLPP3 | 18098.7828 | -1.31367337 | 0.14406571 | -9.1185707 | 7.61E-20 | 7.55E-17 |
| AMEX60DD052774-ENTPD1 | 1480.50416 | -1.313277764 | 0.32042521 | -4.098547 | 4.16E-05 | 0.00254764 |
| AMEX60DD036008-PKDCC | 2041.34381 | -1.283665815 | 0.18735552 | -6.8514973 | 7.31E-12 | 1.87E-09 |
| AMEX60DD001004-EMID1 | 7308.09216 | -1.264861261 | 0.19686039 | -6.4251689 | 1.32E-10 | 3.03E-08 |
| AMEX60DD028577 | 1321.32344 | -1.262353305 | 0.2260491 | -5.5844209 | 2.34E-08 | 3.51E-06 |
| AMEX60DD001838-PTX3 | 50987.6973 | -1.248917624 | 0.17394136 | -7.1801074 | 6.97E-13 | 2.05E-10 |
| AMEX60DD045225-CPE | 940.961677 | -1.238757914 | 0.24386402 | -5.0797075 | 3.78E-07 | 4.20E-05 |
| AMEX60DD014035-CPAMD8 | 15019.7225 | -1.237477495 | 0.11861955 | -10.432324 | 1.77E-25 | 2.80E-22 |
| AMEX60DD014152-CRLF1 | 4087.29505 | -1.237293188 | 0.1332404 | -9.2861712 | 1.60E-20 | 1.81E-17 |
| AMEX60DD017526-CDH13 | 1493.00083 | -1.228331768 | 0.27933656 | -4.3973183 | 1.10E-05 | 0.00079994 |
| AMEX60DD020537-PTH1R | 699.065941 | -1.217992768 | 0.30098825 | -4.0466455 | 5.20E-05 | 0.00306982 |
| AMEX60DD030673-NTN1 | 4615.27492 | -1.214084683 | 0.1635444 | -7.4235786 | 1.14E-13 | 3.82E-11 |
| AMEX60DD006827-PTHLH | 1151.51399 | -1.209192588 | 0.27910629 | -4.3323732 | 1.48E-05 | 0.00104051 |
| AMEX60DD027622-R3HDML | 1110.89122 | -1.206491732 | 0.2284927 | -5.28022 | 1.29E-07 | 1.63E-05 |
| AMEX60DD018972-AK5 | 4221.21616 | -1.184540473 | 0.24242398 | -4.8862348 | 1.03E-06 | 0.00010132 |
| AMEX60DD051000-LMO4 | 5762.20934 | -1.171535729 | 0.22139688 | -5.2915638 | 1.21E-07 | 1.55E-05 |
| AMEX60DD014684-ANGPTL4 | 1202.50933 | -1.162037443 | 0.2982507 | -3.8961767 | 9.77E-05 | 0.00537379 |
| AMEX60DD018528-DDR2 | 981.344926 | -1.160555347 | 0.26913361 | -4.3121903 | 1.62E-05 | 0.00113516 |
| AMEX60DD003787-THSD4 | 4528.18517 | -1.156400564 | 0.19913241 | -5.807194 | 6.35E-09 | 1.13E-06 |
| AMEX60DD052816-CRTAC1 | 4611.44867 | -1.15509378 | 0.16171993 | -7.1425568 | 9.16E-13 | 2.64E-10 |
| AMEX60DD035184-SRD5A2 | 5667.73939 | -1.143950113 | 0.19300763 | -5.9269685 | 3.09E-09 | 5.97E-07 |
| AMEX60DD016542-NTN1 | 9019.96556 | -1.143902186 | 0.2300058 | -4.9733624 | 6.58E-07 | 6.74E-05 |
| AMEX60DD051456-GPR153 | 723.04829 | -1.116441745 | 0.28000231 | -3.9872591 | 6.68E-05 | 0.00380227 |
| AMEX60DD033627-COL12A1 | 90200.8947 | -1.114222582 | 0.16412379 | -6.7889159 | 1.13E-11 | 2.80E-09 |
| AMEX60DD015718-CRABP2 | 16671.4165 | -1.104369399 | 0.145927 | -7.5679578 | 3.79E-14 | 1.40E-11 |
| AMEX60DD009964-NGFR | 1543.70786 | -1.094511995 | 0.24171553 | -4.5280996 | 5.95E-06 | 0.00046994 |
| AMEX60DD046939-RABL3 | 2967.96611 | -1.087722657 | 0.15241696 | -7.1364934 | 9.57E-13 | 2.67E-10 |
| AMEX60DD028631-EMILIN3 | 2851.59946 | -1.053869784 | 0.26045192 | -4.0463122 | 5.20E-05 | 0.00306982 |
| AMEX60DD040413-HAS2 | 4701.91051 | -1.049252588 | 0.2707715 | -3.8750481 | 0.0001066 | 0.0057159 |
| AMEX60DD043399-BNC2 | 2331.18423 | -1.044435736 | 0.19505282 | -5.3546304 | 8.57E-08 | 1.12E-05 |
| AMEX60DD002008-PID1 | 3766.15586 | -1.03417846 | 0.17899906 | -5.7775636 | 7.58E-09 | 1.29E-06 |
| AMEX60DD012076-PAPLN | 2140.7147 | -1.015133212 | 0.17516707 | -5.7952286 | 6.82E-09 | 1.19E-06 |
| AMEX60DD039130-LAMA1 | 1395.8691 | -1.013952815 | 0.23619376 | -4.2928856 | 1.76E-05 | 0.001217 |
| AMEX60DD039128-ARHGAP28 | 2557.39667 | -1.013536462 | 0.24173303 | -4.1927926 | 2.76E-05 | 0.00181869 |
| AMEX60DDU001000022-UGCG | 807.980095 | -0.998202089 | 0.26275435 | -3.7989936 | 0.00014529 | 0.00732005 |
| AMEX60DD035020-SLC16A10 | 1933.92486 | -0.987392117 | 0.1789146 | -5.51879 | 3.41E-08 | 4.92E-06 |
| AMEX60DD029025-CLK4 | 3622.54533 | -0.971096464 | 0.15803843 | -6.1446855 | 8.01E-10 | 1.70E-07 |
| AMEX60DD029855-GPD1 | 1180.26165 | -0.929474987 | 0.23064544 | -4.0298866 | 5.58E-05 | 0.00324418 |
| AMEX60DD017214-CRISPLD2 | 5668.74564 | -0.926187592 | 0.14267374 | -6.4916472 | 8.49E-11 | 1.98E-08 |
| AMEX60DD001527-CHRD | 5748.75414 | -0.918562679 | 0.14438121 | -6.3620652 | 1.99E-10 | 4.45E-08 |
| AMEX60DD021181-LFNG | 1989.70677 | -0.892786002 | 0.21882736 | -4.0798646 | 4.51E-05 | 0.00274015 |
| AMEX60DD026060-CSPG5 | 2224.3562 | -0.885631228 | 0.18869574 | -4.6934351 | 2.69E-06 | 0.00023688 |
| AMEX60DD051919-N/A | 3885.96342 | -0.884248561 | 0.20566803 | -4.2993973 | 1.71E-05 | 0.00119216 |
| AMEX60DD045814-QDPR | 5034.3828 | -0.883040666 | 0.20974769 | -4.2100138 | 2.55E-05 | 0.00170283 |
| AMEX60DD043210-PCSK5 | 4999.43042 | -0.881173123 | 0.15183104 | -5.8036428 | 6.49E-09 | 1.14E-06 |
| AMEX60DD044712-PCDH18 | 1579.45515 | -0.875039803 | 0.20652432 | -4.2369819 | 2.27E-05 | 0.00153653 |
| AMEX60DD051354-MMEL1 | 1191.7879 | -0.865166156 | 0.23021363 | -3.7581014 | 0.00017121 | 0.00851798 |

| | | | | | | |
|---|---|---|---|---|---|---|
| AMEX60DD048505-EGFL6 | 8627.65393 | -0.864325484 | 0.12270735 | -7.0437955 | 1.87E-12 | 5.03E-10 |
| AMEX60DD052157-PLAU | 5025.12539 | -0.859293262 | 0.2143311 | -4.0091861 | 6.09E-05 | 0.00352918 |
| AMEX60DD044651-NPNT | 3918.23697 | -0.856487101 | 0.21568147 | -3.9710741 | 7.15E-05 | 0.00404114 |
| AMEX60DD045587-CORIN | 3252.86717 | -0.851020218 | 0.17393292 | -4.8928071 | 9.94E-07 | 9.86E-05 |
| AMEX60DD053448-CDON | 10079.3829 | -0.848501068 | 0.10704207 | -7.9267996 | 2.25E-15 | 9.65E-13 |
| AMEX60DD015640-LOC108800614 | 4669.04 | -0.831837939 | 0.18379678 | -4.5258571 | 6.02E-06 | 0.0004726 |
| AMEX60DD022617-COL6A6 | 17219.8576 | -0.831659369 | 0.18108035 | -4.5927644 | 4.37E-06 | 0.0003597 |
| AMEX60DD022752-ZNF385D | 2930.91358 | -0.828870742 | 0.16666848 | -4.9731703 | 6.59E-07 | 6.74E-05 |
| AMEX60DD049175-POSTN | 328298.719 | -0.828029956 | 0.10267285 | -8.0647409 | 7.34E-16 | 3.43E-13 |
| AMEX60DD034102-SMPDL3A | 2517.66135 | -0.825621653 | 0.18055393 | -4.572715 | 4.81E-06 | 0.00038787 |
| AMEX60DD046940-HGD | 13281.7243 | -0.821919948 | 0.14089492 | -5.833567 | 5.43E-09 | 9.90E-07 |
| AMEX60DD045563-RASL11B | 4764.80976 | -0.805750598 | 0.21293248 | -3.7840662 | 0.00015429 | 0.00772456 |
| AMEX60DD017420-CD82 | 3323.35702 | -0.804100117 | 0.14682791 | -5.4764802 | 4.34E-08 | 6.09E-06 |
| AMEX60DD048164-ITGBL1 | 17043.675 | -0.800340444 | 0.14372803 | -5.5684369 | 2.57E-08 | 3.78E-06 |
| AMEX60DD024053-FBLN2 | 9726.5494 | -0.793794924 | 0.13937634 | -5.6953349 | 1.23E-08 | 2.01E-06 |
| AMEX60DD035799-ISM1 | 9466.91442 | -0.779163331 | 0.1217125 | -6.4016706 | 1.54E-10 | 3.48E-08 |
| AMEX60DD019448-JUN | 85018.0985 | -0.777870771 | 0.18096785 | -4.2983921 | 1.72E-05 | 0.00119235 |
| AMEX60DD034416-SGK1 | 4322.89952 | -0.775530841 | 0.20876617 | -3.7148301 | 0.00020334 | 0.00973257 |
| AMEX60DD011360-HSP90AA1 | 2014.63393 | -0.769427947 | 0.18154458 | -4.2382316 | 2.25E-05 | 0.00153457 |
| AMEX60DD003511-SYNM | 2585.07366 | -0.759752302 | 0.17938107 | -4.2354096 | 2.28E-05 | 0.00154074 |
| AMEX60DD041795-TCF4 | 6743.41714 | -0.722564022 | 0.19416941 | -3.7213073 | 0.00019819 | 0.00961939 |
| AMEX60DD002519-ADGRA2 | 5082.33277 | -0.719104604 | 0.16101319 | -4.4661223 | 7.97E-06 | 0.00059911 |
| AMEX60DDU001005418-SEMA3F | 6149.82378 | -0.712649484 | 0.12687116 | -5.6171116 | 1.94E-08 | 3.02E-06 |
| AMEX60DD016897-CADM3 | 4319.30758 | -0.711943772 | 0.15249352 | -4.6686821 | 3.03E-06 | 0.00026147 |
| AMEX60DD002821-GFRA2 | 1891.9653 | -0.697374597 | 0.18707811 | -3.7277188 | 0.00019322 | 0.00940677 |
| AMEX60DD024186-GNAI2 | 5249.54012 | -0.689900313 | 0.13811002 | -4.9952953 | 5.87E-07 | 6.17E-05 |
| AMEX60DD014782-SEMA6D | 3359.55838 | -0.687062715 | 0.16783829 | -4.0935993 | 4.25E-05 | 0.00259264 |
| AMEX60DD016957-TAT | 4247.50604 | -0.668534429 | 0.16496624 | -4.052553 | 5.07E-05 | 0.00302275 |
| AMEX60DD014711-LPL | 4363.87236 | -0.660440841 | 0.17246571 | -3.8294038 | 0.00012845 | 0.00661914 |
| AMEX60DD006034-COL16A1 | 16251.1998 | -0.647877618 | 0.14254907 | -4.5449446 | 5.49E-06 | 0.00043605 |
| AMEX60DD021967-TRIL | 6508.84744 | -0.635746031 | 0.15164689 | -4.1922788 | 2.76E-05 | 0.00181869 |
| AMEX60DD039042-TMEFF1 | 4427.68491 | -0.630547781 | 0.16264763 | -3.8767721 | 0.00010585 | 0.00569481 |
| AMEX60DD044974-HHIP | 3556.94877 | -0.629027157 | 0.15208636 | -4.1359868 | 3.53E-05 | 0.00222592 |
| AMEX60DD039671-SNAI2 | 10120.1447 | -0.603165745 | 0.15592335 | -3.8683478 | 0.00010958 | 0.0058358 |
| AMEX60DD028164-MFAP4 | 23910.4664 | -0.59668289 | 0.12008112 | -4.9689984 | 6.73E-07 | 6.85E-05 |
| AMEX60DD030355-SLIT3 | 10377.1244 | -0.59325914 | 0.13777085 | -4.3061297 | 1.66E-05 | 0.00116156 |
| AMEX60DD039942-ZFHX4 | 6203.76701 | -0.590178723 | 0.11607079 | -5.0846445 | 3.68E-07 | 4.12E-05 |

**Extended Data Table 5. Significant genes associated with GO term "Extracellular matrix".**

| Gene ID and Symbol | baseMean | log2FoldChange | lfcSE | stat | pvalue | padj |
|---|---|---|---|---|---|---|
| ADAMTS15 | 49.7214696 | 6.279766739 | 1.48093962 | 4.24039349 | 2.23E-05 | 0.00152641 |
| ADAMTS17 | 1045.75699 | -1.335682523 | 0.28797684 | -4.6381596 | 3.52E-06 | 0.00029676 |
| ADAMTS5 | 1426.64432 | 1.72120583 | 0.29923357 | 5.75204791 | 8.82E-09 | 1.47E-06 |
| EGFL6 | 8627.65393 | -0.864325484 | 0.12270735 | -7.0437955 | 1.87E-12 | 5.03E-10 |
| SMOC2 | 4361.98956 | -1.922237066 | 0.16503035 | -11.647779 | 2.36E-31 | 6.23E-28 |
| CCN1 | 38850.2766 | -1.33115672 | 0.20137848 | -6.6102232 | 3.84E-11 | 9.09E-09 |
| COL3A1 | 1421960.7 | 0.499407033 | 0.09620593 | 5.19102127 | 2.09E-07 | 2.50E-05 |
| COL6A6 | 17219.8576 | -0.831659369 | 0.18108035 | -4.5927644 | 4.37E-06 | 0.0003597 |
| COL8A1 | 2203.62174 | -1.379899396 | 0.27529397 | -5.0124578 | 5.37E-07 | 5.69E-05 |
| COL11A1 | 326.320566 | 2.5619916 | 0.54737983 | 4.68046401 | 2.86E-06 | 0.00024824 |
| COL16A1 | 16251.1998 | -0.647877618 | 0.14254907 | -4.5449446 | 5.49E-06 | 0.00043605 |
| COL17A1 | 7154.99358 | 1.425025615 | 0.16854798 | 8.45471779 | 2.80E-17 | 1.85E-14 |
| CRISPLD2 | 5668.74564 | -0.926187592 | 0.14267374 | -6.4916472 | 8.49E-11 | 1.98E-08 |
| FBLN2 | 9726.5494 | -0.793794924 | 0.13937634 | -5.6953349 | 1.23E-08 | 2.01E-06 |
| MMP1 | 48.8004415 | 8.667396854 | 2.06802382 | 4.19114944 | 2.78E-05 | 0.00182021 |
| NPNT | 3918.23697 | -0.856487101 | 0.21568147 | -3.9710741 | 7.15E-05 | 0.00404114 |
| PAPLN | 2140.7147 | -1.015133212 | 0.17516707 | -5.7952286 | 6.82E-09 | 1.19E-06 |
| PTX3 | 50987.6973 | -1.248917624 | 0.17394136 | -7.1801074 | 6.97E-13 | 2.05E-10 |
| POSTN | 328298.719 | -0.828029956 | 0.10267285 | -8.0647409 | 7.34E-16 | 3.43E-13 |
| THSD4 | 4528.18517 | -1.156400564 | 0.19913241 | -5.807194 | 6.35E-09 | 1.13E-06 |

**Extended Data Table 6. Significant genes associated with GO term "Cell Adhesion".**

| Gene ID and Symbol | baseMean | log2FoldChange | lfcSE | stat | pvalue | padj |
|---|---|---|---|---|---|---|
| CD34 | 1568.35589 | 1.032026831 | 0.22443686 | 4.59829481 | 4.26E-06 | 0.00035211 |
| EGFL6 | 8627.65393 | -0.864325484 | 0.12270735 | -7.0437955 | 1.87E-12 | 5.03E-10 |
| EPHA4 | 1559.31708 | -1.822645064 | 0.20313111 | -8.972752 | 2.89E-19 | 2.55E-16 |
| AJAP1 | 1082.95382 | 1.098173741 | 0.22843592 | 4.80736022 | 1.53E-06 | 0.00014448 |
| ACAN | 2490.26505 | 0.994854433 | 0.23882297 | 4.1656564 | 3.10E-05 | 0.00200296 |
| CDON | 10079.3829 | -0.848501068 | 0.10704207 | -7.9267996 | 2.25E-15 | 9.65E-13 |
| CCN1 | 38850.2766 | -1.33115672 | 0.20137848 | -6.6102232 | 3.84E-11 | 9.09E-09 |
| COL6A3 | 211560.903 | -0.475382885 | 0.12410385 | -3.8305249 | 0.00012787 | 0.00661052 |
| COL6A6 | 17219.8576 | -0.831659369 | 0.18108035 | -4.5927644 | 4.37E-06 | 0.0003597 |
| COL8A1 | 2203.62174 | -1.379899396 | 0.27529397 | -5.0124578 | 5.37E-07 | 5.69E-05 |
| COL12A1 | 90200.8947 | -1.114222582 | 0.16412379 | -6.7889159 | 1.13E-11 | 2.80E-09 |
| COL16A1 | 16251.1998 | -0.647877618 | 0.14254907 | -4.5449446 | 5.49E-06 | 0.00043605 |
| DDR2 | 981.344926 | -1.160555347 | 0.26913361 | -4.3121903 | 1.62E-05 | 0.00113516 |
| ENTPD1 | 1480.50416 | -1.313277764 | 0.32042521 | -4.098547 | 4.16E-05 | 0.00254764 |
| FN1 | 57753.2051 | 0.75250987 | 0.09000369 | 8.36087779 | 6.23E-17 | 3.66E-14 |
| ISLR | 3994.21959 | 0.772821134 | 0.1623958 | 4.75887383 | 1.95E-06 | 0.00017913 |
| ITGBL1 | 17043.675 | -0.800340444 | 0.14372803 | -5.5684369 | 2.57E-08 | 3.78E-06 |
| ICAM5 | 976.15079 | 1.352357644 | 0.30739378 | 4.39943072 | 1.09E-05 | 0.00079748 |
| LAMA1 | 1395.8691 | -1.013952815 | 0.23619376 | -4.2928856 | 1.76E-05 | 0.001217 |
| LAMA4 | 6227.24234 | 0.653641319 | 0.12664745 | 5.16110928 | 2.45E-07 | 2.91E-05 |
| LAMB2 | 1063.00385 | 1.22505409 | 0.26074329 | 4.69831489 | 2.62E-06 | 0.00023258 |
| LAMC1 | 16317.0264 | 0.46080403 | 0.102433 | 4.49858957 | 6.84E-06 | 0.00052448 |
| MFAP4 | 23910.4664 | -0.59668289 | 0.12008112 | -4.9689984 | 6.73E-07 | 6.85E-05 |
| NPNT | 3918.23697 | -0.856487101 | 0.21568147 | -3.9710741 | 7.15E-05 | 0.00404114 |
| NCAM1 | 9044.87059 | 0.566152141 | 0.1295067 | 4.37160514 | 1.23E-05 | 0.00087387 |
| POSTN | 328298.719 | -0.828029956 | 0.10267285 | -8.0647409 | 7.34E-16 | 3.43E-13 |
| PCDH18 | 1579.45515 | -0.875039803 | 0.20652432 | -4.2369819 | 2.27E-05 | 0.00153653 |
| PCDH19 | 1854.51964 | 0.808726282 | 0.20248724 | 3.99396176 | 6.50E-05 | 0.00371902 |
| TNXB | 14399.8852 | 1.025301077 | 0.21312368 | 4.81082658 | 1.50E-06 | 0.00014369 |
| THBS2 | 27875.7261 | -1.471111044 | 0.12247635 | -12.011388 | 3.10E-33 | 9.83E-30 |
| THBS3 | 4237.65202 | 1.653942957 | 0.20915441 | 7.90776031 | 2.62E-15 | 1.09E-12 |
| VCAN | 58932.4203 | -0.540519768 | 0.14395394 | -3.7548105 | 0.00017347 | 0.00855026 |
